# Noradrenergic infraslow rhythm during sleep is the critical link between heart-rate dynamics and memory consolidation

**DOI:** 10.64898/2026.01.06.697926

**Authors:** Sofie S. Jacobsen, Allison B. Morehouse, Pin-Chun Chen, Yi Qian, Ryszard S. Gomolka, Mie Andersen, Maiken Nedergaard, Sara C. Mednick, Celia Kjaerby

**Author notes:** Corresponding authors: Celia Kjaerby Sara C. Mednick. Shared last authorship (Sara C. Mednick, Celia Kjaerby).

## Abstract

Recent work shows that the brain’s arousal system remains active during sleep, with rhythmic locus coeruleus (LC) activity shaping sleep architecture and supporting memory consolidation. The LC releases norepinephrine (NE) in infraslow (∼0.02 Hz) bouts that gate NREM sleep spindles. Here, we demonstrate that heart rate (HR) fluctuations during NREM are tightly phase-locked to these NE rhythms, identifying the LC as a key driver of very-low-frequency HR variability (VLF-HRV), an understudied autonomic signal. Using optogenetics, transient LC inhibition blunts HR slowing, whereas LC activation produces rapid HR acceleration, directly linking LC output to cardiac control during sleep. We further show that infraslow HR variability is a cross-species marker of spindle-dependent memory processing. In mice, the amplitude of HR decelerations during NREM correlate with spindle activity and subsequent memory performance. Remarkably, human sleepers show the same pattern: stronger VLF-HR fluctuations during NREM correspond to increased spindle expression and better overnight memory retention. These findings reveal a mechanistic pathway through which LC activity modulates autonomic physiology during sleep and identify infraslow HR variability as a non-invasive marker of brainstem function and memory-promoting sleep. Because LC degeneration occurs early in neurodegenerative disease, sleep-derived HR metrics may provide a scalable indicator of emerging neuromodulatory dysfunction.

## Introduction

Heart rate variability (HRV) is a well-established, noninvasive biomarker of autonomic nervous system function and cardiovascular health. Reduced HRV is associated with increased morbidity and mortality and is frequently observed in neurological disorders such as Alzheimer’s disease, where it may reflect central autonomic network dysfunction (Natarajan et al., 2020; Zulli et al., 2005). Yet, despite its widespread use, decoding of heart rate signals and the central drivers of HRV remain elusive. One potential orchestrator of brain-body communication is the locus coeruleus-norepinephrine (LC-NE) system; LC is a brainstem nucleus that is part of the brain’s arousal systems. A recent discovery of an infraslow phasic pattern (∼30-50 s) of LC-NE activity during non-rapid eye movement (NREM) sleep has linked LC-NE activity with a wide range of critical cortical functions, including memory consolidation as well as glymphatic clearance (Hauglund et al., 2025; Kjaerby et al., 2022; Osorio-Forero et al., 2021). The ascending phase of NE fluctuations is often coupled to brief periods of intrinsic arousals known as micro-arousals (MAs) (Andrillon et al., 2011). In contrast, the declining phase of NE is associated with an increase in sleep spindles, which are discrete oscillatory events lasting 1 to 2 seconds in the sigma band (8–15 Hz) implicated in memory consolidation (Andrillon et al., 2011; Fernandez and Lüthi, 2020; Mednick et al., 2013). The LC-generated infraslow phasic fluctuations of NE lead to a microstructural organization of NREM sleep shifting between arousal and spindle-rich segments (Kjaerby et al., 2022; Osorio-Forero et al., 2021). Additionally, infraslow NE activity is involved in organization of ripple-dependent memory consolidation (Chang et al., 2025) and elicit vasomotion-dependent cerebrospinal flow (Hauglund et al., 2025), indicating a broader functional role of infraslow phasic NE fluctuations in sleep restoration. With age, LC-NE function deteriorates along with restorative NREM sleep and memory performance (Grudzien et al., 2007; Mann et al., 1980). Yet, the LC brain structure and physiological markers are difficult to access in humans. Thus, there is a need to explore more approachable biomarkers for sleep-dependent memory consolidation.

Sleep spindles, along with their role in memory consolidation, also show tight coupling with the peripheral autonomic nervous system (Chen et al., 2020; Naji et al., 2019a). Specifically, phasic bursts in heart rate (HR) during NREM sleep are preceded by surges in delta power (1-4 Hz) and sleep spindles, and this temporal coupling has been linked to memory improvements across sleep (Chen et al., 2020; Naji et al., 2019a). It has previously been reported that HR fluctuates at similar infraslow frequencies as NE fluctuations and sigma power in mice (Lecci et al., 2017; Osorio-Forero et al., 2021) and that phasic HR fluctuations correlate with infraslow changes in pupil diameter (Carro-Domínguez et al., 2025), a proxy for changes in NE levels (Murphy et al., 2014; Reimer et al., 2016). Importantly, optogenetic manipulation of the LC has demonstrated that infraslow LC activity coordinates sleep spindle clustering and heart rate fluctuations during NREM sleep (Osorio-Forero et al., 2021). Together, these findings support a functional coupling between the central LC–NE system and peripheral cardiac dynamics, further supported by findings that HR increases accompany MAs during NREM sleep (Carro-Domínguez et al., 2025; Osorio-Forero et al., 2025).

The sympathetic and parasympathetic autonomic nervous system are involved in HRV, which is conventionally analyzed across three primary frequency bands: high frequency (HF) HRV, low frequency (LF) HRV, and very low frequency (VLF) HRV (Berntson et al., 1997). Due to the frequency overlap with VLF HRV, we wondered if central infraslow NE dynamics could be linked to this poorly understood HRV indicator. Furthermore, are infraslow NE fluctuations directly reflected by HRV under different physiological states or does LC–HR coupling scales differently with LC output? Specifically, if infraslow NE oscillations display faster frequencies - as occurs during sleep fragmentation - will cardiac dynamics exhibit corresponding changes?

Conversely, given that stronger infraslow NE dynamics correlate with memory consolidation through their regulation of sleep spindles, could the peripheral autonomic signatures provide an accessible cross-species biomarker of spindle-dependent memory consolidation? Addressing these questions could help bridge mechanistic insights into LC-mediated sleep regulation with established HRV metrics used in human physiology.

In this study, we combined real-time fluorescent imaging of NE with simultaneous heart rate recordings in mice to test whether phasic infraslow LC–NE activity during NREM sleep is reflected in HRV dynamics. We further examined whether these brain–heart relationships change when LC–NE oscillatory activity is experimentally shifted toward faster frequencies using titrated optogenetic stimulation during sleep. Finally, we investigated whether cardiac dynamics associated with these neuromodulatory rhythms relate to spindle regulation and memory consolidation in both mice and humans.

## Results

### Norepinephrine and heart rate correlate during NREM sleep

Separate areas of research has demonstrated that both NE and HR signals fluctuate every 30-50s. Here, we used causal experimental methods to explore whether these independent infraslow fluctuations may be coupled and how they are impacted by different sleep transitions (Kjaerby et al., 2022; Osorio-Forero et al., 2025, 2021; Tanoue et al., 2022; Watson et al., 1979).

Toward this goal, we expressed the fluorescent NE biosensor GRAB-NE2m(3.1) in the medial prefrontal cortex (mPFC) of adult C57BL/6 mice and recorded extracellular NE using fiber photometry simultaneously with EEG and EMG recordings covering multiple sleep–wake episodes (Fig. 1a). mPFC was selected as a representative cortical readout of LC-mediated NE dynamics, as infraslow NE fluctuations are coordinated across widespread brain regions. R-peaks of the QRS complex of the heart were extracted from the EMG (for details on HR/RR extraction, see the Method section ‘Heart rate detection’ and Supplementary Fig. 1a-c) and the inter-beat-interval (RR) was calculated (Sörnmo and Laguna, 2005) (Supplementary Fig. 1d-e). A shorter RR interval reflects faster HR and vice-versa. HR during the NREM period was on average 493 bpm (SD ± 59) while REM HR was 519 bpm (SD ± 67) both of which was consistent with the expected 450-600 bpm, based on the literature (D’Souza et al., 2021; Ho et al., 2011; Sakata et al., 2005).

**Figure 1:**
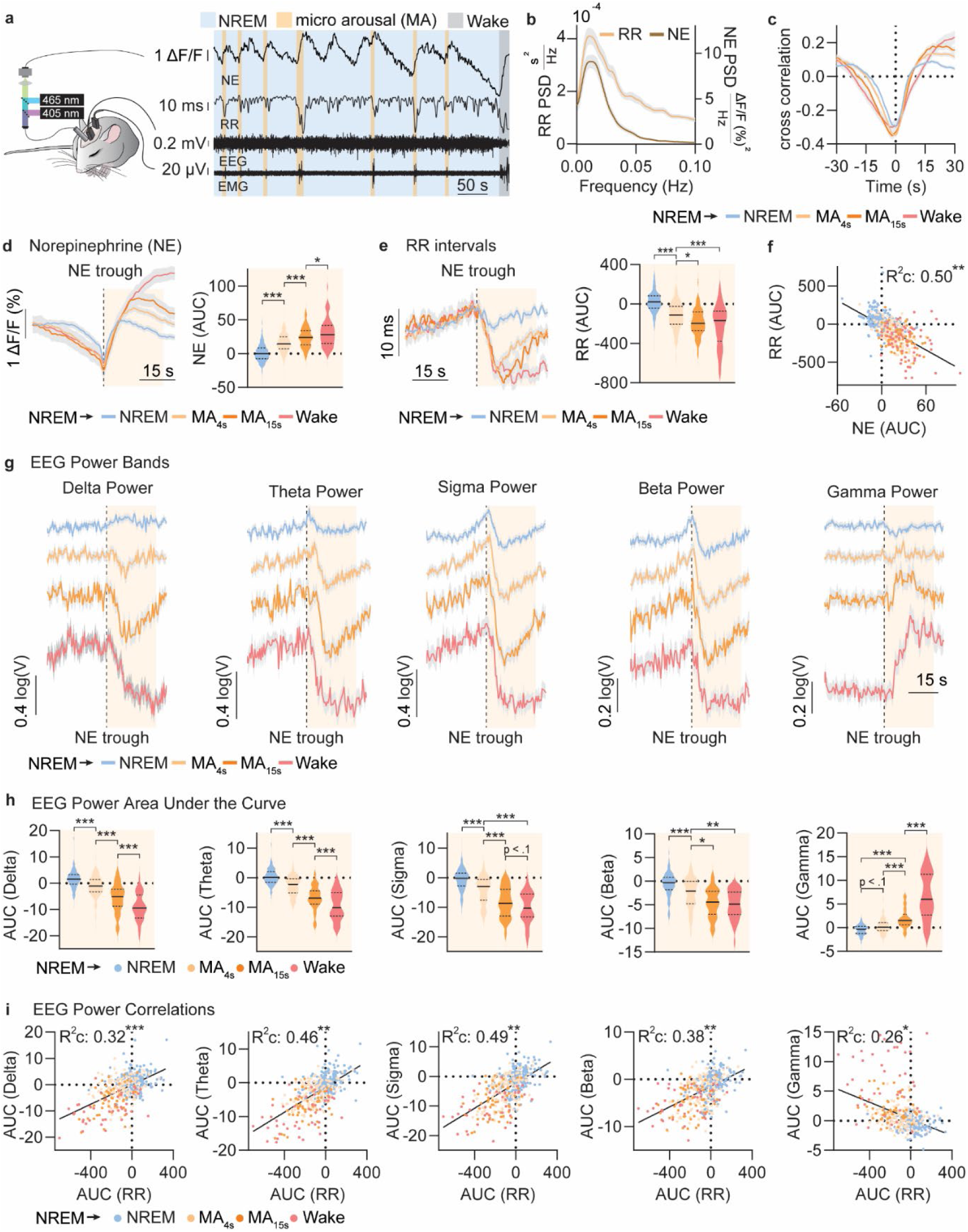
Norepinephrine and heart rate correlate during NREM sleep transitions. **a.** Left: Experimental setup visualizing the fiber photometric acquisition of extracellular norepinephrine (NE) levels in the medial prefrontal cortex, EEG, and EMG recordings. Right: Example trace of NE, EEG, EMG, and RR intervals (RR) during NREM sleep (blue shade), microarousals (MAs) <15 s (yellow shade), and wake (gray shade). RR ranged from 0.0980 to 0.1439 (417-612 BPM). **b.** Power spectral density (PSD) of RR and NE from 0-0.1 Hz. **c.** Cross correlation (how strongly and at what temporal offset the two signals covary) between NE and RR during transitions from NREM sleep to NREM sleep (blue), short MAs (beige), long MAs (orange), and wake (pink). These color codes were used for the rest of the figure. **d.** Left: mean trace with standard error of the mean (SEM) of NE with yellow background shade for parts of the trace that are quantified through area under the curve (AUC) estimations. Right: AUC for time 0 to 25 s after the NE trough with AUC of -25 to 0 as baseline correction. **e.** Left: mean trace with SEM of RR during sleep transitions. RR ranged from 0.105 to 0.13 (462-571 BPM). Right: RR AUC for time 0 to 25 after the NE trough with AUC of -25 to 0 as baseline correction. **f.** Correlation between RR AUC and NE AUC across all sleep transitions. **g.** Mean and SEM of EEG power bands surrounding the NE trough event marker during sleep transitions. **h.** AUC for time 0 to 25 after the NE trough for EEG power bands with AUC of - 25 to 0 as baseline correction. **i.** Correlations between the AUC of RR and the EEG power bands. n=7 mice, NREM events = 245, NREM to short MA events = 174, NREM to long MA events = 75, NREM to wake events = 66. *p < 0.05., **p < 0.01., and ***p < 0.001. Data is shown as mean±SEM. All frequency domain analysis figures have visualizations based on weighted estimates (see methods). For a more detailed overview of the statistics, see section 1c-i in ‘Statistics’ under Supplementary Materials.

From the simultaneous recording of NE and EMG-derived RR, example traces suggested that there was an inverse correlation between NE and RR, where RR would decrease (rapid burst in HR) around the same time NE would increase (Fig. 1a). To highlight the potential temporal relationship between NE and RR, we conducted a frequency-domain analysis of the infraslow fluctuations in both signals (Fig. 1b, Supplementary Fig. 1e-f).

To explore the NE-RR relationship across sleep-wake transitions, we examined four progressive sleep-to-wake transitions using the preceding NE trough as the time stamp (time 0): continued NREM (lowest arousal), NREM to short MA (movement ≤ 4 s) transitions, NREM to long MA (movement 5 ≤ 15 s), and NREM to wake (movement > 15 s, highest vigilance state). Multiple transition events were extracted from each animal’s recording and pooled across animals for subsequent analysis. Cross-correlation analysis revealed a predominantly negative relationship between NE and RR across vigilance-state transitions, indicating that increases in NE were associated with reductions in RR (i.e., faster heart rate; Fig. 1c). Additionally, we found that the magnitude of NE ascent corresponded to progressively greater shifts in vigilance state during sleep (Fig. 1d). Similarly, when comparing the RR, we found a corresponding drop in values (faster HR) across state transitions (Fig. 1e). When correlating the NE and RR changes across the sleep transitions, a negative correlation was found (Fig. 1f), indicating that phasic fluctuations in NE and peripheral autonomic activity were highly correlated. These comparisons were performed using area under the curve (AUC) estimates of NE and R-R responses. Importantly, comparing AUC and peak amplitude measures showed a similar overall relationship, and the coupling between NE and RR remained significant when only peak amplitudes were considered (Suppl. Fig. 1g), indicating that the findings are not solely driven by the response duration aspect of AUC. The somewhat stronger relationship observed with AUC-based measures may reflect rapid saturation of heart-rate responses across vigilance-state transitions, making response persistence an informative component of the physiological signal.

To appropriately categorize the different vigilance events, we calculated the power of delta (1 - 4 Hz), theta (4 - 8 Hz), sigma (8 - 15 Hz), beta (15 - 30 Hz), gamma (80 - 100 Hz) bands from the spectrum of EEG signals surrounding NE throughs (Fig. 1g). Power in all bands, except gamma, decreased as the vigilance state increased (Fig. 1h). Inversely, as vigilance state increased, gamma also increased as reported before (Osorio-Forero et al., 2025). Lastly, changes in band power were correlated with the decline in RR across all sleep transitions. All EEG bands showed a positive correlation with RR, except for gamma, which was negatively correlated (Fig. 1i). Thus, these results indicate that NE-locked RR changes predicted EEG-defined vigilance states.

In conclusion, changes in cardiac rhythm are tightly coupled with the infraslow phasic NE fluctuations during NREM sleep, and HR accelerations may serve as a proxy for estimating temporal arousability dynamics during sleep. Interestingly, the systematic periodicity of this heart-brain coupling falls within the very low frequency (VLF) band (0-0.15 Hz for mice (Pizzo et al., 2022)), previously noted in the heart rate variability literature (Berntson et al., 1997; Tobaldini et al., 2013).

### Closed-loop optogenetic activation of the locus coeruleus during NREM sleep increases heart rate

After finding the inverse correlation between NE and RR in natural sleep transitions, we next sought to characterize the causal influence of LC activity on HR dynamics, using a closed-loop optogenetic approach to modulate NE oscillatory frequency during NREM sleep. To this end, we designed a stimulation paradigm to modulate NE oscillatory frequency during NREM sleep. Specifically, we optogenetically stimulated the LC bilaterally in TH-Cre transgenic mice through Cre-dependent Channelrhodopsin (ChR2) activation. We employed a closed-loop paradigm, where LC stimulations (2 s 20 Hz (10 ms) blue laser pulses with a light intensity of 5 mW) were triggered when NE levels fell below increasing thresholds (-15, -10, -5, 0 and 5 ΔF/F (%), Fig. 2a-b, Methods). This approach enabled controlled compression of the infraslow NE cycle by triggering LC activation during the descending phase of the endogenous NE signal, thereby increasing the effective oscillatory frequency while preserving sleep continuity. This strategy allowed us to test whether heart-rate responses continue to track LC-driven NE fluctuations as the infraslow rhythm becomes progressively faster.

**Figure 2:**
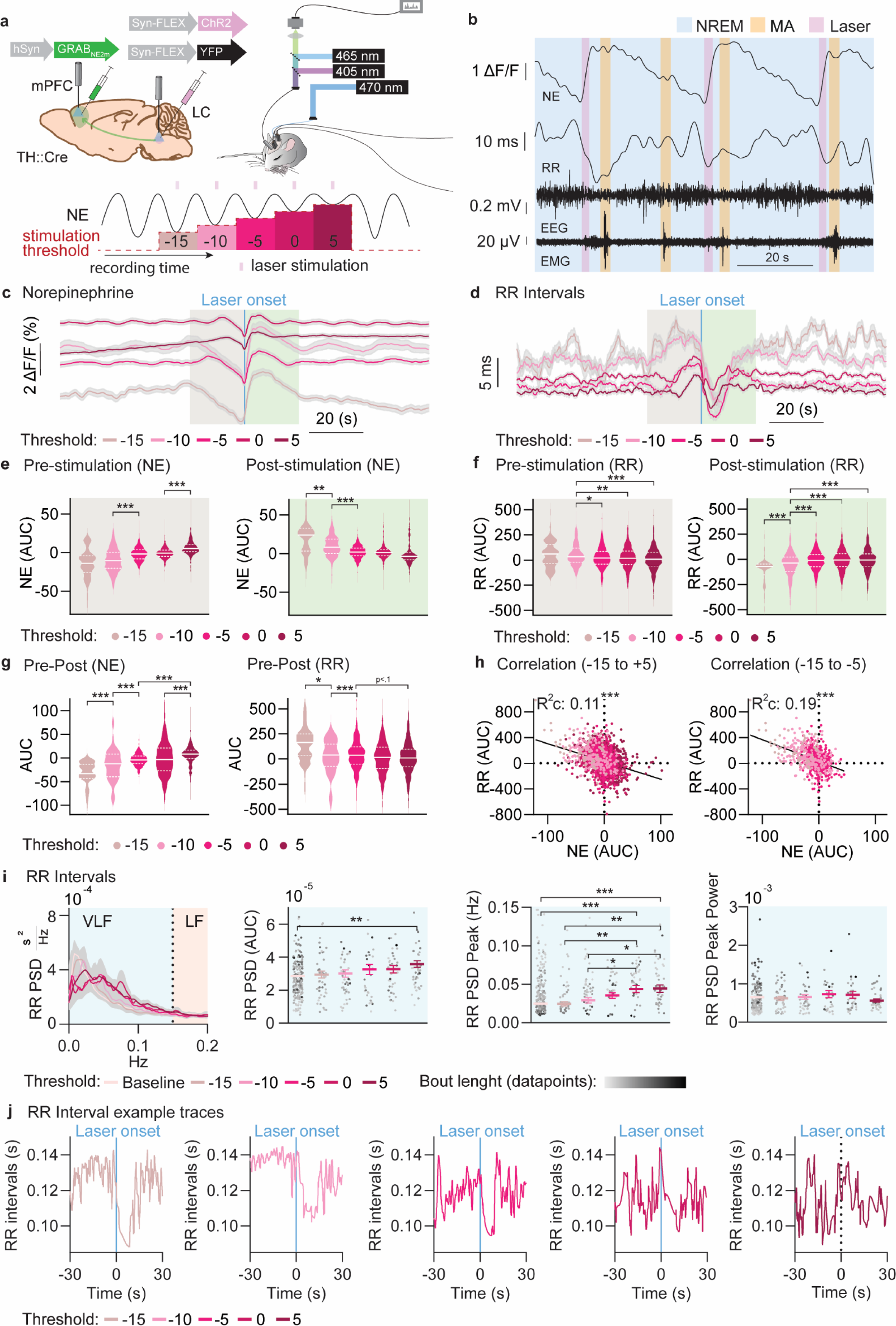
Closed-loop optogenetic activation of locus coeruleus during NREM sleep increases heart rate. **a.** Experimental overview. Top: Channelrhodopsin (ChR2) was used to enable Locus Coeruleus (LC) stimulation through light activation. NE release was recorded with fiber photometry during simultaneous EEG/EMG recordings. Bottom: LC stimulation was activated when the real-time ΔF/F (%) calculations of NE levels fell below the pre-specified thresholds of -15, -10, -5, 0, and 5 ΔF/F (%) difference of the mean ΔF/F (%) from the last two minutes. **b.** Example trace of NE, EEG, EMG, and RR intervals (RR) from the first threshold (-15) period during NREM sleep (blue shade), microarousals (MAs) <15 s (yellow shade), and laser stimulation (light pink shade). RR ranged from 0.0940 to 0.1310 (458-638 BPM). **c.-d.** Mean traces with standard error of the mean (SEM) of NE (left) and RR (right) surrounding the onset of a laser burst with beige (pre-stimulation) and green (post-stimulation) background shade for all five thresholds: -15 (beige), -10 (light pink), -5 (pink), 0 (dark pink), and 5 (wine red) ΔF/F (%) difference. These color codes remain consistent throughout the figure. RR ranged from 0.1190 to 0.1370 (438 - 504 BPM). **e.-d.** Distributions of area under the curve (AUC) of NE (**e.**) and RR (**d.**) either 20-0 s before stimulation (left) or 0-20 s after stimulation (right) for all five thresholds. Baseline estimates were subtracted from the observed values at 20-40 s before and after stimulation respectively. **g.** Pre-stimulation AUC minus the post-stimulation AUC for NE (left) and RR (right) for all thresholds. **h.** Left: correlation of the pre-stimulation AUC minus the post-stimulation AUC for NE and RR across all thresholds. Right: correlation of the pre-stimulation AUC minus the post-stimulation AUC for NE and RR across thresholds -15, -10, and -5. **i.** From left to right: 1) Power spectral density (PSD) for RR between 0-0.2 Hz across all thresholds. 2) AUC of the RR PSD across all thresholds in the very low frequency (VLF) range. 3) Frequency at the peak of the PSD across thresholds for RR in the VLF range. 4) Power value at the peak of the PSD across thresholds for RR in the VLF range. Datapoint darkness is proportional to bout length. **j.** Example traces for R-R intervals during different LC stimulations. n=10 mice (6 ChR2). Threshold -15 events = 108 (97 ChR2), Threshold -10 events = 260 (205 ChR2), Threshold -5 events = 777 (481 ChR2), Threshold 0 events = 1444 (934 ChR2), and Threshold 5 events = 1148 (771 ChR2). Number of ChR2 bouts for frequency domain analysis were, from baseline to highest threshold: 212, 44, 41, 30, 37, and 43. Data is shown as mean±SEM. *p < 0.05., **p < 0.01., and ***p < 0.001. All frequency domain analysis figures have visualizations based on weighted estimates (see Methods). For a more detailed overview of the statistics, see section 2e-i in ‘Statistics’ under Supplementary Materials.

Mean traces of NE and RR were aligned to LC stimulation onset (Fig. 2c-d, for number of laser stimulations see Fig. 2 legend or Methods). As apparent from the traces, the stimulation became more frequent with higher NE thresholds, the NE baseline increased, while the RR baseline decreased (faster HR). The magnitude of pre-stimulation NE descent and post-stimulation NE ascent was reduced as the thresholds increased, indicating less pronounced NE dynamics as LC stimulation became more frequent (Fig. 2e, for direct comparison with YFP control, see Suppl. Fig. 2c-d). A reversed pattern was observed for the RRs (Fig. 2f). These laser-induced changes were consistent with subcortical arousal responses through power reductions for sigma and beta (Supplementary Fig. 3a-c) (Lüthi and Nedergaard, 2025). An inverse NE–RR relationship was present in both ChR2 (Fig. 2h) and YFP (Suppl. Fig. 3f) groups but was significantly stronger in ChR2 animals than in YFP controls (Suppl. Fig. 3h). When exploring the shifts in NE and RR surrounding the LC stimulations, the negative association between NE and RR plateaued beyond the 3^rd^ stimulation threshold (also known as ‘5’, Fig. 2g-h). These findings indicate a causal relationship in which LC-mediated NE modulation affects HR; however, this relationship breaks down at higher infraslow LC activation frequencies. Since pre-stimulation NE baseline levels progressively became higher in the ChR2 condition compared with YFP controls (Fig. 2c+e, Supplementary Fig. 2a-f), it demonstrates that they arise from stimulation-dependent modulation of noradrenergic tone rather than nonspecific signal drift. As a result, the pre-stimulation state at higher thresholds differed between ChR2 and control conditions, which should be considered when interpreting the immediate effects of laser stimulation across thresholds. To understand if there was a mechanistic link between the PSD overlap between NE and RR observed in Fig. 1c, the PSD was calculated for RR across stimulation thresholds (Fig. 2i). As LC stimulation became more frequent (higher thresholds), the peak frequency of the VLF range increased, indicating that the VLF band might be driven by LC activations (Fig. 2i). Surprisingly, the overall amount of power for the PSD also increased without changes to the peak power level, suggesting that a wider range of fluctuations, and thus a wider frequency distribution, emerge as LC stimulation increases (Fig. 2i).

In conclusion, we demonstrate a causal relationship between LC and HR, showing that LC activations drive HR bursts and that HR reliably tracks infraslow NE oscillations under physiological conditions. However, as the frequency of infraslow NE fluctuations increases beyond the normal range, this relationship progressively weakens, indicating that HR primarily reflects LC–NE activity during intact, restorative infraslow sleep regimes.

### Heart rate decelerations in response to sustained high norepinephrine levels

Next, we wanted to explore the unexpected widening of R-R frequency distribution in response to more frequent LC stimulation (Fig. 2i) in more detail. Although the mean trace of the RRs at higher LC stimulation thresholds did not reveal such dynamics, the variability of RRs was elevated compared to lower thresholds (Fig. 3a-b). Mean HR also increased as LC stimulation became more frequent, and this correlated with the variability of the RRs (Fig. 3c-d). Thus, HR fluctuations became more frequent with more stimulation of LC. This was also evident from individual raw RR traces (Fig. 3e).

**Figure 3:**
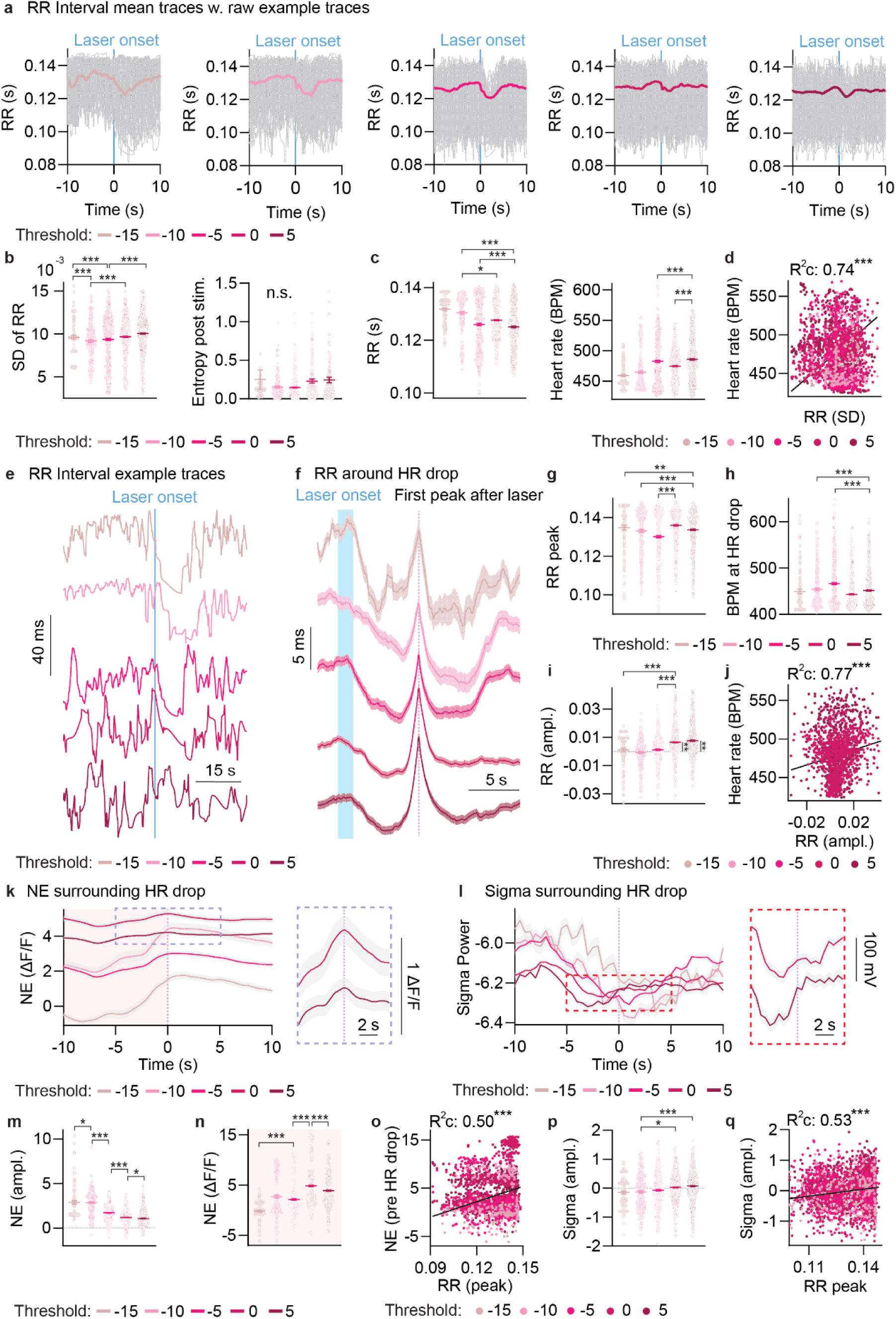
Heart rate decelerations in response to sustained high cortical norepinephrine levels. **a.** Mean RR intervals (RRs) surrounded by raw RR traces for the five thresholds: -15 (beige), -10 (light pink), -5 (pink), 0 (dark pink), and 5 (wine red) ΔF/F (%) difference. Color codes in this figure remain consistent. **b.** Standard deviation (SD) of the RRs +/- 30s around the LC stimulation (left). Sample entropy of RRs 10 s after LC stimulation (right). **c.** Mean RRs +/- 30s around the LC stimulation (left). Mean heart rate (HR) measured in beats per minute (BPM) +/- 30s around the LC stimulation (right). **d.** Correlation between HR and SD of RRs across stimulation thresholds. **e.** RR example traces surrounding laser onset. **f.** Mean RR traces surrounding HR decelerations. **g.** RR value at HR decelerations. **h.** HR at HR decelerations. **i.** RR amplitude leading up to HR decelerations. **j.** Correlation between HR and RR value at HR decelerations across thresholds. **k.** Norepinephrine (NE) mean-trace surrounding HR decelerations across thresholds (left). NE mean-trace surrounding HR decelerations at thresholds 0 and 5 (right). **l.** Sigma power mean-trace surrounding HR decelerations across thresholds (left). Sigma power mean-trace surrounding HR decelerations at thresholds 0 and 5 (right). **m.** NE amplitude leading up to HR decelerations. **n.** NE mean values leading up to HR decelerations. **o.** Correlation between mean NE values and RR values at HR decelerations across thresholds. **p.** Sigma power amplitude leading up to HR decelerations. **q.** Correlation between sigma power amplitude and RR values at HR decelerations across thresholds. Color codes for laser thresholds: -15 (beige), -10 (light pink), -5 (pink), 0 (dark pink), and 5 (wine red). n=6. Threshold -15 events = 97 ChR2, Threshold - 10 events = 205, Threshold -5 events = 481 ChR2, Threshold 0 events = 934 ChR2, and Threshold 5 events = 771 ChR2. Data is shown as mean±SEM. *p < 0.05., **p < 0.01., and ***p < 0.001. For a more detailed overview of the statistics, see section 3b-q in ‘Statistics’ under Supplementary Materials.

To quantify the HR decelerations that happened after LC activation, we took the maximal RR value (so slowest HR) 2-7 s after LC stimulation and baseline corrected the value to the mean RR during the 8–10 s period preceding the RR peak to obtain their amplitude. We found that the amplitude only differed from 0 in the higher stimulation thresholds (0 and 5) and correlated with mean HR (Fig. 3 f-j). Thus, it seems that at high LC stimulation, where HR is elevated, HR decelerations occur. This may reflect compensatory decelerations in response to sustained elevated sympathetic activity and thus HR (Airaksine et al., 2001). To understand the potential mechanism behind the HR decelerations, we explored NE and sigma power surrounding the HR decelerations (Fig. 3k-l). As LC stimulation frequency increased, general NE levels elevated, while infraslow NE fluctuations shrank (Fig. 3m-n). Notably, larger HR decelerations correlated with a higher preceding mean NE value, further supporting the interpretation that HR decelerations occurred when NE levels were high and sustained over time (Fig. 3o). Lastly, we observed that as LC activations increased, the sigma power amplitude in the seconds leading up to the HR decelerations shifted from negative to positive, indicating that HR decelerations were associated with sigma power increases (Fig. 3p). The sigma amplitude furthermore positively correlated with the RR value at the HR peak (Fig. 3q). This indicates that HR decelerations, despite high NE levels, are still associated with neuronal synchronization in the sigma frequency range, since NE can still display subtle fast declines (Fig. 3k).

In conclusion, frequent LC activations do not lead to sustained elevation in HR; rather, the heart exhibits compensatory HR decelerations, leading to large dynamic HR shifts following sigma changes.

### Heart rate responses to locus coeruleus suppression reflect memory consolidation

To complement the experiments increasing infraslow NE frequency, we next tested how slowing the infraslow NE rhythm affected the association between RR and NE by expressing a Cre-dependent inhibitory opsin, Archaerhodopsin (Arch), in the LC of TH-Cre transgenic mice, with YFP serving as a control. We measured NE levels through an optic fiber implanted in mPFC in addition to EEG/EMG measurements (Fig. 4a-b). When exploring the NE and RR response to 2 min (4 min stimulus-interval) LC suppression during NREM sleep, we observed a gradual NE decline sustained throughout the laser period that was not observed in the YFP condition, while HR did not deviate between Arch and YFP (Fig. 4c-e). Notably, despite the absence of a robust group-level HR effect, NE and RR responses remained negatively correlated across trials (Fig. 4e), indicating that HR dynamics continued to track the magnitude of noradrenergic suppression at the individual-response level. A similar negative relationship was observed in YFP controls (Fig. 4e), and the strength of the NE–RR association did not differ significantly between Arch and YFP animals (Suppl. File, Statistics), suggesting that LC suppression did not substantially alter the underlying coupling between these measures. To better understand how LC suppression impacted the NE and RR oscillations, a frequency-domain analysis was conducted for NE and RR for YFP and Arch mice (Fig. 4f). When exploring the PSD for NE and RR (VLF), the analysis revealed no shifts in infraslow dynamics between the Arch or YFP condition for either NE or RR (Fig. 4f-g). This likely reflects the relatively mild and intermittent LC suppression paradigm, which was designed to remain within physiological ranges and therefore did not globally reorganize infraslow sleep dynamics.

**Figure 4:**
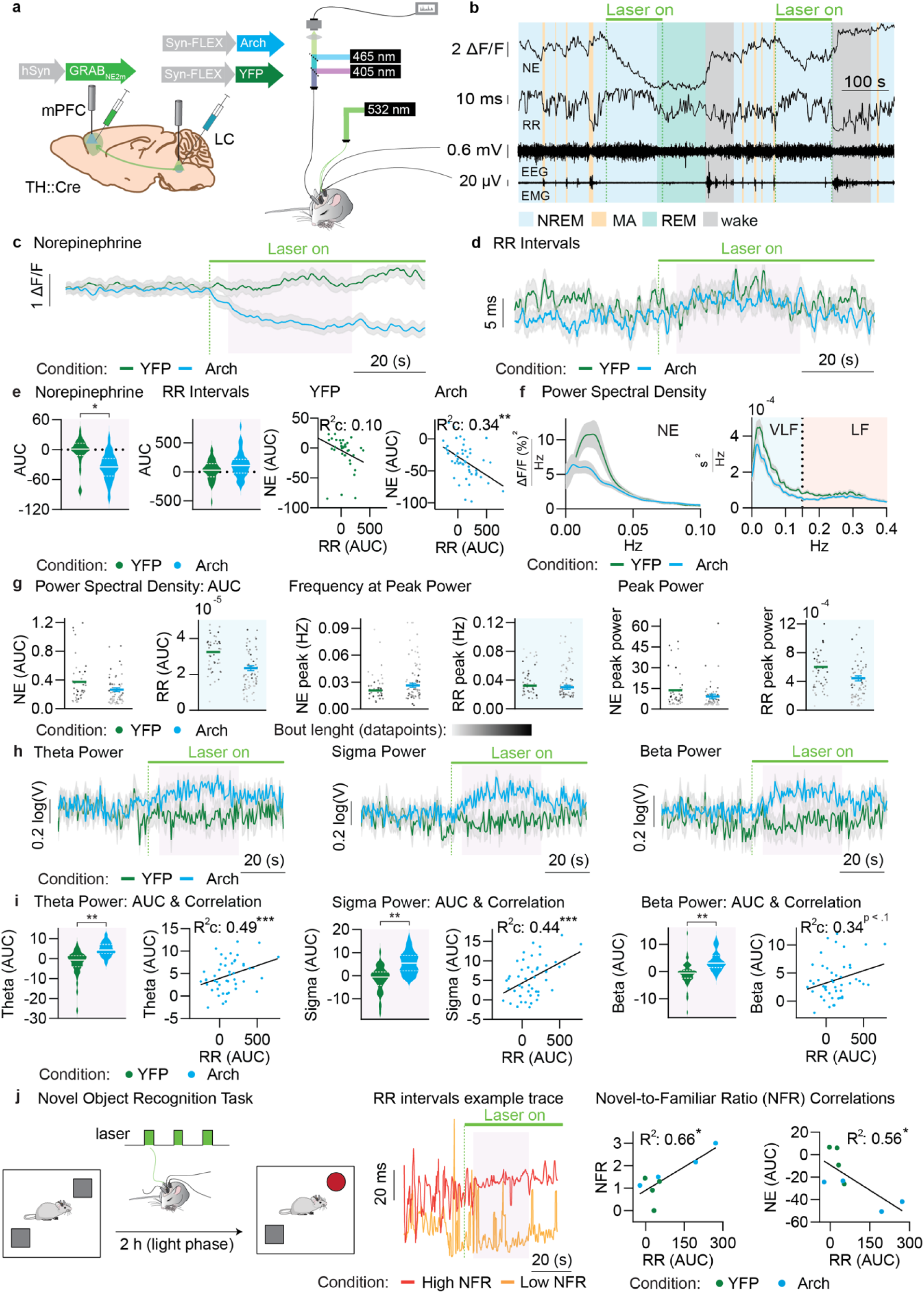
HR modulation to LC suppression improves memory consolidation. **a.** Experimental overview: Archaerhodopsin (Arch) or Yellow fluorescent protein (YFP) was injected into the locus coeruleus (LC) while GRAB-NE2m(3.1) was expressed in the medial prefrontal cortex (mPFC). Arch was stimulated through green light activation, leading to decreased norepinephrine (NE) release in the mPFC. NE release was recorded with fiber photometry during simultaneous EEG/EMG recordings. Mean HR across the recordings for the Arch group was 525 BPM (+/- 60 SD) or 0.1143 (+/- 0.0131 SD) s between beats and for the YFP group 517 BPM (+/- 61 SD) or 0.1161 (+/- 0.0137 SD) s between beats. **b.** Example trace of NE, EEG, EMG, and RR intervals (RR) during NREM sleep (blue shade), microarousals (MAs) <15 s (yellow shade), REM sleep (green shade), and wake (gray shade). RR ranged from 0.0949 to 0.1453 (413-632 BPM). **c.-d.** Mean trace with standard error of the mean (SEM) of NE (left) and RR (right) surrounding the laser onset event marker with purple shade to mark the area quantified through area under the curve (AUC) calculations for Arch-(blue) and YFP- (green) related animals. RR ranged from 0.1109 to 0.1220 (492-541 BPM). **e.** AUC of the NE trace (left) and RR trace (middle) under LC suppression with the two measures correlated (right) for Arch- (blue) and YFP- (green) related animals. **f.** Power spectral density (PSD) of NE (left) and RR (right) for YFP mice (green) and Arch mice (blue). **g.** From left to right: 1-2) AUC of NE PSD and RR PSD in the very low frequency (VLF) range (blue shade), 3-4) frequency at the peak of the NE and RR PSD, and 5-6) power at the peak of the NE and RR PSD for YFP mice (green) and Arch mice (blue). Datapoint darkness is proportional to bout length. **h.** Mean trace with SEM of theta power (left), sigma power (middle), and beta power (right) surrounding the laser onset event marker. Purple shade marks the area quantified through AUC calculations for YFP (green) and Arch (blue) animals. **i.** AUC for theta (left), sigma (middle), and beta (right) during laser activation for Arch- (blue) and YFP- (green) related animals along with their correlations with RR AUC for the Arch condition. **j.** Left: Overview of the Novel Object Recognition task. Mice were habituated to the environment before neural recordings and were then tested in the same environment with one object being replaced with a novel one. Middle: Representative RR interval traces from a high novel-to-familiar ratio (NFR) mouse and a low NFR mouse during inhibition. Right: NFR correlated with RR AUC, and RR AUC correlated with NE AUC for Arch- (blue) and YFP- (green) related mice. n=8 (4 Arch). Arch events = 48, YFP events = 38. PSD bouts: Arch = 66, YFP = 1. 43. Data is shown as mean±SEM. *p < 0.05., **p < 0.01., and ***p < 0.001. All frequency domain analysis figures have visualizations based on weighted estimates (see methods). For a more detailed overview of the statistics, see section 4e-j in ‘Statistics’ under Supplementary Materials.

Lastly, we were interested in understanding the relationship between HR during LC suppression and memory consolidation. As sleep spindle density during NE decline correlate with memory performance (Kjaerby et al., 2022), we wanted to understand if changes in HR would correlate with post-sleep performance and thus if the HR/NE coupling during this phase is important for memory consolidation. We confirmed a sigma power upregulation and general neuronal synchronization during LC suppression. Furthermore, we found a correlation between RR responses to LC suppression and sigma power, suggesting that a stronger HR reduction response is linked to higher spindle power; similar correlations were also observed in the theta and beta frequency ranges (Fig. 4h-i). To test whether HR changes were associated with memory outcomes, we performed a novel object recognition task. The mice were habituated to an environment with two objects prior to the LC suppression paradigm and were subsequently re-introduced to the environment after the recording, but now with one known object exchanged for a new one. By measuring approaches to the new vs. old object, the novel-to-familiar ratio was extracted and used as a behavioral measure of memory consolidation (Fig. 4j). Interestingly, across pooled Arch and YFP animals, larger RR increases following LC suppression were associated with better subsequent memory performance. Additionally, RR and NE responses to LC suppression were negatively correlated indicating that animals showing stronger NE reductions also exhibited larger RR changes. Together, these findings suggest that heart-rate dynamics covary with noradrenergic responses during sleep and may reflect physiological processes relevant for sleep-dependent memory consolidation.

### Sigma power surges prior to heart rate burst predict next day memory performance

To further understand the cross-species relationship between shifts in autonomic activity and memory consolidation, we explored sigma power dynamics surrounding phasic HR accelerations, also known as HR bursts (HRBs, increases in HR larger than 2 or 2.2 SD of the mean in windows of 180 or 24 s for humans or mice respectively) (Chen et al., 2022b, 2022a, 2020; Naji et al., 2019a, 2019b), in both humans and mice. In mice, sigma power has been previously shown to inversely correlate with NE fluctuations and predict memory performance (Kjaerby et al., 2022). As expected, NE increases in mice preceded phasic HRBs and peaks at the time of the HRB (Fig. 5b-c). Additionally, the amplitude of the NE increases associated with the strength of the HRBs (Fig. 5d). We showed that sigma power dropped directly prior to HRBs, consistent with our expectation given the negative correlation between sigma power and NE in prior work (Kjaerby et al., 2022) (Fig. 5e-g). A similar analysis was performed in a human dataset, where participants underwent memory testing and EEG/ECG recordings overnight. While the relationship between sigma power and RR intervals was conserved across species, we observed a temporal shift in the human data (Fig. 5h). Here, the surge in sigma power preceding the HRBs was confined to the 5-s interval prior to the HRB (Fig. 5i-j), while the sigma for mice peaked in the 15-10 s range prior to the event. These temporal differences may reflect species-specific physiology differences in cardiac timescales, which has also been previously reported (Bergel et al., 2025; Carro-Domínguez et al., 2025; Lecci et al., 2017). When analyzing the relationship between the pre-HRB sigma power and the slowing of the heart following the HRB in humans, we observe a positive correlation, suggesting the importance of the pre-HRB sigma power for HR baseline recovery (Fig. 5k). Lastly, we measured improvements in hippocampal-dependent, episodic memory through the word-pair association (WPA) task before and after a night of sleep (Fig. 5l). Sleep-dependent memory improvement (post-pre-difference) was associated with pre-HRB sigma power (baseline-corrected, p = 0.05), suggesting that surges in spindle activity linked to HRBs were important for memory consolidation (Fig. 5m).

**Figure 5.**
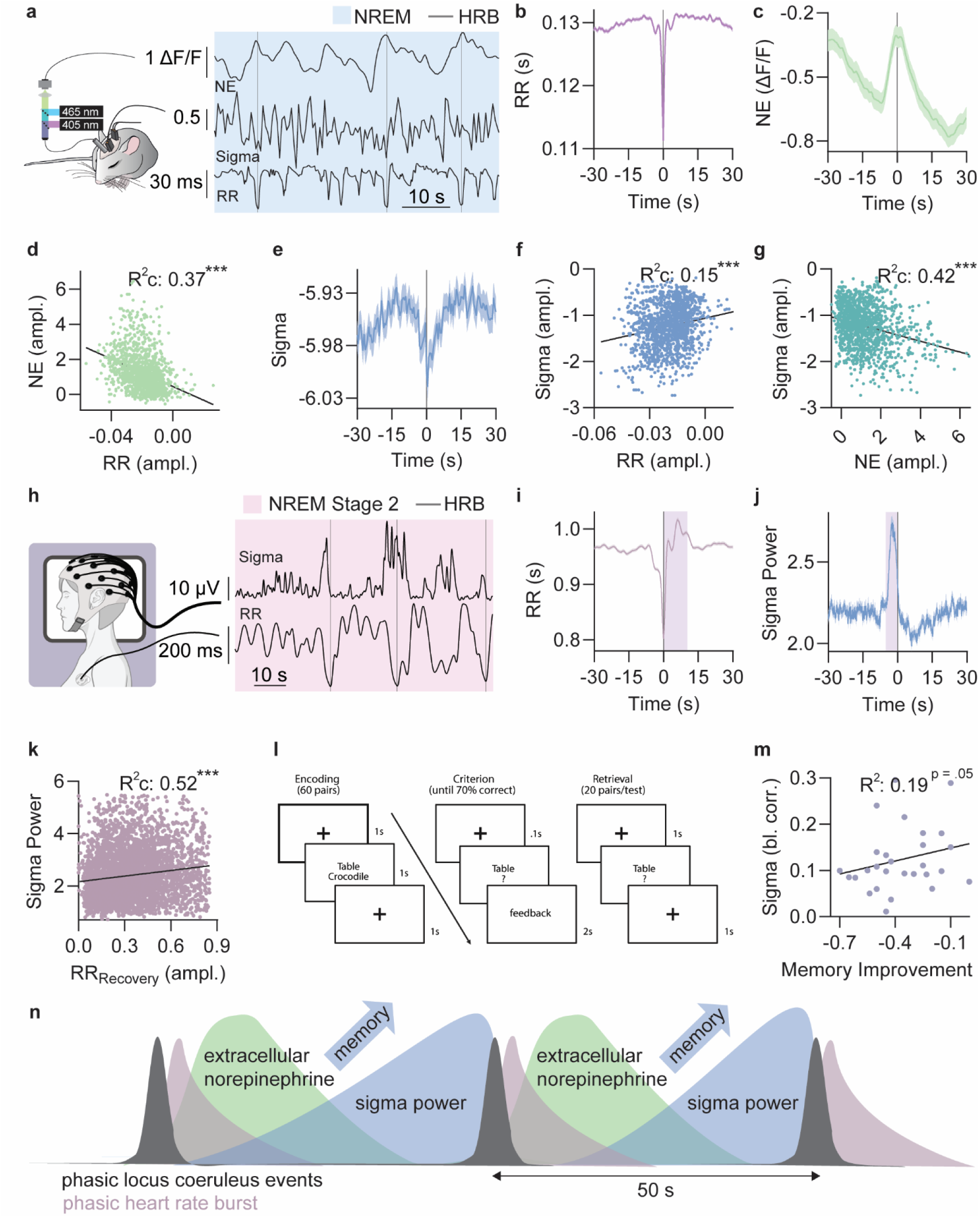
Pre ‘Heart Rate Burst’ Sigma surges linked to improved memory. **a.** Example trace showing norepinephrine (NE), sigma power, and RR-interval (RR) during NREM sleep (blue shade) and heart rate burst (HRB) events (gray lines). **b.-d.** Mean trace with standard error of the mean (SEM) of RR (left), NE (right), and sigma power (below) surrounding the HRB events. **e-g.** Correlation between the NE and RR amplitude (left), Sigma power and RR amplitude (middle), and Sigma and NE (right) amplitude. **h.** Example trace of human RR and sigma power during NREM stage 2 sleep (pink shade) during HRB events (gray lines). **i.** Mean trace of RR intervals surrounding HRB events. Purple shade represents the time during which the RRrecovery amplitude was measured. **j.** Mean trace of sigma power intervals surrounding HRB events. Purple shade represents the sigma power interval quantified in following sub-figures. **k.** Correlation between sigma power pre-HRB and RR amplitude following the HRB 1. **l.** Overview of the word-paired association (WPA) memory task. **m.** Correlation between memory improvement and baseline corrected pre-HRB sigma power. Mice: n=7, HRB events = 920. **n.** Schematic diagram showing the relationship between phasic locus coeruleus and heart rate events, which are followed by norepinephrine descends and sigma power upregulations correlating with memory performance. Hypothesis figure: Humans: n=28, HRB events = 4753. Data is shown as mean±SEM. *p < 0.05., **p < 0.01., and ***p < 0.001. For a more detailed overview of the statistics, see section 5d-m in ‘Statistics’ under Supplementary Materials.

In conclusion, mice and humans displayed evidence of conserved autonomic-central coupling, reflected in coordinated HR and sigma power dynamics surrounding phasic cardiac events.

However, the temporal relationship between these events differed across species, likely reflecting differences in sleep and cardiovascular physiology. In mice, causal manipulation of LC activity demonstrated that HR dynamics covary with LC-mediated noradrenergic and spindle-related processes linked to memory consolidation. In humans, sigma power preceding HR bursts was associated with both post-HRB heart-rate slowing and sleep-dependent memory improvement, suggesting that autonomic–central coupling surrounding HR events may provide a translational marker of restorative sleep processes (Fig. 5n).

## Discussion

Variability in HR is a noninvasive biomarker of autonomic nervous system function and is frequently disrupted in aging and Alzheimer’s disease. Here, by combining real-time measurements of mPFC NE dynamics with simultaneous HR recordings in freely sleeping mice, we demonstrate that HR closely tracks the infraslow phasic activity of the LC–NE system. Specifically, we show that infraslow NE oscillations are reflected in HRV within the VLF range, providing mechanistic evidence that this previously poorly understood HRV component indexes central noradrenergic dynamics. When infraslow NE activity shifts toward faster frequencies, as observed during sleep fragmentation, HR dynamics follow these changes only within a limited range, after which compensatory mechanisms begin to obscure the coupling, suggesting a physiological boundary to brain–heart synchrony. Importantly, we further demonstrate that HR predicts the memory-relevant interaction between NE activity and sleep spindle organization in both mice and humans, identifying HRV as an accessible peripheral marker of sleep-dependent memory consolidation. Together, these findings bridge LC-mediated regulation of sleep microstructure with established HRV metrics and support VLF-HRV as a translational readout of restorative sleep processes and central noradrenergic function.

### Heart rate is modulated by infraslow norepinephrine fluctuations during NREM sleep

Utilizing HR detection and fiber photometry, we found that NE and HR correlate across NREM sleep transitions. To determine causality, we utilized TH-Cre mice with ChR2/YFP injected in the LC to activate the nucleus. We found that LC stimulations were associated with increases in HR, implying that NE levels mediate HR changes.

Many studies have linked HR to NE (Fawaz and Simaan, 1963; Sundaram et al., 1991; Tanoue et al., 2022; Watson et al., 1979). Many rely on plasma levels of NE, which, while linked to central NE (Gurguis and Uhde, 1998), has low temporal resolution, making causal interpretations harder. In recent years, the use of biosensors and fiber photometry allows for very reliable estimate of the temporal dynamics of NE changes making association to HR more precise (Osorio-Forero et al., 2021). The periodicity of phasic fluctuations of LC and HR overlaps and corresponds with previous findings from HR studies, namely those centered on exploring HRV. HRV is a well-established metric within the health sciences literature (Berntson et al., 1997), with greater HRV consistently associated with a broad spectrum of positive health and cognitive outcomes, many of which are linked to frontal lobe functioning (Forte et al., 2019). HRV is conventionally analyzed across three primary frequency bands: high frequency (HF), low frequency (LF), and very low frequency (VLF) (Berntson et al., 1997). While HF HRV and LF HRV and their relation to autonomous functioning have been characterized (Berntson et al., 1997; Malik et al., 1996; Shaffer et al., 2014), the physiological mechanisms underlying VLF HRV remain poorly understood (Bonaduce et al., 1994; Brämer et al., 2019; Kitney, 1980). The frequency range for VLF in humans has been reported to be ≤0.04 Hz (Berntson et al., 1997; Tobaldini et al., 2013) but have in mice been extended to ≤0.15 Hz (Pizzo et al., 2022) due to the faster HR dynamics.

Here, we show that VLF HRV power overlaps with NE infraslow phasic fluctuations (0-0.1 Hz) in rodents, and recent evidence from pupil response in humans suggests that this overlap is conserved cross-species (Carro-Domínguez et al., 2025). Additionally, stimulating the LC pushed VLF HRV towards higher frequencies, suggesting that one potential driving force of VLF HRV is LC-mediated NE infraslow oscillations. Prior findings demonstrate that LC is linked to the autonomic nervous system. Stimulation of LC projections decrease parasympathetic cardiac vagal activity (Wang et al., 2014) and also influences sympathetic output through direct projections to the preganglionic cells in the sympathetic nervous system (Karemaker, 2017; Nygren and Olson, 1977; Samuels and Szabadi, 2008). While the mechanisms generating VLF HRV are not well defined (Armour, 2003; Shaffer et al., 2014; Wang et al., 2014), there is a clear parasympathetic component (Taylor et al., 1998). This combined with the ability of pharmacological blockage of the parasympathetic system to block infraslow oscillations of HR (Osorio-Forero et al., 2021) and pupil diameter during sleep (Yüzgeç et al., 2018) led us to expect a slowing of HR during LC suppression due to parasympathetic disinhibition. Interestingly, we found no consistent modulation in HR during LC suppression, suggesting that the LC-HR connection may also partly be driven by sympathetic outflow. Previous research had indicated that LF power may also represent sympathetic activity, but this interpretation has been challenged due to the mixed contribution of both autonomic branches (Houle and Billman, 1999; Japundzic et al., 1990; Reyes et al., 2013). LF and HF ratio (LF/HF) were traditionally thought to reflect balance between sympathetic and parasympathetic activities (i.e., the sympatho-vagal balance), though currently considered as an oversimplification of non-linear integration of autonomic signals (Billman, 2013; Pagani et al., 1986). Our findings implicate the VLF may offer a precise marker for central arousal states, given its overlaps with infraslow phasic fluctuations of LC-NE levels. *These comparisons were performed* important for the expression of VLF oscillations, growing evidence suggests that VLF dynamics reflect broader interactions between the heart and autonomic nervous system rather than simple sympathetic or parasympathetic control alone (Armour, 2003; Shaffer et al., 2014). Within this framework, infraslow LC–NE dynamics may represent one central contributor to these slow cardiac fluctuations during sleep either directly through sympathetic activation or indirectly by inhibition of cardiac vagal activity. Furthermore, heart rate and HRV may be a promising highly accessible read-out of LC activity, which has important clinical implications.

### Heart rate as a cross-species memory marker

Spindle density, implicated in sleep-dependent memory consolidation, increases during periods of NE decline (Kjaerby et al., 2022). Specifically, the infraslow descent of NE creates spindle-rich periods. Now we report a correlation between memory consolidation and coupled HR-NE decrease during LC suppression, suggesting that HR can be used as a biomarker of memory consolidation in sleep. This finding was extended to humans, where the sigma power surges prior to heart rate bursts (HRBs) predicted sleep-dependent memory improvement. In mice, we saw that the HRBs co-occurred with a peak in NE, which had been on the rise prior to HR changes. Since NE changes cause HR alterations, this suggests that HRBs are triggered by increases in NE signaling in mice. Thus, a similar NE increase may exist in humans (Carro-Domínguez et al., 2025).

Our findings also suggest that the pre-HRB surge in sigma power could play a role in memory consolidation. Previous work in humans demonstrated that the strength of infraslow sigma oscillations correlates with sleep-dependent memory consolidation in humans (Lecci et al., 2017). Thus, given that HRB follows the infraslow NE rhythm, the specific timing of the sigma power relative to the HRB might be important for memory consolidation. These findings replicate past findings (Naji et al., 2019a) in daytime sleep and further generalize them to overnight sleep and thus overnight memory changes.

Interestingly, when comparing mouse data to human data, we see temporal differences in the sigma power leading up to the HRB. The mouse sigma peaks around 10 s earlier and has more oscillatory dynamics as opposed to the stark rise in sigma power pre-HRB in humans. This pattern has previously been reported (Bergel et al., 2025; Carro-Domínguez et al., 2025; Lecci et al., 2017) and has been attributed to species-specific differences in anatomy and physiology in the neural coupling to the heart (Lecci et al., 2017). These species-specific differences could involve intrinsic cardiac neuron (ICN) properties and cardiac neural circuitry. A comparative study shows that mouse ICNs exhibit more phasic firing and direct cholinergic projections to the sinoatrial node, enabling faster and more oscillatory heart-brain coupling, while human ICNs are predominantly tonic, supporting slower, more sustained modulation (Tompkins et al., 2024). These anatomical and physiological differences likely contribute to the earlier and more dynamic sigma peak observed in mice compared to humans and further research efforts to map cross-species heart-brain interactions may reveal critical differences that can be used in translational techniques.

Importantly, infraslow fluctuations of sigma power in human sleep have received growing attention. Recent work demonstrates that the infraslow fluctuation of sigma power segments N2 sleep into functional phases associated with arousal and memory-related sleep markers (Dimitriades et al., 2024). Complementary findings using pupillometry show that pupil diameter fluctuates on similar infraslow timescales during NREM sleep and is inversely related to spindle clustering, providing indirect evidence that arousal-related noradrenergic dynamics shape human sleep microstructure (Carro-Domínguez et al., 2025). However, LC activity during human sleep remains inferred rather than directly measured, and systematic perturbation studies linking LC output to spindle–autonomic coupling in humans are currently lacking. Together, these observations underscore both the promise and the current limitations of cross-species comparisons of infraslow sleep dynamics.

In summary, these findings imply that HR serves as a surrogate marker for NE in humans as well as a biomarker for memory consolidating processes during sleep. Loss of LC neurons occurs with general aging and is an early marker of Alzheimer’s disease (Grudzien et al., 2007; Mann et al., 1980). Furthermore, maintaining LC neural density prevents neurodegeneration (Wilson et al., 2013). Given the reported reduction in HRV in aging and Alzheimer’s disease (Natarajan et al., 2020; Zulli et al., 2005), suppressed LC responses might be responsible for the blunted autonomic cardiac modulation during sleep. The link to HRV could be further investigated by neuroimaging studies quantifying LC volume (atrophy) and determines whether the extent of LC loss could be a possible predictor of future cognitive decline through its relationship to sleep spindles. Indeed, evidence suggests that the LC activity during sleep is compromised (e.g., increased arousals, fragmented sleep, elevated cortisol) and linked to memory failure in both aging (Mander et al., 2017, 2013) and Alzheimer’s disease (Lim et al., 2013).

### Locus-coeruleus activation drives compensatory heart rate decelerations via autonomic feedback mechanisms

A surprising finding was that diminishing the power of NE fluctuations by repetitive LC activation did not lead to less power in the HR fluctuations; rather, the opposite happened. This was caused by large decelerations in HR in conditions with repetitive LC activation and high HR. We propose that the deceleration in HR during repeated LC stimulation may be explained by a dynamic balance between heightened sympathetic control due to increased LC activity and compensatory baroreflex responses. This is likely occurring as NE activates alpha-1 receptors on the smooth muscle in blood vessels, leading to vasoconstriction and increased blood pressure, triggering the baroreflex (Tanoue et al., 2002; Webb, 2003). This baroreflex signals to the nucleus of the solitary tract, which activates inhibitory neurons. These neurons suppress the activity of the rostral ventrolateral medulla, ultimately resulting in reflex parasympathetic activation with sympathetic inhibition lowering the HR (Aicher et al., 2000; Lanfranchi and Somers, 2002). An interesting finding was that the heart rate decelerations at high NE levels were associated with increases in neuronal synchronization (i.e., increased sigma power), which seemed to be closely linked to HR decelerations and smaller NE fluctuations. It is possible that the LC receives autoinhibitory feedback from the baroreflex responses, but that this is not reflected in our cortical NE recordings.

Overall these findings suggest that infraslow phasic fluctuations of LC-NE can drive large HR fluctuations consistent with the VLF range. However, when NE levels reach artificially high and sustained levels, compensatory mechanisms such as baroreceptor reflex activation may intervene. This adaptive feedback could serve as a safety valve, reducing the coupling between central NE activity and cardiac oscillations to prevent excessive or destabilizing cardiovascular responses.

## Conclusion

We show that LC-mediated NE release plays a role in generating VLF HRV. HR appears to be more sensitive to stimulation than inhibition of the LC, indicating a potential role for NE in modulating cardiac function. These results may point toward some contribution of sympathetic activity in addition to the well-known parasympathetic influence, though further investigation is needed to determine the precise autonomic mechanisms involved. Furthermore, we demonstrate that shifting NE fluctuations during NREM sleep toward faster frequencies, the LC-HR coupling starts to disassociate due to obscuring compensatory mechanisms. Lastly, we conclude that HR changes through their known association with sigma power reflect memory consolidation processes in mice and humans. This research opens doors to unexplored possibilities of using HR as a non-invasive, cost-effective biomarker not only for sleep quality and LC-NE function, but potentially also for conditions of cognitive decline associated with sleep disturbances.

## Methods

### Mice

This study included female and male mice, which were group-housed according to sex with unlimited access to food and water. The housing environment had a 12-hour light/dark cycle at 21°C with 40-60% humidity. The data for all mouse-based experiments were acquired in connection to past publications (Kjaerby et al., 2022). For experiments associated with sleep transitions, seven C57BL/6 (wild-type) male mice were used. The heterozygous TH-Cre transgenic mice used for optogenetic experiments were bred on a C57BL/6 background. Ten mice were included in the LC activation experiments, four of which were controls (2 males/2 females) and six of which were ChR2 (4 males/2 females), and nine were involved in the LC suppression experiments. Of the nine mice, four were put in the control condition (4 females) and four Arch (3 females/1 male). One Arch mouse was excluded because it took over one-third of the recording period to enter NREM. The number of mice included in the analysis was lower than in previous publications due to the exclusion of animals with poor EMG signal quality. All experiments received approval from the Danish Animal Experiments Inspectorate and were supervised by the University of Copenhagen’s Institutional Animal Care and Use Committee (IACUC), in accordance with the European Communities Council Directive of 22 September 2010 (2010/63/EU) on the protection of animals used for scientific purposes.

### Surgery

Mice underwent surgery at 7-15 weeks of age. General anesthesia was induced with 5% isoflurane and maintained at 1-3% isoflurane. Additionally, 0.05 mg/kg of preoperative buprenorphine was administered for general analgesia, and 0.03 mg/kg of lidocaine was injected at the incision site. A stereotactic frame was used to ensure stability and alignment throughout the surgery. The fur was trimmed from the upper part of the skull, and a cut was made between the ears. The skull was cleaned with ethanol, and an electric drill (Tech2000, RAM Microtorque) was used to make 3-5 burr holes in the skull following the stereotactic coordinates relative to the bregma.

In the mPFC burr hole, AAV9-hSyn-GRAB-NE2m(3.1) (see Table 1 for supplier information) was injected under the neuronal hSyn promoter (stereotactic coordinates: A/P +1.7 mm, M/L −0.3 mm, and D/V −2.00, −2.25, −2.50, and −2.75 mm, with 125 nL of virus injected at each level). Additionally, for mice included in the optogenetic experiments, an AAV5/2 (Adeno-associated virus 5/2) encoding floxed Arch (ssAAV5/2-shortCAG-dlox-Arch3.0-eGFP-dlox-WPRE-SV40p(A)) or ChR2 (ssAAV5/2-hEF1a-dlox-hChR2(H134R)-eYFP(rev)-dlox-WPRE-hGHp(A)) was injected bilaterally into the LC of the transgenic TH-Cre mice (stereotactic coordinates: A/P −5.5 mm, M/L ±0.9 mm, and D/V −3.75 mm, with 300 nL of virus injected). YFP (AAV5-EF1a-DIO-eYFP) was injected in control animals. For both virus injections, the needle was kept in place for 7 minutes after the injection (speed: 100 nL/min) before slow withdrawal.

**Table 1:** Information on the supplier of the different viral constructs. Adopted from Kjærby et al. (Kjaerby et al., 2022).

| **Construct** | **Company/provider** | **Cat#** |
| --- | --- | --- |
| pAAV9-hSyn-GRAB-NE2m(3.1) | Yulong Li lab | 208686 |
| ssAAV5/2-shortCAG-dlox-Arch3.0-eGFP-dlox- | Zürich vector Core | V461-5 |
| ssAAV5/2-hEF1a-dlox-hChR2(H134R)-eYFP(rev)- | Zürich vector Core | v214-5 |
| AAV5-EF1a-DIO-eYFP | UNC Vector Core | 27056- |

For EMG acquisition, two wires from W3 Wires International were placed into the musculus trapezius. For EEG acquisition, two 0.8 mm stainless steel screws from NeuroTek with low impedance were screwed into holes above the cerebellum (reference point) and the frontal cortex. Contralateral to the EEG screws, a mono fiber-optic cannula (400 μm, 0.48 NA, Doric Lenses), attached to a metal ferrule (2.5 mm diameter), was placed in the mPFC at D/V −2.50 mm. For mice that had received Arch, ChR2 or YFP injections in the LC, dual fiberoptic cannulas (200 μm, 0.22 NA, Doric Lenses) were implanted bilaterally above the LC at D/V −3.65 mm. Connectors linked to the EEG screws and EMG wires, along with the cannulas, were fixed to the head with SuperBond dental cement.

Mice received 5 mg/kg of carprofen (s.c.) prior to waking up and two days after as post-op care. They were given at least two weeks post-operation for recovery and virus expression before experimentation. The surgeries were conducted following established institutional guidelines. Viral expression and injection sites were validated through immunostaining of perfused brain slices from the experimental animals (see Kjaerby et al. (Kjaerby et al., 2022) for more information).

### Fiber photometry

Two light-emitting diodes (LEDs) with wavelengths of 465 nm and 405 nm (Doric Lenses, Tucker Davis Technologies) were connected to a minicube (Doric Lenses) using attenuator patch cords (400-μm core, NA = 0.48, Doric Lenses). The fiber-optic patch cords were secured to the fiber implants on the animals’ heads through zirconia sleeves. The minicube, equipped with dichroic mirrors and cleanup filters, enabled the detection of fluorophores by matching the relevant excitation and emission spectra. The LEDs were controlled by LED drivers (Thorlabs, Doric, Tucker Davis Technologies) and interfaced with an RZ-5 or RZ10-X real-time processor (Tucker-Davis Technologies). In the mPFC, both the 465 nm and 405 nm excitation lights were transmitted through a single patch cord to activate GRAB-NE2m(3.1) fluorescence, with the 405 nm light serving as an isosbestic control wavelength to adjust for bleaching and movement-related signal fluctuations. The 465 nm and 405 nm excitations were modulated sinusoidally at 531 Hz and 211 Hz, respectively. The two modulated signals generated by the LEDs were each retrieved using standard synchronous demodulation techniques, which were implemented on the RZ-5/RZ10-X real-time processor at a sampling rate of 1000 Hz. Signal processing was managed using Synapse software (Tucker-Davis Technologies), which also synchronized the fluorescent signals with video and EEG/EMG signals via TTL pulses. The resulting data were then exported for further analysis in MATLAB R2023b (MathWorks).

To calculate ΔF/F, the 405 nm signal was fitted to the 465 nm signal using a first-degree polynomial regression (MATLAB polyfit function). The slope and intercept obtained from the regression were used to generate a scaled 405 nm control signal. This fitted control signal was then smoothed using a moving average filter (MeanFilter) with an order of 1000 to reduce noise. The ΔF/F was calculated by normalizing the 465 nm signal against the smoothed control signal, subtracting the fitted 405 nm signal from the 465 nm signal, and dividing by the fitted 405 nm signal. The resulting normalized signal was expressed as a percentage to represent the change in fluorescence relative to baseline levels.

### Locus coeruleus optogenetics

To induce LC excitation, two-second stimulations consisting of 10 ms pulses at 20 Hz (resulting in a 40 ms break between pulses) at a 5-mW intensity were applied bilaterally to the LC using a 430-490 nm laser (Shanghai Laser & Optics Century Co., Ltd., model: BL473-200FC). A two-hour baseline without stimulation was implemented before and after the five-hour stimulation paradigm. The simulations were conducted in a closed loop, where stimulation was triggered if the smoothed, real-time calculated NE ΔF/F (%) crossed the threshold based on an average ΔF/F (%) value of the last two minutes. The threshold was initially set to −15 ΔF/F (%) but increased by 5 every hour until reaching the final level of +5 ΔF/F (%). Throughout the baseline and stimulation paradigm, fiber photometry recordings from the mPFC, along with EMG and EEG, were acquired. Recordings took place during the light phase to encourage natural sleep behavior.

To induce LC inhibition, two-minute Arch stimulations at 5 mW intensity, followed by a four-minute break, were applied bilaterally to the LC using a 532 nm laser (Changchun New Industries Optoelectronics Tech. Co., Ltd., model: 16030476). Simultaneously, GRAB-NE2m(3.1) recordings were made at the mPFC site, along with the previously described EEG and EMG measurements. The two-hour recordings took place during the light phase when the mice were expected to exhibit natural sleep/wake behavior undisturbed. The mice were recorded with an infrared camera.

### Novel object recognition task

The NOR task was performed on 12-20 weeks old mice involved in the LC suppression experiments in an open-field arena (PVC, custom-made, 40 x 42 x 40 cm). The mice were habituated to the environment for 10 minutes as part of the encoding phase, where the mice familiarized themselves with two objects. Following this, the animals underwent the LC suppression paradigm and were then given five minutes in the NOR task arena for the recall phase to determine the novel-to-familar ratio. In the recall phase, one of the two known objects had been exchanged for an unknown object and the ratio of approaches to the novel vs. familiar object was recorded. Natural sleep was encouraged between the encoding and recall phases by performing the task during the light phase in an environment where the mice had been habituated for over 12 hours prior to the experiment. The mice were tracked using Ethovision XT 11.5 (Noldus).

### EMG signal preprocessing

The R-peak detection algorithm was applied to EMG recordings, that were priorly subjected to filtration in frequency domain using in-house pipeline in MATLAB. First, all the signals were subjected to removal of frequencies associated with amplitude current using a bandstop infinite impulse response (IRR) Butterworth 4^th^ order filter between frequencies of 59 and 61 Hz. In case of recordings acquired in US, the cut-off frequencies for the notch filter were of 60 to 64 Hz, and the filter order of 2 was used due to different frequency span of the nuisance signal. Subsequently, a high-pass IIR Butterworth filter with 0.6 steepness was applied to the frequencies <2.5Hz to gradually remove fluctuation of the EMG signal baseline from the recordings. This was followed with low-pass filtration >80Hz, using the same filter but of 0.9 steepness. The steepness’s of the filters were chosen to minimally induce artefacts at the border conditions, and either gradually (high-pass) or rapidly (low-pass) remove the signals that affect detecting R peaks due to either increased fluctuations of the baseline (i.e., insufficient grounding or slow motion) or high frequency noise (relatively high spectral power of muscle signals). In a few recordings, both low-pass and high-pass frequencies were adapted to be slightly above 4Hz (but not higher than 5 Hz) and 45Hz respectively, to better remove the signal baseline fluctuations or higher amplitude high-frequencies due to presence of artefacts significantly affecting the R peaks detection. This was, however, applied in recordings of low signal-to-noise ratio in the frequency range of HR signals, and was carefully evaluated for appropriateness in all cases, so that the bandwidth of the signals within the range of heart pulsations was preserved. For that purpose, at each step of the signal preprocessing, periodograms of the filtered and unfiltered signals were computed and compared to evaluate the effects of the applied filtrations. After filtration, all signals were saved in the .mat format and were used for further R peaks detection.

### Heart rate detection

R-peak detection in mice was based on preprocessed EMG recordings to obtain HR without needing additionally invasive procedures. EMG is acquired in most sleep studies to aid sleep scoring, and the method has previously been validated in Lecci et al. (Lecci et al., 2017) and subsequently used for other publications (Osorio-Forero et al., 2025, 2021). User settings for this function included adjusting the movement threshold and peak prominence to ensure optimal detection for different traces. The MATLAB function first determines the movement threshold based on the overall mean and SD of the trace. It then runs peak detection in windows of two seconds of data at a time. The threshold for the peaks is based on the mean and SD within each window to ensure peak detection even in cases of power loss. If movement is detected within a section, data that falls 100 ms before and 250 ms after the movement is not considered for peak detection due to noise and is also excluded from the peak threshold calculation within the window.

When all windows are analyzed, the timestamps at which R-peaks have been found are processed through a diff function to determine the time between peaks. The original time vector is maintained to ensure that the times of the RR are preserved for event extraction. To reduce any artifacts in the data, RR that suggested a HR faster than 800 BPM or slower than 400 BPM were removed, as previous findings suggest that the HR typically falls within 450-650 BPM throughout a 24-hour cycle (D’Souza et al., 2021; Ho et al., 2011; Sakata et al., 2005). Lastly, the data and associated time vector were interpolated to 64 Hz using the p-chip method, as in Chen et al. (Chen et al., 2023), since this method was found to have the lowest error when analyzing the frequency domain (Benchekroun et al., 2023). Interpolation was performed to obtain a consistent sampling rate, but also to get RR interval estimates during periods of movement based on the values leading up to and after the movement periods.

### EEG power

For the mouse data, the power within each EEG band was determined by applying the spectrogram function to extract the short-time Fourier transformation of the EEG in one-second windows for frequencies between 0 and 100 Hz. To obtain frequency power bands, the absolute log of the short-time Fourier-transformed data was taken, and the mean values within each frequency band (slow oscillations: 0.5 - 1 Hz, delta: 1 - 4 Hz, theta: 4 - 8 Hz, sigma: 8 - 15 Hz, beta: 15 - 30 Hz, gamma (high): 80 - 100 Hz) were extracted. The sigma band was defined as 8 - 15 Hz to maintain consistency with our previous publications. Modest differences in sigma-band boundaries are not expected to affect the main conclusions.

### Mouse sleep scoring

The mice were recorded in chambers using ViewPoint Behavior Technology, with EEG and EMG electrodes connected to a commutator (Plastics One, Bilaney, SL12C) through cables. Habituation in the recording environment lasted at least 12 hours before recordings, during which the EEG and EMG cables were connected. The mice were attached to fiber optic implants immediately before the recording, which lasted for 2–9 hours during their light phase. The EEG and EMG signals were amplified using a model 3500, 16-channel AC amplifier from National Instruments. A bandpass filter was applied to the EEG at 1-100 Hz and to the EMG at 10-100 Hz. Both signals had an added notch filter at 50 Hz to reduce power line noise. The EMG and EEG were digitized through a Multifunction I/O DAQ device from National Instruments (USB-6343) and sampled at 512 Hz. An infrared camera from FLIR Systems was used to record the mice to aid in scoring vigilance states.

Sleep scoring for mice was performed manually through visual inspection of the EEG and EMG traces, along with the video, to divide the traces into 5-second and, later, 1-second windows. Scoring was performed using SleepScore software from ViewPoint Behavior Technology. Vigilance states included ’wake’ (high-amplitude EMG and high-frequency, low-amplitude EEG), ’NREM sleep’ (low EMG amplitude and low-frequency, high-amplitude EEG), and ’REM sleep’ (low EMG amplitude and high-frequency, low-amplitude EEG). Later in the analysis pipeline, wake periods between 5-15 s were classified as long MAs, and wake periods below 5 s were scored as short microarousals.

### Event marker selection

All time-domain analyses were based on the timestamps of event markers in the data. To determine the NE trough timestamps during the sleep transition analysis, similar approaches were used for the different stages. Across all stages, the NE trace was inverted, and peak detection was performed on the data. For the NE troughs during NREM, a peak prominence criterion of 0.7 to 3 was set. For NREM to MA (short and long), only data from 30 s prior to the transition were considered, and the peak prominence criterion was set to 0.8 to 1.8. Additionally, there had to be at least 15 s between troughs, with the one closest to the transition being favored. Lastly, for transitions to wake, data from 15 s prior to the scored transitions were considered. The peak prominence criterion was set to 0.1 to 0.4, and the peak height was set to a minimum of the mean of the trace minus five. After automatic trough detection, manual verification was performed across all transitions to ensure optimal trough detection.

The event marker for the data presented in Fig. 2 and 3a-b was the onset of the stimulation burst. As the laser had an online measure of NE and turned on for 2 s at 20 Hz, the event marker was defined as the point when the laser turned on after being off for over 100 ms. These timestamps were divided into groups based on when the threshold shifted. For the rest of Fig. 3, the RR peak value 2-7 s after onset of stimulation burst was utilized as the event marker for HR decelerations. The event marker for the data presented in Fig. 4 was set to when the laser turned on. For both LC excitation data, the exclusion criteria required that the animal be in NREM for 30 s before and after the laser turning on to be included and for LC suppression that the animal was in NREM for 30 s before and 60 s after laser onset.

For figure 5, the event marker was HRBs. These events have previously been reported in humans, where they are defined as when the RR intervals drop below 2 SD of the mean in a 3 min window (Chen et al., 2020; Naji et al., 2019a). This event marker was for mice adjusted to be 2.2 SD from the mean in 24 s windows. This adjustment was done based on differences in dynamics. The time window was adjusted based on the differences in mean HR between humans and mice.

### Data analysis

Fiber photometry, EMG, and EEG data were extracted from all mice, and RR was calculated for each recording using the R-peak detection algorithm. For analysis pertaining to the sleep transition data, the EEG data were excluded from further analysis for two animals in the sleep transitions analysis due to noise, resulting in 245 events for NREM to NREM transitions (191 with EEG), 174 events for NREM to short MA transitions (133 with EEG), 75 events for NREM to long MAs (54 with EEG), and 66 events for NREM to wake transitions (42 with EEG) across the seven mice.

For the LC activation-related analysis, 108 events were found for Threshold -15 (11 of these being YFP), 260 events for Threshold -10 (55 of these YFP), 777 events for Threshold -5 (296 of these YFP), 1,444 events for Threshold 0 (510 of these YFP), and 1,148 events for Threshold 5 (377 of these YFP) across ten animals (four being YFP). For the LC suppression experiment, 48 events were found for the four Arch mice and 38 for the four YFP mice. All data preprocessing was performed using MATLAB. For the mouse data related to HRB events, 1654 events were found across the 7 mice also analyzed in connection with sleep transitions.

For all four experiments, the event variables were generated, and data from RR, NE, and the EEG frequency bands were extracted from 60 s before and after each event. Each data type was averaged across all events from all mice within each relevant group for each experiment, leading to the standard error being based on the number of events rather than the number of animals.

Across all experiments and data types, the AUC was calculated as a measure of change between the different conditions. For the sleep transitions experiments, the AUC from -25 to 0 s (zero being the event time) was used as a baseline and was subtracted from the AUC from 0 to 25 s after the event. For the ChR2/YFP experiment, three types of AUC measures were used: the pre-stimulation AUC was defined as the data from -40 to -20 s before the event (baseline) subtracted from the data from -20 to 0 s before the event. Similarly, the post-stimulation AUC was defined as the data from 20 to 40 s before the event (baseline) subtracted from the data from 0 to 20 s after the event. The pre-stimulation minus the post-stimulation AUC was defined as the difference between the two measures outlined above. Lastly, the AUC for the Arch/YFP-related experiments were defined as -40 to -5 s before the event marker (baseline) subtracted from 5 to 40 s after the event marker.

The data presented in Fig. 3 was based on the same +/- 60 s data files surrounding the LC stimulation onset as seen in Fig. 2, though only ChR2 animals were included in this analysis, as it focused on HR changes in response to stimulation. In Fig. 3a, mean traces across thresholds were shown and 97 raw traces were randomly extracted for each threshold to visualize variability, as this was the lowest number of total events present within a threshold. In Fig 3b, the RR, BPM and RR SD estimate was based on the full 2 min data file, while the sample entropy (m = 2) only included the 10 s after laser onset (Richman et al., 2004). The following analysis was based on the HR deceleration event marker. Here, the RR amplitude was calculated as the RR at HR decelerations minus the mean RR 8-10 s prior. Similarly, the NE amplitude was determined by subtracting the lowest NE value 5-10 s before the HR decelerations from the NE value at the HR decelerations. The mean NE values also reflected the NE values in the 10 s leading up to the HR decelerations. Lastly, the sigma amplitude was determined as the mean sigma 2-3 s before the HR decelerations subtracted from the sigma value at the HR decelerations.

For the analysis related to HRB events, amplitudes were analyzed rather than AUC to use similar measures as in the human data. For the mouse data, the RR amplitude was defined as the RR value at the HRB minus the mean across the 15 to 10 s leading up to the event. The NE amplitude was calculated as the peak value in the 2.5 s interval around the HRB event minus the minimum in the 15 to 10 s interval preceding the event. Lastly, the sigma amplitude was defined as the smallest observation in the 2.5 s interval around the HRB event minus the largest value found between 15 to 5 s range preceding the event.

The frequency-domain analysis was based on the non-parametric Welch approach, and similar procedures were used on NREM data across all experiments. The data underwent detrending with a 5th degree polynomial fitted to the raw data before analysis as in Ferreira et al. (Ferreira et al., 2017). Across experiments, frequencies between 0-0.1 Hz with a frequency resolution of 0.0002 Hz were analyzed for NE, and frequencies between 0-0.4 Hz with a resolution of 0.002 Hz were calculated for RR. Across all frequency-domain analyses, consecutive periods of minimally 300 s of NREM sleep (with MAs) were included for RR/NE, as this was the minimal NREM period duration suggested to capture VLF (Shaffer and Ginsberg, 2017). However, as the ability to interpret VLF power within short term recordings has previously been questioned (Thayer et al., 2022), we also extracted NREM sleep periods of minimum 11 minutes (allowing 10 cycles of VLF 0-0.015 Hz) to visualize an overlap in the frequency distribution (Supplementary Fig. 1f). Given the large similarities between the frequency distributions, we chose to utilize the minimum of 300 s NREM as the data selection criteria, as it is uncommon for mice to remain in NREM sleep for over 10 min, making frequency-domain analyses across different experimental paradigms challenging. Each bout of data was run through Welch separately, and its AUC, peak power value, and frequency at peak power were extracted along with information about the length of the period (bout length). Bout length was subsequently used to weigh the final mean/SEM estimates, which was utilized both for statistical analysis and visualization generation. SEM was based on the number of bouts. The PSD presented in Fig. 1 and Supplementary Fig. 1 were based on the same data from the mice used to analyze sleep transitions and held 144 bouts. For the LC activation data, the PSD was made separately for ChR2/YFP and for each stimulation threshold including the baseline before and after stimulation. The number of bouts for the ChR2 data (from baseline to threshold 5 were 212, 44, 41, 30, 37, and 43. For YFP, these numbers were 147, 24, 28, 21, 27, and 28. For the LC suppression data, the PSD was split by Arch/YFP, which consisted of 66 and 43 bouts respectively.

### Statistical analysis

Linear mixed-effects models (LMMs) were utilized across all figures to estimate the effects using R. One exception was made for models related to memory estimates, as only one estimate was available per night per subject. Thus, a standard linear model was employed instead. The choice of applying LMMs to the data was made, as the model type allowed for event-based analysis by adding a random intercept for subject ID. By doing so, a hierarchy was created in the data to ensure that issues with independence did not occur (Yu et al., 2022). When continuous predictors were present in the model, the predictor was added as a random slope and compared to the base model (with only a random intercept) to determine which model was best using the Akaike and Bayesian information criteria. If the random slope model had a lower Akaike/Bayesian information criterion, converged, and did not have a singular fit, the statistics were summarized based on that model.

Models correlating sleep transitions with an array of dependent variables were built by setting the four sleep transitions as independent variable factors, which were re-leveled to ensure that the lowest level of arousal (NREM to NREM) was set as the baseline. For models associated with Fig. 2-3, the thresholds were similarly set as a factor, with -15 as the baseline threshold, except for the frequency-domain analysis, where the baseline was set as the intercept. The thresholds and conditions (ChR2 or YFP) were both set as predictors in Fig. 2 and had an interaction effect to explore the differences between the two conditions at different thresholds. In Fig. 3, the thresholds were the main predictor utilized, as only ChR2 data was included. Lastly, models for Fig. 4 had a similar predictor setup with condition (YFP or Arch) as the predictor. All LMMs had SubjectID as a random intercept.

After fitting the LMMs, they were assessed for any violations of the fixed effects assumptions through visual inspection of the residual distributions. While LMMs have been found to be robust to assumption violations (Schielzeth et al., 2020), bootstrapping of 1000 samples was performed to obtain robust estimates of the model in cases where one or more assumptions had been violated. For models where autocorrelation of residuals was present, block bootstrapping was chosen over normal bootstrapping. The block bootstrapping resampled the data in groups based on subject ID, which was the recommended solution to issues with autocorrelations of residuals. Additionally, when relevant to the analysis, weights were added to the LMMs and factored into the bootstrapping to ensure the bout length was appropriately weighed in the model.

For models with a continuous independent variable, the marginal and conditional R squared values were reported in the figure and was also the measure that was bootstrapped (Efron, 1983; Nakagawa and Schielzeth, 2012). In cases where the predictor had more than two levels or there were multiple predictors, the estimates of the pairwise comparisons were bootstrapped. Lastly, in cases where only a single predictor with two levels was present, the difference between model estimates for the two levels was bootstrapped. All pairwise comparisons were adjusted for multiple comparisons using Turkey’s method.

### Participants

28 human participants (Mage=20.46 ± 2.73 years, 17 females) were included in the analysis of HRB. For information regarding demographics, see the supplementary table. Participants had no history of psychological, neurological or chronic illnesses and were previously included in another publication (Chen et al., 2021). All participants provided informed consent approved by both the Western Institutional Review Board and the Human Research Review Board at the University of California, Riverside. Individuals were excluded from participation if they exhibited irregular sleep/wake cycles, had diagnosed sleep disorders, personal or familial histories of psychopathology, substance abuse or dependence, a history of epilepsy or loss of consciousness exceeding two minutes, current psychotropic medication use, or cardiac or respiratory illnesses potentially influencing cerebral metabolism. Eligibility was determined during an in-person psychiatric assessment conducted by trained research personnel. Additionally, participants underwent a medical evaluation, including a detailed medical history review and physical examination by a staff physician, to confirm overall physical health. Participants were instructed to avoid caffeine, alcohol, and all stimulants for at least 24 hours prior to and during the experimental day. Compliance with consistent sleep patterns for one week prior to each experimental session was verified using sleep diaries completed daily and wrist-based activity monitors (Actiwatch Spectrum, Philips Respironics). Participants were required to achieve a minimum of 7 hours of sleep per night during this period, including the night immediately before the study day. Monetary compensation and/or course credits were provided to participants in acknowledgment of their time and participation.

### Word-paired association task

Human participants underwent the word-paired association task, which consisted of an encoding, criterion and testing phase. In the encoding phase, participants were presented with 60 word-pairs one at a time for 1 s with an inter-trial interval of 1s. During the criterion phase, the participants were probed with one of the words from a pair and were subsequently presented with the missing word for 2 s followed by a 100 ms inter-trial interval. This process continued until the participants obtained 70% accuracy overall. Both the encoding and criterion phase occurred in the morning on the first day. Lastly, the testing phase was conducted three times: shortly after the criterion phase, in the evening before participants slept in the lab and the following morning. At each test, participants were tested on their memory for 20 of the half word-pairs. Participants were shown one word and asked to recall the missing word. No feedback was provided during the testing phase and there was a 1 s inter-trial interval between pairs. Performance was measured as the difference score between the first test and the final test approximately 24 hours later.

### Heart rate detection in humans

In humans, ECG data were recorded using a modified Lead II Einthoven configuration at a sampling rate of 1,000 Hz. Heart rate variability (HRV) analysis was conducted on the R-wave intervals across the entire sleep-wake period using Kubios HRV Analysis Software 2.2 (Biosignal Analysis and Medical Imaging Group, University of Kuopio), following Task Force guidelines(“Heart rate variability: Standards of measurement, physiological interpretation, and clinical use,” 1996). The R-wave peaks within the ECG QRS complex were automatically detected by Kubios software and subsequently verified visually by trained technicians. Any incorrectly identified R-peaks were manually adjusted, and missing beats were interpolated using cubic spline interpolation. After computing inter-beat intervals, a third-order polynomial detrending filter was applied to eliminate trend-related components in the data. Artifacts were further corrected using Kubios software’s built-in automatic medium-level artifact correction filter.

### Human sleep recordings

EEG data were collected using a 32-channel electrode cap (EASEYCAP GmbH) equipped with Ag/AgCl electrodes arranged according to the international 10-20 system (Jasper, 1958). Among these electrodes, 22 were utilized for scalp EEG recordings, while the remaining electrodes were dedicated to ECG, electromyogram (EMG), electrooculogram (EOG), ground, an online common reference channel located at FCz (retained post-referencing), and mastoid references (A1 and A2). EEG signals were recorded at a sampling rate of 1,000 Hz and subsequently referenced offline to the contralateral mastoid electrode (A1 and A2).

Data preprocessing was performed using BrainVision Analyzer 2.0 software (BrainProducts). Eight scalp electrodes (F3, F4, C3, C4, P3, P4, O1, O2), along with EMG and EOG recordings, were selected for scoring nighttime sleep stages. EEG and EOG signals were filtered with high-pass and low-pass settings of 0.3 Hz and 35 Hz, respectively. Raw data were visually staged in 30-second epochs into Wake, Stage 1, Stage 2, Slow Wave Sleep (SWS), and REM sleep, according to the criteria of Rechtschaffen and Kales, using the custom MATLAB-based scoring toolbox, HUME. Following staging, epochs containing artifacts and arousals were visually identified and removed prior to spectral analysis.

### Human data analysis

HRBs were used as the event marker for human data and was defined as when the RR intervals drop below 2 SD of the mean in a 3 min window (Chen et al., 2020; Naji et al., 2019a). This resulted in 4753 events across the 28 subjects during stage 2 NREM sleep. For these events, the RRRecovery amplitude reflected the mean RR values 5 to 10 s after the HRB minus the RR value at the event-time. The sigma power (12-15 Hz) trace was extracted and its value in the 5 to 0 s preceding the event was used to correlate to the RRRecovery. To correlate the sigma values to memory estimates, we followed the same procedure as past papers(Naji et al., 2019a). Sigma power was extracted in +/- 20 s around the HRB and then segmented into four-time bins: 10 to 5 s preceding the event, 5 to 0 s preceding the event, 0 to 5 s following the event, and 5 to 10 s following the event. For each participant, sigma power was averaged across all events within each bin, resulting in one mean value per bin. To correlate sigma power before the HRB and memory performance, baseline sigma power was subtracted from the sigma power 5 s before the HRB, following the equation below. This baseline represents mean sigma power during NREM sleep outside of 20 s windows surrounding event extracted data. All statistical tests followed the same principles outlines in the ‘Statistical Analysis’ section.

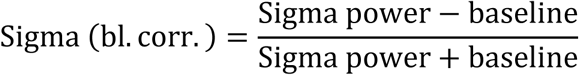

## List of Supplementary Materials

### Supplementary figures

Supplementary figure 1. Heartbeat extraction from EMG recordings during sleep (belonging to Figure 1)

Supplementary figure 2. Electroencephalography responses to locus coeruleus stimulation (belonging to Figure 2)

Supplementary figure 3. YFP control for closed-loop optogenetic activation of locus coeruleus (belonging to Figure 2)

### Supplementary table

Supplementary table: Demographics of human subjects

### Statistics

Figure 1: Sleep Transitions (belonging to Figure 1)

Figure 2: Locus coeruleus stimulation (belonging to Figure 2)

Figure 3: HR decelerations (belonging to Figure 3)

Figure 4: Locus coeruleus suppression (belonging to Figure 4)

Figure 5: Heart rate bursts (belonging to Figure 5)

Supplementary Figure 2: YFP for Figure 2 (belonging to Supplementary Figure 2) Supplementary Figure 3: EEG for Figure 2 (belonging to Supplementary Figure 3)

## Acknowledgements and Funding

Rodent work was supported by the Lundbeck Foundation Fellowship (R413-2022-622), Lundbeck Foundation Seed grant (R471-2024-458) and Independent Research Fund Denmark (2100-00018B). Work with human participants was supported by the Office of Naval Research (grant N00014-14–1-0513) and the National Institutes of Health (NIH; grant R01 AG046646). Marie Skłodowska-Curie Postdoctoral Fellowship (HORIZON-MSCA-2022-PF-01-01; SEP-210878699) was awarded to P.C.

## Author contributions

Conceptualization: C.K., S.M., M.N., S.J.. Data collection: M.A. and P.C.. Database handling: S.J. and A.M.. Visualization: C.K., S.J., A.M.. Data preprocessing: S.J., R.G., A.M.. Data analysis: S.J.. Supervision: C.K, S.M.. Writing—original draft: C.K. and S.J.. Writing—review and editing: C.K., S.J., R.G., S.M., A.M., P.C., M.N, M.A..

## Data availability

The human dataset is available from: https://osf.io/g6emj/. Mouse datasets are available upon reasonable request from the corresponding authors.

## Code availability

Code for this project is available on Github under https://github.com/sofie-jac/HR-NE-fluctuations

