## Supplementary File for "Noradrenergic infraslow rhythm during sleep is the critical link between heart-rate dynamics and memory consolidation"

### Supplementary Materials

**Supplementary figures:**

Supplementary figure 1 (belonging to Figure 1)

Supplementary figure 2 (belonging to Figure 2)

Supplementary figure 3 (belonging to Figure 2)

**Supplementary table:**

Supplementary table: Demographics of human subjects

**Statistics:**

Figure 1: Sleep Transitions (belonging to Figure 1)

Figure 2: Locus coeruleus stimulation (belonging to Figure 2)

Figure 3: HR decelerations (belonging to Figure 3)

Figure 4: Locus coeruleus suppression (belonging to Figure 4)

Figure 5: Heart rate bursts (belonging to Figure 5)

Supplementary Figure 2: YFP for Figure 2 (belonging to Supplementary Figure 2)

Supplementary Figure 3: EEG for Figure 2 (belonging to Supplementary Figure 3)

#### **Supplementary Figures**

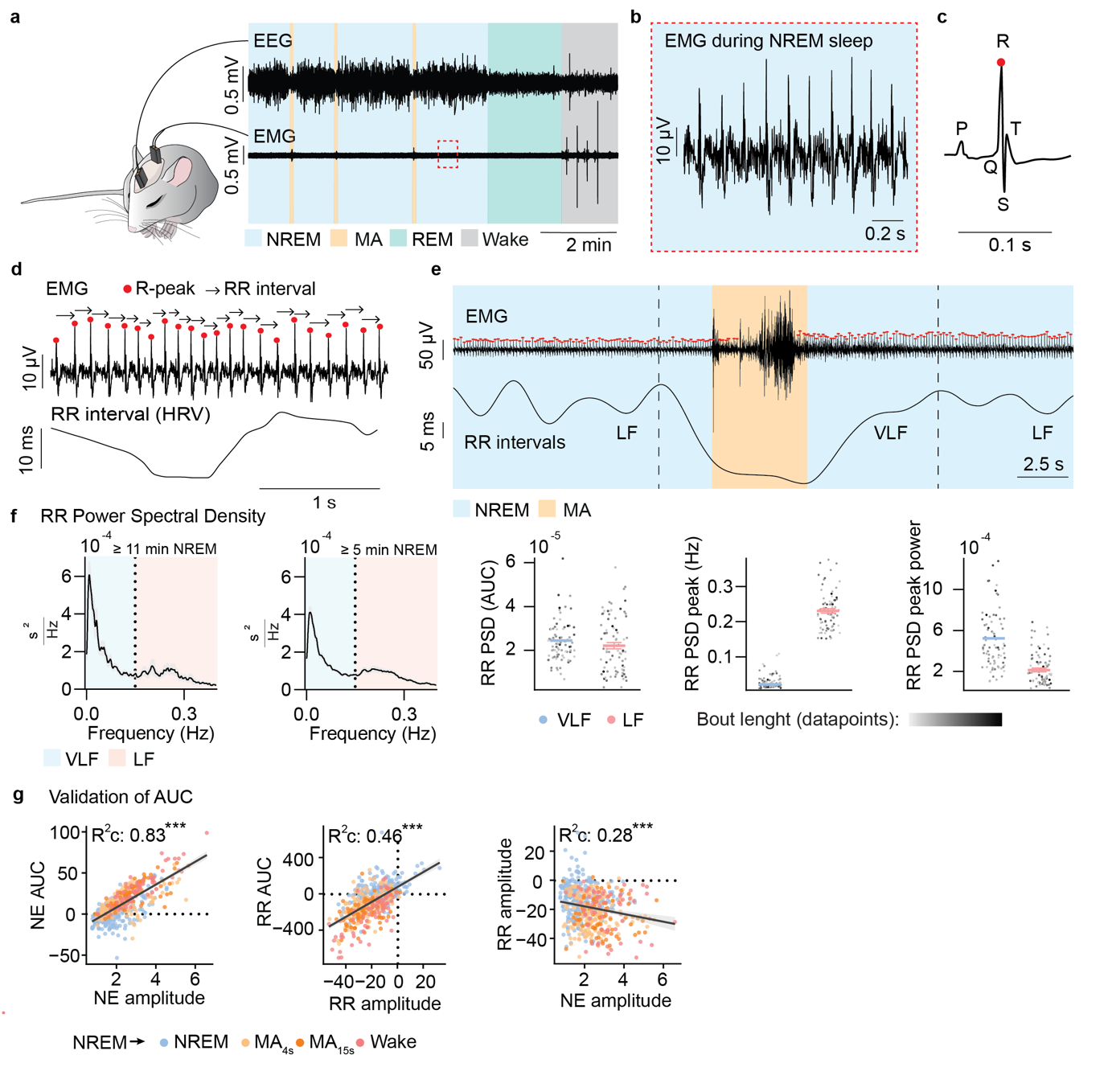
**Supplementary figure 1: Heartbeat extraction from EMG recordings during sleep**. **a.** Example trace of the EEG and EMG with the area selected for further inspection (NREM = blue shade, microarousal (MA) = yellow shade, REM = green shade, wake = gray shade). **b.** Area highlighted in example trace. **c.** QRS complex with the R-peak highlighted in red. **d.** Example of detected R-peaks from an EMG trace (top) and the calculated RR intervals (RR) from the trace (bottom). **e.** Example trace of EMG and RR (NREM = blue shade, MA = yellow shade). RR ranged from values 0.1 to 0.139 (432-600 BPM). **f.** From left to right: 1) RR power spectral density (PSD) across the very low frequency (VLF, blue shade), low frequency (LF, orange shade) based on minimally 11 min long NREM periods. 2) RR PSD based on minimally 5 min long NREM periods. All the following sub-figures will be based on this frequency distribution. 3) Area under the curve for the VLF and LF. 4) Frequency at the highest point of the PSD for VLF and LF. 5) Power value at the highest point of the PSD for the VLF and LF. Datapoint darkness is proportional to bout length (n=7, bouts=144). Data shown is mean ± standard error of the mean (SEM). All frequency domain analysis figures have visualizations based on weighted estimates (see methods). **g.** Validation of area under the curve (AUC) measurements for NE and RR intervals.

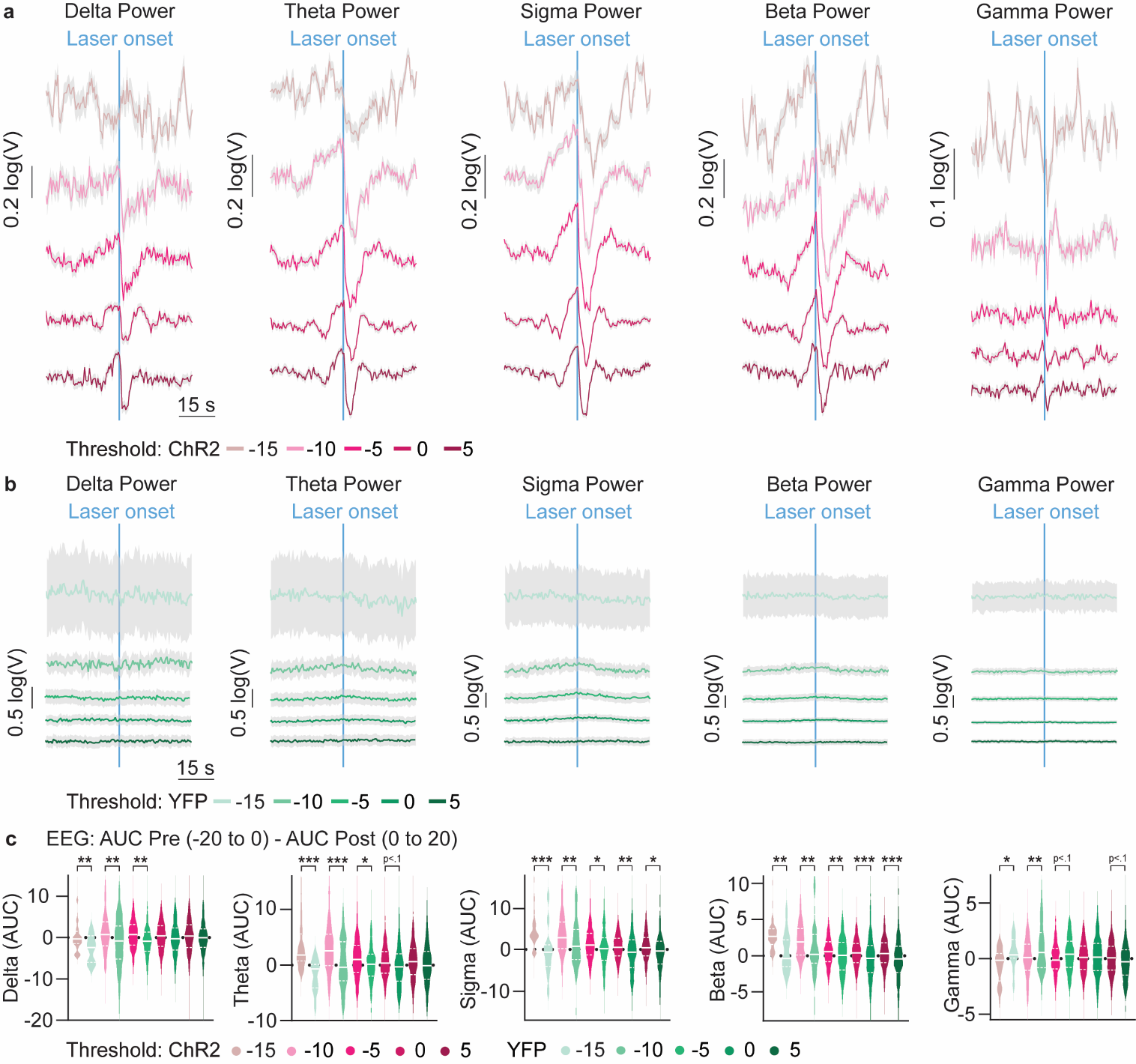
**Supplementary figure 2**: **Electroencephalography responses to locus coeruleus stimulation**. **a.** Mean traces with standard error of the mean (SEM) of (from left to right) delta, theta, sigma, beta, and gamma power in ChR2 animals surrounding the onset of a laser burst for all five thresholds: -15 (beige), -10 (light pink), -5 (pink), 0 (dark pink), and 5 (wine red). **b.** Mean traces with SEM of (from left to right) delta, theta, sigma, beta, and gamma power in YFP animals surrounding the onset of a laser burst for all five thresholds: -15 (pastel green), -10 (light green), -5 (green), 0 (emerald), and 5 (dark green). **c.** Pre-stimulation area under the curve (AUC) minus the post-stimulation AUC from ChR2 and YFP animals for (from left to right) delta, theta, sigma, beta, and gamma power across all thresholds: ChR2 -15 (beige), ChR2 -10 (light pink), ChR2 -5 (pink), ChR2 0 (dark pink), ChR2 5 (wine red), YFP -15 (pastel green), YFP -10 (light green), YFP -5 (green), YFP 0 (emerald), and YFP 5 (dark green). *p < 0.05., **p < 0.01., and ***p < 0.001. For a more detailed overview of the statistics, see section Supplementary figure 3 in ‘Statistics’ under Supplementary Materials.

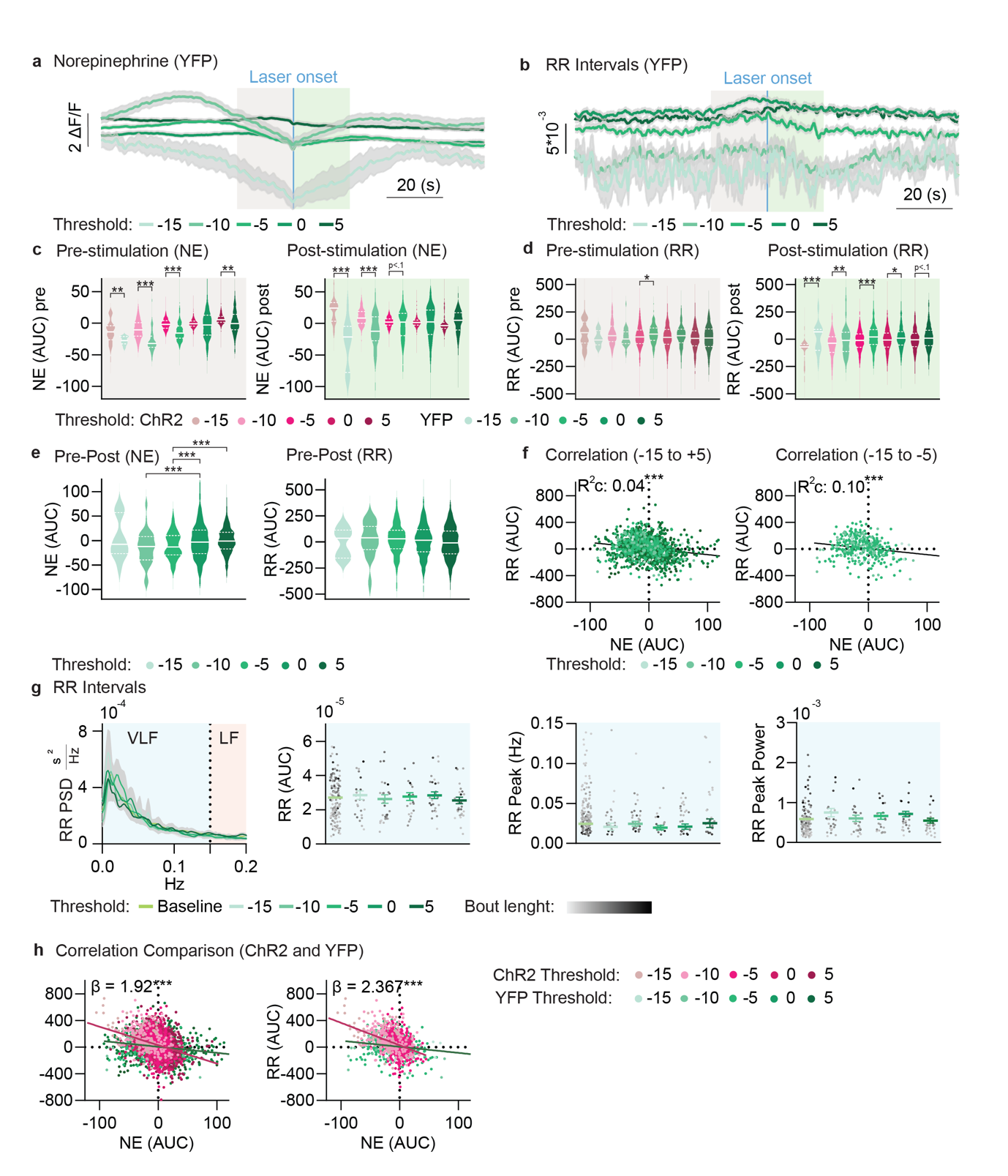

**Supplementary figure 3: YFP control for closed-loop optogenetic activation of locus coeruleus**. **a.-b.** Mean traces with standard error of the mean (SEM) of YFP norepinephrine (NE, left) and YFP RR intervals (RR, right) surrounding the onset of a laser burst with beige (pre-stimulation) and green (post-stimulation) background shade for all five thresholds: YFP -15 (pastel green), YFP -10 (light green), YFP -5 (green), YFP 0 (emerald), and YFP 5 (dark green). **c.-d.** Comparisons of area under the curve (AUC) of NE (**c.**) and RR (**d.**) between ChR2 and YFP animals either 20-0 s before stimulation (left) or 0-20 s after stimulation (right) for all five thresholds: ChR2 -15 (beige), ChR2 -10 (light pink), ChR2 -5 (pink), ChR2 0 (dark pink), ChR2 5 (wine red), YFP -15 (pastel green), YFP -10 (light green), YFP -5 (green), YFP 0 (emerald), and YFP 5 (dark green). **e.** Pre-stimulation AUC minus the post-stimulation AUC for NE (left) and RR (right) for YFP animals at all thresholds: -15 (pastel green), -10 (light green), -5 (green), 0 (emerald), and 5 (dark green). **f.** Left: correlation of the pre-stimulation AUC minus the post-stimulation AUC for NE and RR for YFP animals across all thresholds (-15 (pastel green), -10 (light green), -5 (green), 0 (emerald), and 5 (dark green)). Right: correlation of the pre-stimulation AUC minus the post-stimulation AUC for NE and RR for YFP animals across thresholds -15, -10, and -5. **g.** From left to right: 1) PSD for RR between 0-0.2 Hz across all thresholds: Baseline (lime), -15 (pastel green), -10 (light green), -5 (green), 0 (emerald), and 5 (dark green). 2) AUC of the RR PSD across all thresholds in the very low frequency (VLF) range. 3) Frequency at the peak of the PSD distribution across thresholds for RR in the VLF range. 4) Power value at the peak of the PSD across thresholds for RR in the VLF range. Datapoint darkness is proportional to bout length. Number of YFP bouts for frequency domain analysis were, from baseline to higest threshold: 147, 24, 28, 21, 27, and 28. *p < 0.05., **p < 0.01., and ***p < 0.001. All frequency domain analysis figures have visualizations based on weighted estimates (see methods). For a more detailed overview of the statistics, see section 3e-i & Supplementary figure 2 in ‘Statistics’ under Supplementary Materials. **h.** Left: correlation comparison between ChR2 and YFP across all thresholds. Right: correlation comparison between ChR2 and YFP across thresholds -15, -10, and -5.

#### **Supplementary Table**

**Supplementary table**: Human demographics.

| **Ethnicity** | **Female** | **Male** | **Total** |
| --- | --- | --- | --- |
| Black | 0 | 1 | 1 |
| Asian | 5 | 3 | 8 |
| White | 6 | 3 | 9 |
| Hispanic | 5 | 4 | 9 |
| Mixed Race | 1 | 0 | 1 |
| **All** | 17 | 11 | 28 |

#### **Statistics**

##### Figure 1: Sleep Transitions

###### 1d: NE predicted by Sleep Stage transitions

The analysis revealed significant effects of Sleep Transition on NE (AUC). Specifically, compared to the reference category (NREM), all other levels showed significant increases in NE. Pairwise comparisons confirmed these differences (all p < .05). Given the slight violation of the normality of residuals assumption violations and the presence of heteroscedasticity, bootstrapping was used to provide robust CIs, which supported the primary findings (see supplementary materials).

**Fixed Effects of Sleep Stage Transitions**

| **Predictor** | **Estimate** | **Std. Error** | **df** | **t-value** | **p-value** |
| --- | --- | --- | --- | --- | --- |
| (Intercept) | 1.60 | 1.61 | 9.31 | 0.994 | 0.345 |
| NREM | 0 | - | - | - | - |
| NREM before MA (short) | 14.82 | 1.46 | 555.91 | 10.15 | < .001*** |
| NREM before MA (long) | 22.57 | 1.95 | 555.14 | 11.56 | < .001*** |
| NREM before Wake | 28.98 | 2.10 | 542.46 | 13.82 | < .001*** |

**Pairwise Comparisons with Tukey Adjustment**

| Contrast | Estimate | SE | df | t-ratio | p-value | Lower Bound (2.5%) | Upper Bound (97.5%) |
| --- | --- | --- | --- | --- | --- | --- | --- |
| NREM - NREM before MA (short) | -14.82 | 1.47 | 556 | -10.11 | < .0001*** | -17.36 | -12.15 |
| NREM - NREM before MA (long) | -22.57 | 1.96 | 555 | -11.50 | < .0001*** | -26.56 | -18.89 |
| NREM - NREM before Wake | -28.98 | 2.12 | 543 | -13.70 | < .0001*** | -34.46 | -23.89 |
| NREM before MA (short) - NREM before MA (long) | -7.75 | 2.02 | 556 | -3.85 | 0.0008*** | -11.57 | -4.10 |
| NREM before MA (short) - NREM before Wake | -14.17 | 2.14 | 554 | -6.61 | < .0001*** | -19.51 | -9.36 |
| NREM before MA (long) - NREM before Wake | -6.41 | 2.45 | 554 | -2.62 | 0.0443* | -12.11 | -1.18 |

###### 1e: RR predicted by Sleep Stage transitions

The analysis revealed significant effects of Sleep Stage Transition on RR (AUC). Specifically, compared to the reference category (NREM), all other levels showed significant decreases in RR. Pairwise comparisons confirmed these differences (all p<.05), except for the comparison between NREM to MA (long) and NREM to Wake, where the evidence was non-significant. Given the presence of heteroscedasticity, bootstrapping was used to provide robust CIs, which supported the primary findings.

**Fixed Effects of Sleep Stage Transitions**

| **Predictor** | **Estimate** | **Std. Error** | **df** | **t-value** | **p-value** |
| --- | --- | --- | --- | --- | --- |
| (Intercept) | 15.941 | 18.722 | 7.968 | 0.851 | 0.419 |
| NREM | 0 | - | - | - | - |
| NREM to MA (short) | -146.200 | 13.762 | 555.317 | -10.624 | < .001*** |
| NREM to MA (long) | -203.970 | 18.418 | 555.797 | -11.074 | < .001*** |
| NREM to Wake | -245.994 | 19.830 | 554.361 | -12.405 | < .001*** |

**Pairwise Comparisons with Tukey Adjustment**

| **Contrast** | **Estimate** | **SE** | **df** | **t-ratio** | **p-value** | **Lower Bound (2.5%)** | **Upper Bound (97.5%)** |
| --- | --- | --- | --- | --- | --- | --- | --- |
| NREM - NREM to MA (short) | 146.2 | 13.8 | 555 | 10.598 | < .0001*** | 122.69 | 168.92 |
| NREM - NREM to MA (long) | 204.0 | 18.5 | 556 | 11.042 | < .0001*** | 173.29 | 239.32 |
| NREM - NREM to Wake | 246.0 | 19.9 | 554 | 12.338 | < .0001*** | 195.52 | 299.23 |
| NREM to MA (short) - NREM to MA (long) | 57.8 | 19.0 | 554 | 3.046 | 0.0130* | 22.56 | 94.71 |
| NREM to MA (short) - NREM to Wake | 99.8 | 20.2 | 556 | 4.943 | < .0001*** | 48.62 | 153.25 |
| NREM to MA (long) - NREM to Wake | 42.0 | 23.0 | 553 | 1.827 | 0.2618 | -13.03 | 97.16 |

###### 2f: NE (AUC) & RR (AUC) correlation

The correlation between RR and NE showed a negative relationship. Due to issues with heteroscedasticity, the conditional and marginal R squared were bootstrapped, which supported the primary findings.

| **Statistic** | **Value** |
| --- | --- |
| **Fixed Effects Estimates** |  |
| Intercept Estimate | -24.62 |
| Intercept SE | 15.95 |
| Intercept df | 5.48 |
| Intercept t-value | -1.54 |
| Intercept p-value | 0.18 |
| NE Estimate | -5.73 |
| NE SE | 1.29 |
| NE df | 5.87 |
| NE t-value | -4.45 |
| NE p-value | 0.005 |
| **Random Effects** |  |
| SubjectID (Intercept) Variance | 1385.00 |
| SubjectID (Intercept) SD | 37.22 |
| SubjectID NE Variance | 10.77 |
| SubjectID NE SD | 3.28 |
| SubjectID NE Corr | -0.61 |
| Residual Variance | 15748.02 |
| Residual SD | 125.49 |
| **R-squared Values** |  |
| Marginal R-squared (R²m) | 0.35 |
| Bootstrapped 2.5% CI | 0.31 |
| Bootstrapped 97.5% CI | 0.50 |
| Conditional R-squared (R²c) | 0.50 |
| Bootstrapped 2.5% CI | 0.47 |
| Bootstrapped 97.5% CI | 0.66 |
| **Correlation of Fixed Effects** |  |
| Correlation (Intercept, NE) | -0.60 |

###### 1h: Delta predicted by Sleep Stage Transitions

The analysis revealed significant effects of Sleep Stage Transition on delta. Specifically, compared to the reference category (NREM), all other levels showed significant decreases in delta. Pairwise comparisons confirmed these differences (all p<.001). Given the presence of heteroscedasticity and slight issues with normality of residuals, bootstrapping was used to provide robust CIs, which supported the primary findings.

**Fixed Effects of Sleep Stage Transitions**

| **Predictor** | **Estimate** | **Std. Error** | **df** | **t-value** | **p-value** |
| --- | --- | --- | --- | --- | --- |
| (Intercept) | 1.088 | 0.574 | 5.584 | 1.895 | 0.111 |
| NREM | 0 | - | - | - | - |
| NREM to MA (short) | -2.619 | 0.469 | 415.597 | -5.581 | < .001*** |
| NREM to MA (long) | -6.696 | 0.642 | 415.402 | -10.436 | < .001*** |
| NREM to Wake | -10.031 | 0.713 | 415.108 | -14.064 | < .001*** |

**Pairwise Comparisons with Tukey Adjustment**

| **Contrast** | **Estimate** | **SE** | **df** | **t-ratio** | **p-value** | **Lower Bound (2.5%)** | **Upper Bound (97.5%)** |
| --- | --- | --- | --- | --- | --- | --- | --- |
| NREM - NREM to MA (short) | 2.62 | 0.472 | 416 | 5.546 | < .0001*** | 1.76 | 3.47 |
| NREM - NREM to MA (long) | 6.70 | 0.646 | 415 | 10.366 | < .0001*** | 5.24 | 8.17 |
| NREM - NREM to Wake | 10.03 | 0.718 | 415 | 13.964 | < .0001*** | 8.40 | 11.51 |
| NREM to MA (short) - NREM to MA (long) | 4.08 | 0.661 | 416 | 6.172 | < .0001*** | 2.61 | 5.60 |
| NREM to MA (short) - NREM to Wake | 7.41 | 0.722 | 415 | 10.261 | < .0001*** | 5.83 | 9.04 |
| NREM to MA (long) - NREM to Wake | 3.33 | 0.829 | 413 | 4.024 | 0.0004*** | 1.36 | 5.28 |

###### 1h: Theta predicted by Sleep Stage Transition

The analysis revealed significant effects of Sleep Stage Transition on theta. Specifically, compared to the reference category (NREM), all other levels showed significant decreases in theta. Pairwise comparisons confirmed these differences (all p<.001). Given issues with normality of residuals and heteroscedasticity, bootstrapping was used to provide robust CIs, which supported the primary findings.

**Fixed Effects of Sleep Stage Transitions**

| **Predictor** | **Estimate** | **Std. Error** | **df** | **t-value** | **p-value** |
| --- | --- | --- | --- | --- | --- |
| (Intercept) | -0.302 | 0.627 | 4.790 | -0.482 | 0.651 |
| NREM | 0 | - | - | - | - |
| NREM to MA (short) | -2.859 | 0.375 | 415.473 | -7.629 | < .001*** |
| NREM to MA (long) | -6.364 | 0.513 | 415.550 | -12.413 | < .001*** |
| NREM to Wake | -8.875 | 0.570 | 415.632 | -15.573 | < .001*** |

**Pairwise Comparisons with Tukey Adjustment**

| **Contrast** | **Estimate** | **SE** | **df** | **t-ratio** | **p-value** | **Lower Bound (2.5%)** | **Upper Bound (97.5%)** |
| --- | --- | --- | --- | --- | --- | --- | --- |
| NREM - NREM to MA (short) | 2.86 | 0.376 | 415 | 7.605 | < .0001*** | 2.229 | 3.487 |
| NREM - NREM to MA (long) | 6.36 | 0.514 | 416 | 12.373 | < .0001*** | 5.270 | 7.464 |
| NREM - NREM to Wake | 8.88 | 0.572 | 416 | 15.520 | < .0001*** | 7.409 | 10.300 |
| NREM to MA (short) - NREM to MA (long) | 3.50 | 0.526 | 414 | 6.660 | < .0001*** | 2.477 | 4.550 |
| NREM to MA (short) - NREM to Wake | 6.02 | 0.575 | 414 | 10.459 | < .0001*** | 4.510 | 7.468 |
| NREM to MA (long) - NREM to Wake | 2.51 | 0.660 | 412 | 3.807 | 0.0009*** | 0.781 | 4.114 |

###### 1h: Sigma predicted by Sleep Stage Transitions

The analysis revealed significant effects of Sleep Stage Transition on sigma. Specifically, compared to the reference category (NREM), all other levels showed significant decreases in sigma. Pairwise comparisons confirmed these differences, except for the comparison between NREM to MA (long) and NREM to Wake, which was only marginally significant. Given issues with non-normality of residuals and heteroscedasticity, bootstrapping was used to provide robust CIs, which supported the primary findings.

**Fixed Effects of Sleep Stage Transitions**

| **Predictor** | **Estimate** | **Std. Error** | **df** | **t-value** | **p-value** |
| --- | --- | --- | --- | --- | --- |
| (Intercept) | -1.999 | 1.294 | 4.224 | -1.544 | 0.194 |
| NREM | 0 | - | - | - | - |
| NREM to MA (short) | -3.091 | 0.439 | 413.367 | -7.042 | < .001*** |
| NREM to MA (long) | -5.940 | 0.601 | 413.411 | -9.893 | < .001*** |
| NREM to Wake | -7.723 | 0.668 | 413.461 | -11.569 | < .001*** |

**Pairwise Comparisons with Tukey Adjustment**

| **Contrast** | **Estimate** | **SE** | **df** | **t-ratio** | **p-value** | **Lower Bound (2.5%)** | **Upper Bound (97.5%)** |
| --- | --- | --- | --- | --- | --- | --- | --- |
| NREM - NREM to MA (short) | 3.09 | 0.439 | 413 | 7.036 | < .0001*** | 2.276 | 3.885 |
| NREM - NREM to MA (long) | 5.94 | 0.601 | 413 | 9.884 | < .0001*** | 4.600 | 7.297 |
| NREM - NREM to Wake | 7.72 | 0.668 | 413 | 11.558 | < .0001*** | 6.129 | 9.409 |
| NREM to MA (short) - NREM to MA (long) | 2.85 | 0.615 | 413 | 4.633 | < .0001*** | 1.491 | 4.129 |
| NREM to MA (short) - NREM to Wake | 4.63 | 0.672 | 413 | 6.891 | < .0001*** | 3.083 | 6.308 |
| NREM to MA (long) - NREM to Wake | 1.78 | 0.771 | 412 | 2.313 | 0.0968 . | -0.058 | 3.719 |

###### 1h: Beta predicted by Sleep Stage Transitions

The analysis revealed significant effects of Sleep Stage Transition on beta. Specifically, compared to the reference category (NREM), all other levels showed significant decreases in beta. Pairwise comparisons confirmed these differences, except for the comparison between NREM to MA (long) and NREM to Wake, which was not significant. Given potential issues with normality of residuals, bootstrapping was used to provide robust CIs, which supported the primary findings.

**Fixed Effects of Sleep Stage Transitions**

| **Predictor** | **Estimate** | **Std. Error** | **df** | **t-value** | **p-value** |
| --- | --- | --- | --- | --- | --- |
| (Intercept) | -1.744 | 0.918 | 4.182 | -1.900 | 0.127 |
| NREM | 0 | - | - | - | - |
| NREM to MA (short) | -1.345 | 0.277 | 413.102 | -4.859 | < .001*** |
| NREM to MA (long) | -2.530 | 0.379 | 413.137 | -6.681 | < .001*** |
| NREM to Wake | -2.777 | 0.421 | 413.178 | -6.595 | < .001*** |

**Pairwise Comparisons with Tukey Adjustment**

| **Contrast** | **Estimate** | **SE** | **df** | **t-ratio** | **p-value** | **Lower Bound (2.5%)** | **Upper Bound (97.5%)** |
| --- | --- | --- | --- | --- | --- | --- | --- |
| NREM - NREM to MA (short) | 1.345 | 0.277 | 413 | 4.855 | < .0001*** | 0.838 | 1.830 |
| NREM - NREM to MA (long) | 2.530 | 0.379 | 413 | 6.676 | < .0001*** | 1.691 | 3.309 |
| NREM - NREM to Wake | 2.777 | 0.421 | 413 | 6.590 | < .0001*** | 1.891 | 3.754 |
| NREM to MA (short) - NREM to MA (long) | 1.185 | 0.388 | 413 | 3.056 | 0.0127* | 0.355 | 1.936 |
| NREM to MA (short) - NREM to Wake | 1.432 | 0.424 | 413 | 3.378 | 0.0044** | 0.562 | 2.386 |
| NREM to MA (long) - NREM to Wake | 0.247 | 0.486 | 412 | 0.508 | 0.9572 | -0.799 | 1.290 |

###### 1h: Gamma predicted by Sleep Stage Transitions

The analysis revealed significant effects of Sleep Stage Transition on gamma. Specifically, compared to the reference category (NREM), NREM to MA (short) showed a significant increase in gamma, while NREM to MA (long) and NREM to Wake showed even larger increases. Pairwise comparisons confirmed these differences, except for the comparison between NREM and NREM to MA (short), which was not significant. Given potential issues with normality of residuals and heteroscedasticity, bootstrapping was used to provide robust CIs, which supported the primary findings.

**Fixed Effects of Sleep Stage Transitions**

| **Predictor** | **Estimate** | **Std. Error** | **df** | **t-value** | **p-value** |
| --- | --- | --- | --- | --- | --- |
| (Intercept) | -0.115 | 0.341 | 5.045 | -0.337 | 0.749 |
| NREM | 0 | - | - | - | - |
| NREM to MA (short) | 0.531 | 0.227 | 415.861 | 2.341 | 0.020* |
| NREM to MA (long) | 1.944 | 0.310 | 415.911 | 6.263 | < .001*** |
| NREM to Wake | 6.489 | 0.345 | 415.955 | 18.804 | < .001*** |

**Pairwise Comparisons with Tukey Adjustment**

| **Contrast** | **Estimate** | **SE** | **df** | **t-ratio** | **p-value** | **Lower Bound (2.5%)** | **Upper Bound (97.5%)** |
| --- | --- | --- | --- | --- | --- | --- | --- |
| NREM - NREM to MA (short) | -0.531 | 0.228 | 416 | -2.331 | 0.0926 . | -0.847 | -0.195 |
| NREM - NREM to MA (long) | -1.944 | 0.312 | 416 | -6.237 | < .0001*** | -2.534 | -1.402 |
| NREM - NREM to Wake | -6.489 | 0.347 | 416 | -18.722 | < .0001*** | -7.867 | -5.117 |
| NREM to MA (short) - NREM to MA (long) | -1.413 | 0.319 | 415 | -4.431 | 0.0001*** | -1.975 | -0.874 |
| NREM to MA (short) - NREM to Wake | -5.958 | 0.349 | 414 | -17.091 | < .0001*** | -7.262 | -4.523 |
| NREM to MA (long) - NREM to Wake | -4.545 | 0.400 | 412 | -11.367 | < .0001*** | -5.970 | -3.146 |

###### 1i: Delta (AUC) & RR (AUC) correlation

The correlation between RR and delta showed a positive relationship. Due to issues with heteroscedasticity, the conditional and marginal R squared were bootstrapped, which supported the primary findings.

| **Statistic** | **Value** |
| --- | --- |
| **Fixed Effects Estimates** |  |
| Intercept Estimate | -58.80 |
| Intercept SE | 17.47 |
| Intercept df | 3.88 |
| Intercept t-value | -3.37 |
| Intercept p-value | 0.03* |
| Delta Estimate | 15.48 |
| Delta SE | 1.28 |
| Delta df | 411.53 |
| Delta t-value | 12.11 |
| Delta p-value | <0.0001*** |
| **Random Effects** |  |
| SubjectID (Intercept) Variance | 1266.00 |
| SubjectID (Intercept) SD | 35.59 |
| Residual Variance | 17480.00 |
| Residual SD | 132.21 |
| **R-squared Values** |  |
| Marginal R-squared (R²m) | 0.27 |
| Bootstrapped 2.5% CI | 0.18 |
| Bootstrapped 97.5% CI | 0.36 |
| Conditional R-squared (R²c) | 0.32 |
| Bootstrapped 2.5% CI | 0.25 |
| Bootstrapped 97.5% CI | 0.41 |
| **Correlation of Fixed Effects** |  |
| Correlation (Intercept, Delta) | 0.14 |

###### 1i: Theta (AUC) & RR (AUC) correlation

The correlation between RR and theta showed a positive relationship. Bootstrapping was not performed, as all assumptions of the model were met.

| **Statistic** | **Value** |
| --- | --- |
| **Fixed Effects Estimates** |  |
| Intercept Estimate | -22.84 |
| Intercept SE | 10.65 |
| Intercept df | 2.82 |
| Intercept t-value | -2.14 |
| Intercept p-value | 0.13 |
| Theta Estimate | 22.43 |
| Theta SE | 3.14 |
| Theta df | 3.82 |
| Theta t-value | 7.15 |
| Theta p-value | 0.002** |
| **Random Effects** |  |
| SubjectID (Intercept) Variance | 285.70 |
| SubjectID (Intercept) SD | 16.90 |
| SubjectID Theta Variance | 39.95 |
| SubjectID Theta SD | 6.32 |
| SubjectID Theta Corr | 0.21 |
| Residual Variance | 14546.88 |
| Residual SD | 120.61 |
| **R-squared Values** |  |
| Marginal R-squared (R²m) | 0.41 |
| Conditional R-squared (R²c) | 0.46 |
| **Correlation of Fixed Effects** |  |
| Correlation (Intercept, Theta) | 0.28 |

###### 1i: Sigma (AUC) & RR (AUC) correlation

The correlation between RR and sigma showed a positive relationship. Due to issues with heteroscedasticity and normality of residuals, the conditional and marginal R squared were bootstrapped, which supported the primary findings.

| **Statistic** | **Value** |
| --- | --- |
| **Fixed Effects Estimates** |  |
| Intercept Estimate | -3.20 |
| Intercept SE | 7.78 |
| Intercept df | 15.37 |
| Intercept t-value | -0.41 |
| Intercept p-value | 0.69 |
| Sigma Estimate | 20.21 |
| Sigma SE | 3.17 |
| Sigma df | 3.83 |
| Sigma t-value | 6.37 |
| Sigma p-value | 0.004** |
| **Random Effects** |  |
| SubjectID (Intercept) Variance | 39.85 |
| SubjectID (Intercept) SD | 6.31 |
| SubjectID Sigma Variance | 42.99 |
| SubjectID Sigma SD | 6.56 |
| SubjectID Sigma Corr | -1.00 |
| Residual Variance | 14985.32 |
| Residual SD | 122.42 |
| **R-squared Values** |  |
| Marginal R-squared (R²m) | 0.42 |
| Bootstrapped 2.5% CI | 0.33 |
| Bootstrapped 97.5% CI | 0.48 |
| Conditional R-squared (R²c) | 0.49 |
| Bootstrapped 2.5% CI | 0.43 |
| Bootstrapped 97.5% CI | 0.59 |
| **Correlation of Fixed Effects** |  |
| Correlation (Intercept, Sigma) | -0.15 |

###### 1i: Beta (AUC) & RR (AUC) correlation

The correlation between RR and beta showed a positive relationship. Due to issues with heteroscedasticity and normality of residuals, the conditional and marginal R squared were bootstrapped, which supported the primary findings.

| **Statistic** | **Value** |
| --- | --- |
| **Fixed Effects Estimates** |  |
| Intercept Estimate | -14.16 |
| Intercept SE | 8.25 |
| Intercept df | 23.20 |
| Intercept t-value | -1.72 |
| Intercept p-value | 0.10 |
| Beta Estimate | 28.86 |
| Beta SE | 4.96 |
| Beta df | 3.84 |
| Beta t-value | 5.82 |
| Beta p-value | 0.005** |
| **Random Effects** |  |
| SubjectID (Intercept) Variance | 28.67 |
| SubjectID (Intercept) SD | 5.35 |
| SubjectID Beta Variance | 97.97 |
| SubjectID Beta SD | 9.90 |
| SubjectID Beta Corr | -1.00 |
| Residual Variance | 17913.55 |
| Residual SD | 133.84 |
| **R-squared Values** |  |
| Marginal R-squared (R²m) | 0.31 |
| Bootstrapped 2.5% CI | 0.20 |
| Bootstrapped 97.5% CI | 0.35 |
| Conditional R-squared (R²c) | 0.38 |
| Bootstrapped 2.5% CI | 0.31 |
| Bootstrapped 97.5% CI | 0.47 |
| **Correlation of Fixed Effects** |  |
| Correlation (Intercept, Beta) | -0.05 |

###### 1i: Gamma (AUC) & RR (AUC) correlation

The correlation between RR and gamma showed a negative relationship. Due to issues with heteroscedasticity and normality of residuals, the conditional and marginal R squared were bootstrapped, which supported the primary findings.

| **Statistic** | **Value** |
| --- | --- |
| **Fixed Effects Estimates** |  |
| Intercept Estimate | -67.04 |
| Intercept SE | 20.28 |
| Intercept df | 4.01 |
| Intercept t-value | -3.31 |
| Intercept p-value | 0.03* |
| Gamma_high Estimate | -22.38 |
| Gamma_high SE | 5.79 |
| Gamma_high df | 3.50 |
| Gamma_high t-value | -3.87 |
| Gamma_high p-value | 0.02* |
| **Random Effects** |  |
| SubjectID (Intercept) Variance | 1741.00 |
| SubjectID (Intercept) SD | 41.73 |
| SubjectID Gamma_high Variance | 116.00 |
| SubjectID Gamma_high SD | 10.77 |
| SubjectID Gamma_high Corr | 0.23 |
| Residual Variance | 20086.00 |
| Residual SD | 141.73 |
| **R-squared Values** |  |
| Marginal R-squared (R²m) | 0.15 |
| Bootstrapped 2.5% CI | 0.06 |
| Bootstrapped 97.5% CI | 0.27 |
| Conditional R-squared (R²c) | 0.26 |
| Bootstrapped 2.5% CI | 0.19 |
| Bootstrapped 97.5% CI | 0.44 |
| **Correlation of Fixed Effects** |  |
| Correlation (Intercept, Gamma_high) | 0.13 |

##### Figure 2: Locus coeruleus stimulation

###### 2e: NE (AUC) pre-stimulation

The analysis examined the effects of laser threshold levels on NE (AUC) pre, with condition (ChR2 and YFP) as an additional predictor. The results revealed significant effects of laser threshold levels on NE (AUC) pre. Specifically, in the ChR2 condition, compared to the reference category (threshold -15), all other levels showed significant increases in NE (AUC) pre. Pairwise comparisons within ChR2 confirmed these differences, with all comparisons being significant except for the comparison between thresholds -15 vs. -10 and -5 vs 0.

In the YFP condition, the differences between laser threshold levels were also significant. Pairwise comparisons within YFP indicated that changes in laser threshold levels led to significant differences in NE (AUC) pre, contradicting the expectation that YFP should not show significant differences across laser levels.

Given the presence of heteroscedasticity, bootstrapping was used to provide robust CIs of the pairwise estimates, which supported the primary findings. The bootstrapped CIs confirmed the significant differences across laser threshold levels within each condition, reinforcing the reliability of the observed effects.

**Fixed Effects of Laser Thresholds and Condition on NE (AUC) pre**

| **Predictor** | **Estimate** | **Std. Error** | **df** | **t-value** | **p-value** |
| --- | --- | --- | --- | --- | --- |
| (Intercept) | -14.553 | 1.769 | 135.096 | -8.225 | < .001 *** |
| Threshold -10 | 4.047 | 1.891 | 3726.678 | 2.140 | 0.032 * |
| Threshold -5 | 12.475 | 1.718 | 3714.947 | 7.263 | < .001 *** |
| Threshold 0 | 14.221 | 1.648 | 3719.405 | 8.631 | < .001 *** |
| Threshold 5 | 21.629 | 1.674 | 3707.315 | 12.921 | < .001 *** |
| Condition YFP | -14.024 | 5.018 | 1079.059 | -2.795 | 0.005 ** |
| Threshold -10 * Condition YFP | -3.566 | 5.372 | 3724.944 | -0.664 | 0.507 |
| Threshold -5 * Condition YFP | 2.941 | 4.973 | 3723.249 | 0.591 | 0.554 |
| Threshold 0 * Condition YFP | 13.059 | 4.931 | 3726.437 | 2.648 | 0.008 ** |
| Threshold 5 * Condition YFP | 8.973 | 4.959 | 3726.946 | 1.809 | 0.070 . |

**Pairwise Comparisons with Bootstrapped Confidence Intervals**

| **Contrast** | **Estimate** | **SE** | **df** | **t-ratio** | **p-value** | **Lower Bound (2.5%)** | **Upper Bound (97.5%)** |
| --- | --- | --- | --- | --- | --- | --- | --- |
| **Within ChR2** | |  |  |  |  |  |  |
| -15 vs -10 | -4.047 | 1.893 | 3727 | -2.138 | 0.204 | -8.291 | -0.084 |
| -15 vs -5 | -12.475 | 1.720 | 3714 | -7.254 | <.0001 *** | -16.513 | -9.141 |
| -15 vs 0 | -14.221 | 1.650 | 3719 | -8.621 | <.0001 *** | -18.190 | -10.817 |
| -15 vs 5 | -21.629 | 1.676 | 3706 | -12.902 | <.0001 *** | -25.762 | -18.026 |
| -10 vs -5 | -8.428 | 1.285 | 3711 | -6.558 | <.0001 *** | -10.616 | -6.129 |
| -10 vs 0 | -10.174 | 1.181 | 3727 | -8.613 | <.0001 *** | -12.210 | -7.926 |
| -10 vs 5 | -17.582 | 1.211 | 3723 | -14.514 | <.0001 *** | -19.756 | -15.269 |
| -5 vs 0 | -1.746 | 0.890 | 3417 | -1.961 | 0.285 | -2.845 | -0.653 |
| -5 vs 5 | -9.154 | 0.921 | 3504 | -9.943 | <.0001 *** | -10.405 | -8.019 |
| 0 vs 5 | -7.408 | 0.746 | 3726 | -9.930 | <.0001 *** | -8.360 | -6.486 |
| **Within YFP** | |  |  |  |  |  |  |
| -15 vs -10 | -0.481 | 5.029 | 3724 | -0.096 | 1.000 | -6.734 | 5.768 |
| -15 vs -5 | -15.416 | 4.667 | 3720 | -3.303 | 0.0086 ** | -19.748 | -10.687 |
| -15 vs 0 | -27.280 | 4.649 | 3726 | -5.868 | <.0001 *** | -32.024 | -22.476 |
| -15 vs 5 | -30.603 | 4.670 | 3726 | -6.553 | <.0001 *** | -35.511 | -25.931 |
| -10 vs -5 | -14.935 | 2.247 | 3725 | -6.646 | <.0001 *** | -20.206 | -9.317 |
| -10 vs 0 | -26.799 | 2.190 | 3704 | -12.236 | <.0001 *** | -31.969 | -21.407 |
| -10 vs 5 | -30.121 | 2.236 | 3682 | -13.469 | <.0001 *** | -35.410 | -24.871 |
| -5 vs 0 | -11.864 | 1.152 | 3576 | -10.294 | <.0001 *** | -14.789 | -8.869 |
| -5 vs 5 | -15.186 | 1.232 | 3511 | -12.322 | <.0001 *** | -18.528 | -12.125 |
| 0 vs 5 | -3.322 | 1.034 | 3723 | -3.213 | 0.0116 * | -6.327 | -0.301 |
| **Between Conditions** | | |  |  |  |  |  |
| Threshold -15 | 14.024 | 5.02 | 1043.0 | 2.794 | 0.0053 ** | 8.497 | 19.298 |
| Threshold -10 | 17.590 | 2.65 | 106.4 | 6.627 | <.0001 *** | 12.424 | 22.509 |
| Threshold -5 | 11.083 | 1.71 | 18.9 | 6.478 | <.0001 *** | 8.762 | 13.373 |
| Threshold 0 | 0.965 | 1.53 | 12.2 | 0.630 | 0.5405 | -1.263 | 3.111 |
| Threshold 5 | 5.051 | 1.60 | 14.6 | 3.147 | 0.0068 ** | 2.885 | 7.045 |

###### 2e: NE (AUC) post-stimulation

The analysis examined the effects of laser threshold levels on NE (AUC) post, with condition (ChR2 and YFP) as an additional predictor. The results revealed significant effects of laser threshold levels on NE (AUC) post. Specifically, in the ChR2 condition, compared to the reference category (threshold -15), all other levels showed significant increases in NE (AUC) post. Pairwise comparisons within ChR2 suggested a plateau in effect one reaching threshold -5.

In the YFP condition, the differences between laser threshold levels were also significant. Pairwise comparisons within YFP indicated that changes in laser threshold levels led to significant differences in NE (AUC) post, contradicting the expectation that YFP should not show significant differences across laser levels.

Given the presence of heteroscedasticity, bootstrapping was used to provide robust CIs of the pairwise estimates, which supported most of the primary findings.

**Fixed Effects of Laser Thresholds and Condition on NE (AUC) post**

| **Predictor** | **Estimate** | **Std. Error** | **df** | **t-value** | **p-value** |
| --- | --- | --- | --- | --- | --- |
| (Intercept) | 17.211 | 2.073 | 143.029 | 8.303 | < .001 *** |
| Threshold -10 | -8.058 | 2.224 | 3726.546 | -3.624 | < .001 *** |
| Threshold -5 | -15.377 | 2.020 | 3713.768 | -7.614 | < .001 *** |
| Threshold 0 | -15.564 | 1.937 | 3718.492 | -8.033 | < .001 *** |
| Threshold 5 | -17.392 | 1.968 | 3705.436 | -8.837 | < .001 *** |
| Condition YFP | -47.444 | 5.893 | 1134.363 | -8.051 | < .001 *** |
| Threshold -10 * Condition YFP | 25.921 | 6.317 | 3725.084 | 4.104 | < .001 *** |
| Threshold -5 * Condition YFP | 43.415 | 5.848 | 3723.405 | 7.424 | < .001 *** |
| Threshold 0 * Condition YFP | 44.864 | 5.798 | 3726.536 | 7.738 | < .001 *** |
| Threshold 5 * Condition YFP | 48.845 | 5.831 | 3726.978 | 8.376 | < .001 *** |

**Pairwise Comparisons with Bootstrapped Confidence Intervals**

| **Contrast** | **Estimate** | **SE** | **df** | **t-ratio** | **p-value** | **Lower Bound (2.5%)** | **Upper Bound (97.5%)** |
| --- | --- | --- | --- | --- | --- | --- | --- |
| **Within ChR2** |  |  |  |  |  |  |  |
| -15 vs -10 | 8.058 | 2.225 | 3727 | 3.621 | 0.003 ** | 3.452 | 12.353 |
| -15 vs -5 | 15.377 | 2.022 | 3713 | 7.605 | <.0001 *** | 11.224 | 19.118 |
| -15 vs 0 | 15.564 | 1.940 | 3718 | 8.024 | <.0001 *** | 11.327 | 19.434 |
| -15 vs 5 | 17.392 | 1.971 | 3704 | 8.823 | <.0001 *** | 12.961 | 21.384 |
| -10 vs -5 | 7.319 | 1.511 | 3709 | 4.843 | <.0001 *** | 5.264 | 9.233 |
| -10 vs 0 | 7.505 | 1.389 | 3727 | 5.403 | <.0001 *** | 5.433 | 9.527 |
| -10 vs 5 | 9.334 | 1.424 | 3722 | 6.553 | <.0001 *** | 7.080 | 11.520 |
| -5 vs 0 | 0.186 | 1.047 | 3398 | 0.178 | 0.9998 | -0.891 | 1.212 |
| -5 vs 5 | 2.015 | 1.082 | 3489 | 1.861 | 0.339 | 0.586 | 3.354 |
| 0 vs 5 | 1.828 | 0.877 | 3726 | 2.084 | 0.227 | 0.722 | 3.027 |
| **Within YFP** | |  |  |  |  |  |  |
| -15 vs -10 | -17.863 | 5.914 | 3724 | -3.021 | 0.021 * | -42.832 | 5.741 |
| -15 vs -5 | -28.037 | 5.488 | 3720 | -5.109 | <.0001 *** | -51.790 | -6.886 |
| -15 vs 0 | -29.301 | 5.466 | 3726 | -5.360 | <.0001 *** | -52.760 | -8.109 |
| -15 vs 5 | -31.453 | 5.491 | 3726 | -5.728 | <.0001 *** | -55.812 | -9.625 |
| -10 vs -5 | -10.174 | 2.642 | 3725 | -3.850 | 0.001 ** | -18.442 | -2.793 |
| -10 vs 0 | -11.438 | 2.575 | 3702 | -4.441 | 0.0001 *** | -19.301 | -4.239 |
| -10 vs 5 | -13.590 | 2.629 | 3678 | -5.168 | <.0001 *** | -21.629 | -6.058 |
| -5 vs 0 | -1.263 | 1.355 | 3564 | -0.932 | 0.884 | -5.401 | 2.372 |
| -5 vs 5 | -3.415 | 1.449 | 3496 | -2.357 | 0.128 | -7.016 | 0.238 |
| 0 vs 5 | -2.152 | 1.216 | 3723 | -1.770 | 0.391 | -5.296 | 1.131 |
| **Between Conditions** | | | |  |  |  |  |
| Threshold -15 | 47.44 | 5.90 | 1085.1 | 8.048 | <.0001 *** | 25.520 | 70.724 |
| Threshold -10 | 21.52 | 3.11 | 110.9 | 6.924 | <.0001 *** | 14.107 | 29.289 |
| Threshold -5 | 4.03 | 1.99 | 19.2 | 2.023 | 0.057 . | 1.109 | 7.200 |
| Threshold 0 | 2.58 | 1.78 | 12.3 | 1.449 | 0.172 | 0.136 | 5.080 |
| Threshold 5 | -1.40 | 1.87 | 14.8 | -0.750 | 0.465 | -3.557 | 0.836 |

###### 2f: RR (AUC) pre-stimulation

The analysis examined the effects of laser threshold levels on RR (AUC) pre, with condition (ChR2 and YFP) as an additional predictor. The results revealed significant effects of laser threshold levels on RR (AUC) pre. Specifically, in the ChR2 condition, compared to the reference category (threshold -15), thresholds above -10 showed significant increases in RR (AUC) pre. Pairwise comparisons within ChR2 found most comparisons to the -10 threshold significant but revealed a plateau in the higher thresholds.

In the YFP condition, the differences between laser threshold levels were generally not significant. This supports the expectation that YFP should not show significant differences across most laser levels.

Given the presence of heteroscedasticity, bootstrapping was used to provide robust CIs, which supported the primary findings. The bootstrapped CIs supported the significant differences across laser threshold levels within each condition.

**Fixed Effects of Laser Thresholds and Condition on RR (AUC) pre**

| **Predictor** | **Estimate** | **Std. Error** | **df** | **t-value** | **p-value** |
| --- | --- | --- | --- | --- | --- |
| (Intercept) | 50.856 | 12.375 | 310.988 | 4.110 | < .001 *** |
| Threshold -10 | -0.466 | 14.110 | 3705.490 | -0.033 | 0.974 |
| Threshold -5 | -27.935 | 12.801 | 3623.267 | -2.182 | 0.029 * |
| Threshold 0 | -32.274 | 12.283 | 3637.806 | -2.628 | 0.009 ** |
| Threshold 5 | -44.725 | 12.469 | 3549.423 | -3.587 | < .001 *** |
| Condition YFP | -37.196 | 36.762 | 2079.484 | -1.012 | 0.312 |
| Threshold -10 * Condition YFP | 10.633 | 40.111 | 3726.932 | 0.265 | 0.791 |
| Threshold -5 * Condition YFP | 63.396 | 37.141 | 3725.643 | 1.707 | 0.088 . |
| Threshold 0 * Condition YFP | 45.910 | 36.806 | 3725.062 | 1.247 | 0.212 |
| Threshold 5 * Condition YFP | 39.670 | 37.011 | 3720.587 | 1.072 | 0.284 |

**Pairwise Comparisons with Bootstrapped Confidence Intervals**

| **Contrast** | **Estimate** | **SE** | **df** | **t-ratio** | **p-value** | **Lower Bound (2.5%)** | **Upper Bound (97.5%)** |
| --- | --- | --- | --- | --- | --- | --- | --- |
| **Within ChR2** |  |  |  |  |  |  |  |
| -15 vs -10 | 0.466 | 14.13 | 3707 | 0.033 | 1.000 | -28.895 | 33.002 |
| -15 vs -5 | 27.935 | 12.83 | 3629 | 2.177 | 0.189 | 0.935 | 54.565 |
| -15 vs 0 | 32.274 | 12.31 | 3643 | 2.621 | 0.067 | 3.768 | 58.139 |
| -15 vs 5 | 44.725 | 12.51 | 3559 | 3.575 | 0.003 ** | 17.085 | 71.445 |
| -10 vs -5 | 27.469 | 9.59 | 3631 | 2.865 | 0.034 * | 9.434 | 44.701 |
| -10 vs 0 | 31.808 | 8.82 | 3716 | 3.606 | 0.003 ** | 15.261 | 48.573 |
| -10 vs 5 | 44.259 | 9.04 | 3670 | 4.894 | <.0001 *** | 26.179 | 61.365 |
| -5 vs 0 | 4.339 | 6.62 | 2748 | 0.655 | 0.966 | -7.635 | 17.316 |
| -5 vs 5 | 16.790 | 6.85 | 2910 | 2.450 | 0.103 | 3.285 | 30.222 |
| 0 vs 5 | 12.451 | 5.57 | 3709 | 2.236 | 0.167 | 1.648 | 24.152 |
| **Within YFP** |  |  |  |  |  |  |  |
| -15 vs -10 | -10.167 | 37.56 | 3726 | -0.271 | 0.999 | -63.918 | 41.101 |
| -15 vs -5 | -35.461 | 34.87 | 3720 | -1.017 | 0.848 | -85.618 | 11.548 |
| -15 vs 0 | -13.636 | 34.72 | 3727 | -0.393 | 0.995 | -65.771 | 34.121 |
| -15 vs 5 | 5.055 | 34.87 | 3726 | 0.145 | 1.000 | -45.285 | 54.365 |
| -10 vs -5 | -25.294 | 16.77 | 3709 | -1.508 | 0.557 | -49.721 | 2.577 |
| -10 vs 0 | -3.469 | 16.33 | 3603 | -0.212 | 0.999 | -28.797 | 24.829 |
| -10 vs 5 | 15.222 | 16.67 | 3518 | 0.913 | 0.892 | -11.151 | 43.884 |
| -5 vs 0 | 21.824 | 8.59 | 3083 | 2.542 | 0.082 | 7.446 | 37.159 |
| -5 vs 5 | 40.515 | 9.18 | 2884 | 4.415 | 0.0001 *** | 24.332 | 55.824 |
| 0 vs 5 | 18.691 | 7.72 | 3725 | 2.421 | 0.110 | 4.310 | 33.311 |
| **Between Conditions** | | |  |  |  |  |  |
| Threshold -15 | 37.20 | 36.79 | 2132.3 | 1.011 | 0.312 | -16.527 | 90.273 |
| Threshold -10 | 26.56 | 18.46 | 281.2 | 1.439 | 0.151 | -4.934 | 53.694 |
| Threshold -5 | -26.20 | 10.55 | 32.7 | -2.484 | 0.018 * | -40.795 | -11.227 |
| Threshold 0 | -8.71 | 8.89 | 16.7 | -0.981 | 0.341 | -20.370 | 3.931 |
| Threshold 5 | -2.47 | 9.57 | 22.2 | -0.259 | 0.798 | -17.233 | 11.725 |

###### 2f: RR (AUC) post-stimulation

The analysis examined the effects of laser threshold levels on RR (AUC) post, with condition (ChR2 and YFP) as an additional predictor. The results revealed significant effects of laser threshold levels on RR (AUC) post. Specifically, in the ChR2 condition, compared to the reference category (threshold -15), all other levels showed significant changes in RR (AUC) post. Pairwise comparisons within ChR2 suggested that the effect of increasing thresholds on RR (AUC) post plateaued around -5.

In the YFP condition, the differences between laser threshold levels were generally not significant, with none of the pairwise comparisons showing significant differences. This supports the expectation that YFP should not show significant differences across most laser levels.

Given the presence of a slight issue with normality, bootstrapping was used to provide robust CIs, which supported the primary findings. The bootstrapped CIs confirmed the significant differences across laser threshold levels within each condition, reinforcing the reliability of the observed effects.

**Fixed Effects of Laser Thresholds and Condition on RR (AUC) post**

| **Predictor** | **Estimate** | **Std. Error** | **df** | **t-value** | **p-value** |
| --- | --- | --- | --- | --- | --- |
| (Intercept) | -110.50 | 12.25 | 366.64 | -9.022 | < .001 *** |
| Threshold -10 | 55.42 | 14.11 | 3689.11 | 3.928 | < .001 *** |
| Threshold -5 | 95.08 | 12.80 | 3571.95 | 7.431 | < .001 *** |
| Threshold 0 | 101.83 | 12.28 | 3586.07 | 8.293 | < .001 *** |
| Threshold 5 | 105.82 | 12.46 | 3454.70 | 8.492 | < .001 *** |
| Condition YFP | 129.76 | 36.67 | 2292.30 | 3.539 | 0.0004 *** |
| Threshold -10 * Condition YFP | -69.63 | 40.12 | 3726.98 | -1.735 | 0.083 . |
| Threshold -5 * Condition YFP | -90.30 | 37.15 | 3726.22 | -2.431 | 0.015 * |
| Threshold 0 * Condition YFP | -106.23 | 36.81 | 3722.38 | -2.886 | 0.004 ** |
| Threshold 5 * Condition YFP | -112.49 | 37.01 | 3714.68 | -3.039 | 0.002 ** |

**Pairwise Comparisons with Bootstrapped Confidence Intervals**

| **Contrast** | **Estimate** | **SE** | **df** | **t-ratio** | **p-value** | **Lower Bound (2.5%)** | **Upper Bound (97.5%)** |
| --- | --- | --- | --- | --- | --- | --- | --- |
| **Within ChR2** |  |  |  |  |  |  |  |
| -15 vs -10 | -55.42 | 14.13 | 3692 | -3.921 | 0.001 *** | -87.889 | -23.605 |
| -15 vs -5 | -95.08 | 12.83 | 3583 | -7.409 | <.0001 *** | -122.084 | -66.815 |
| -15 vs 0 | -101.83 | 12.32 | 3597 | -8.268 | <.0001 *** | -129.420 | -74.061 |
| -15 vs 5 | -105.82 | 12.51 | 3474 | -8.457 | <.0001 *** | -132.810 | -78.654 |
| -10 vs -5 | -39.66 | 9.59 | 3596 | -4.137 | 0.0003 *** | -59.597 | -20.783 |
| -10 vs 0 | -46.41 | 8.82 | 3708 | -5.260 | <.0001 *** | -65.647 | -28.291 |
| -10 vs 5 | -50.40 | 9.04 | 3638 | -5.573 | <.0001 *** | -70.213 | -31.709 |
| -5 vs 0 | -6.75 | 6.61 | 2537 | -1.021 | 0.846 | -17.749 | 4.747 |
| -5 vs 5 | -10.74 | 6.85 | 2697 | -1.569 | 0.518 | -23.600 | 0.704 |
| 0 vs 5 | -3.99 | 5.57 | 3698 | -0.717 | 0.953 | -14.901 | 5.867 |
| **Within YFP** |  |  |  |  |  |  |  |
| -15 vs -10 | 14.20 | 37.57 | 3727 | 0.378 | 0.996 | -81.221 | 104.538 |
| -15 vs -5 | -4.78 | 34.88 | 3720 | -0.137 | 0.999 | -88.016 | 77.310 |
| -15 vs 0 | 4.40 | 34.73 | 3726 | 0.127 | 0.999 | -78.023 | 90.164 |
| -15 vs 5 | 6.67 | 34.88 | 3724 | 0.191 | 0.999 | -78.879 | 91.919 |
| -10 vs -5 | -18.99 | 16.77 | 3701 | -1.132 | 0.790 | -58.290 | 23.878 |
| -10 vs 0 | -9.81 | 16.33 | 3560 | -0.601 | 0.975 | -46.525 | 32.656 |
| -10 vs 5 | -7.54 | 16.67 | 3449 | -0.452 | 0.991 | -46.385 | 34.311 |
| -5 vs 0 | 9.18 | 8.58 | 2887 | 1.070 | 0.822 | -6.242 | 27.325 |
| -5 vs 5 | 11.45 | 9.17 | 2652 | 1.249 | 0.723 | -5.455 | 30.667 |
| 0 vs 5 | 2.27 | 7.72 | 3726 | 0.294 | 0.998 | -13.260 | 17.380 |
| **Between Conditions** | |  |  |  |  |  |  |
| Threshold -15 | -129.8 | 36.70 | 2362.5 | -3.536 | 0.0004 *** | -216.832 | -41.078 |
| Threshold -10 | -60.1 | 18.26 | 347.6 | -3.294 | 0.0011 ** | -103.404 | -13.131 |
| Threshold -5 | -39.5 | 10.19 | 37.8 | -3.873 | 0.0004 *** | -56.358 | -20.774 |
| Threshold 0 | -23.5 | 8.46 | 18.2 | -2.782 | 0.012 * | -35.098 | -22.590 |
| Threshold 5 | -17.3 | 9.17 | 24.8 | -1.883 | 0.072 . | -31.548 | -1.930 |

###### 2g: NE (AUC) pre-stimulation minus post-stimulation

The analysis examined the effects of laser threshold levels on NE (AUC) post-pre, with condition (ChR2 and YFP) as an additional predictor. The results revealed significant effects of laser threshold levels on NE (AUC) post-pre. Specifically, in the ChR2 condition, compared to the reference category (threshold -15), all other levels showed significant increases in NE (AUC) post-pre. Pairwise comparisons within ChR2 confirmed these differences, with all comparisons being significant except for the comparison between thresholds -5 and 0.

In the YFP condition, the differences between laser threshold levels were not as present as in the case of ChR2. This supports the expectation that YFP should not show significant differences across most laser levels.

Given the presence of slight heteroscedasticity, bootstrapping was used to provide robust CIs, which supported the primary findings. The bootstrapped CIs confirmed the significant differences across laser threshold levels within each condition, reinforcing the reliability of the observed effects.

**Fixed Effects of Laser Thresholds and Condition on NE (AUC) post-pre**

| **Predictor** | **Estimate** | **Std. Error** | **df** | **t-value** | **p-value** |
| --- | --- | --- | --- | --- | --- |
| (Intercept) | -31.716 | 2.766 | 100.732 | -11.465 | < .001 *** |
| Threshold -10 | 12.044 | 2.875 | 3726.985 | 4.190 | < .001 *** |
| Threshold -5 | 27.799 | 2.611 | 3721.927 | 10.646 | < .001 *** |
| Threshold 0 | 29.760 | 2.505 | 3724.377 | 11.880 | < .001 *** |
| Threshold 5 | 38.994 | 2.545 | 3718.380 | 15.322 | < .001 *** |
| Condition YFP | 33.333 | 7.693 | 805.349 | 4.333 | < .001 *** |
| Threshold -10 * Condition YFP | -29.407 | 8.164 | 3724.177 | -3.602 | < .001 *** |
| Threshold -5 * Condition YFP | -40.407 | 7.558 | 3722.600 | -5.347 | < .001 *** |
| Threshold 0 * Condition YFP | -31.723 | 7.494 | 3725.681 | -4.233 | < .001 *** |
| Threshold 5 * Condition YFP | -39.779 | 7.537 | 3726.459 | -5.278 | < .001 *** |

**Pairwise Comparisons with Bootstrapped Confidence Intervals**

| **Contrast** | **Estimate** | **SE** | **df** | **t-ratio** | **p-value** | **Lower Bound (2.5%)** | **Upper Bound (97.5%)** |
| --- | --- | --- | --- | --- | --- | --- | --- |
| **Within ChR2** |  |  |  |  |  |  |  |
| -15 vs -10 | -12.044 | 2.88 | 3727 | -4.188 | 0.0003 *** | -18.290 | -6.235 |
| -15 vs -5 | -27.799 | 2.61 | 3722 | -10.636 | <.0001 *** | -33.339 | -22.758 |
| -15 vs 0 | -29.760 | 2.51 | 3724 | -11.870 | <.0001 *** | -35.176 | -24.875 |
| -15 vs 5 | -38.994 | 2.55 | 3718 | -15.304 | <.0001 *** | -44.601 | -33.699 |
| -10 vs -5 | -15.756 | 1.95 | 3719 | -8.065 | <.0001 *** | -18.948 | -12.521 |
| -10 vs 0 | -17.716 | 1.80 | 3727 | -9.868 | <.0001 *** | -20.801 | -14.488 |
| -10 vs 5 | -26.950 | 1.84 | 3726 | -14.639 | <.0001 *** | -30.146 | -23.367 |
| -5 vs 0 | -1.960 | 1.35 | 3527 | -1.448 | 0.597 | -3.533 | -0.392 |
| -5 vs 5 | -11.194 | 1.40 | 3590 | -7.997 | <.0001 *** | -13.014 | -9.199 |
| 0 vs 5 | -9.234 | 1.13 | 3727 | -8.145 | <.0001 *** | -10.771 | -7.677 |
| **Within YFP** |  |  |  |  |  |  |  |
| -15 vs -10 | 17.363 | 7.64 | 3723 | 2.272 | 0.154 | -8.328 | 44.523 |
| -15 vs -5 | 12.607 | 7.09 | 3719 | 1.778 | 0.387 | -10.574 | 37.275 |
| -15 vs 0 | 1.963 | 7.06 | 3725 | 0.278 | 0.999 | -21.654 | 26.853 |
| -15 vs 5 | 0.786 | 7.10 | 3725 | 0.111 | 1.000 | -22.424 | 25.824 |
| -10 vs -5 | -4.756 | 3.42 | 3727 | -1.392 | 0.633 | -14.042 | 4.818 |
| -10 vs 0 | -15.400 | 3.33 | 3715 | -4.626 | <.0001 *** | -24.814 | -5.904 |
| -10 vs 5 | -16.578 | 3.40 | 3702 | -4.876 | <.0001 *** | -26.094 | -6.523 |
| -5 vs 0 | -10.644 | 1.75 | 3638 | -6.075 | <.0001 *** | -15.251 | -6.351 |
| -5 vs 5 | -11.822 | 1.87 | 3597 | -6.309 | <.0001 *** | -16.059 | -7.766 |
| 0 vs 5 | -1.177 | 1.57 | 3722 | -0.749 | 0.945 | -5.226 | 1.131 |
| **Between Conditions** | | | |  |  |  |  |
| Threshold -15 | -33.33 | 7.70 | 775.4 | -4.332 | <.0001 *** | -58.199 | -9.676 |
| Threshold -10 | -3.93 | 4.16 | 79.6 | -0.944 | 0.348 | -13.131 | 5.543 |
| Threshold -5 | 7.07 | 2.79 | 16.4 | 2.532 | 0.022 * | 3.547 | 10.590 |
| Threshold 0 | -1.61 | 2.54 | 11.3 | -0.633 | 0.539 | -4.912 | 1.463 |
| Threshold 5 | 6.45 | 2.64 | 13.2 | 2.438 | 0.030 * | 3.531 | 9.321 |

###### 2g: RR (AUC) pre-stimulation minus post-stimulation

The analysis examined the effects of laser threshold levels on RR (AUC) post-pre, with condition (ChR2 and YFP) as an additional predictor. The results revealed significant effects of laser threshold levels on RR (AUC) post-pre. Specifically, in the ChR2 condition, compared to the reference category (threshold -15), all other levels showed significant changes in RR (AUC) post-pre. Pairwise comparisons within ChR2 suggested a plateau of this effect at -5.

In the YFP condition, the differences between laser threshold levels were not significant, with none of the pairwise comparisons showing significant differences. This supports the expectation that YFP should not show significant differences across most laser levels.

Given the presence of slight heteroscedasticity, bootstrapping was used to provide robust CIs, which supported the primary findings. The bootstrapped CIs confirmed the significant differences across laser threshold levels within each condition, reinforcing the reliability of the observed effects.

**Fixed Effects of Laser Thresholds and Condition on RR (AUC) post-pre**

| **Predictor** | **Estimate** | **Std. Error** | **df** | **t-value** | **p-value** |
| --- | --- | --- | --- | --- | --- |
| (Intercept) | 160.73 | 16.76 | 681.89 | 9.590 | < .001 *** |
| Threshold -10 | -56.08 | 19.90 | 3527.58 | -2.818 | 0.005 ** |
| Threshold -5 | -122.98 | 18.01 | 3225.86 | -6.827 | < .001 *** |
| Threshold 0 | -133.31 | 17.29 | 3162.90 | -7.712 | < .001 *** |
| Threshold 5 | -149.05 | 17.52 | 2760.78 | -8.509 | < .001 *** |
| Condition YFP | -167.28 | 51.41 | 3018.69 | -3.254 | 0.001 ** |
| Threshold -10 * Condition YFP | 80.47 | 56.69 | 3722.83 | 1.420 | 0.156 |
| Threshold -5 * Condition YFP | 153.75 | 52.51 | 3726.41 | 2.928 | 0.003 ** |
| Threshold 0 * Condition YFP | 151.93 | 51.98 | 3692.33 | 2.923 | 0.003 ** |
| Threshold 5 * Condition YFP | 151.22 | 52.25 | 3658.67 | 2.894 | 0.004 ** |

**Pairwise Comparisons**

| **Contrast** | **Estimate** | **SE** | **df** | **t-ratio** | **p-value** | **Lower Bound (2.5%)** | **Upper Bound (97.5%)** |
| --- | --- | --- | --- | --- | --- | --- | --- |
| **Within ChR2** |  |  |  |  |  |  |  |
| -15 vs -10 | 56.08 | 19.98 | 3513 | 2.807 | 0.040 * | 10.503 | 100.384 |
| -15 vs -5 | 122.98 | 18.12 | 3194 | 6.788 | <.0001 *** | 83.054 | 161.775 |
| -15 vs 0 | 133.31 | 17.40 | 3127 | 7.661 | <.0001 *** | 95.144 | 172.267 |
| -15 vs 5 | 149.05 | 17.68 | 2708 | 8.432 | <.0001 *** | 109.194 | 189.365 |
| -10 vs -5 | 66.90 | 13.52 | 3372 | 4.949 | <.0001 *** | 40.400 | 93.797 |
| -10 vs 0 | 77.23 | 12.46 | 3623 | 6.196 | <.0001 *** | 51.124 | 104.550 |
| -10 vs 5 | 92.97 | 12.78 | 3315 | 7.277 | <.0001 *** | 66.218 | 120.731 |
| -5 vs 0 | 10.32 | 9.28 | 1681 | 1.113 | 0.800 | -5.204 | 27.874 |
| -5 vs 5 | 26.07 | 9.62 | 1707 | 2.709 | 0.053 | 7.334 | 45.774 |
| 0 vs 5 | 15.74 | 7.87 | 3590 | 2.001 | 0.266 | 0.875 | 32.014 |
| **Within YFP** |  |  |  |  |  |  |  |
| -15 vs -10 | -24.39 | 53.11 | 3727 | -0.459 | 0.991 | -113.154 | 63.572 |
| -15 vs -5 | -30.76 | 49.32 | 3721 | -0.624 | 0.971 | -113.046 | 48.050 |
| -15 vs 0 | -18.62 | 49.08 | 3714 | -0.379 | 0.996 | -101.136 | 60.081 |
| -15 vs 5 | -2.17 | 49.29 | 3702 | -0.044 | 1.000 | -83.967 | 76.731 |
| -10 vs -5 | -6.37 | 23.68 | 3658 | -0.269 | 0.999 | -57.294 | 41.927 |
| -10 vs 0 | 5.77 | 23.03 | 3290 | 0.251 | 0.999 | -43.797 | 53.115 |
| -10 vs 5 | 22.23 | 23.48 | 3050 | 0.947 | 0.879 | -25.857 | 69.042 |
| -5 vs 0 | 12.14 | 12.07 | 1894 | 1.006 | 0.853 | -9.903 | 37.517 |
| -5 vs 5 | 28.60 | 12.89 | 1599 | 2.219 | 0.173 | 3.429 | 53.411 |
| 0 vs 5 | 16.46 | 10.92 | 3727 | 1.507 | 0.558 | -4.299 | 37.764 |
| **Between Conditions** | | | |  |  |  |  |
| Threshold -15 | 167.3 | 51.5 | 2976.2 | 3.249 | 0.001 ** | 78.221 | 257.561 |
| Threshold -10 | 86.8 | 25.0 | 724.0 | 3.473 | 0.001 *** | 34.007 | 136.714 |
| Threshold -5 | 13.5 | 12.9 | 68.1 | 1.046 | 0.299 | -8.934 | 35.757 |
| Threshold 0 | 15.3 | 10.1 | 27.1 | 1.514 | 0.142 | -1.847 | 31.499 |
| Threshold 5 | 16.1 | 11.3 | 40.2 | 1.421 | 0.163 | -4.359 | 34.002 |

###### 2h: NE (AUC) & RR (AUC): pre-stimulation minus post stimulation correlation (all thresholds)

The correlation between RR and NE showed a negative relationship. Due to slight issues with normality of residuals, the marginal and conditional R squared estimates were bootstrapped, which supported the primary findings.

| **Statistic** | **Value** |
| --- | --- |
| **Fixed Effects Estimates** |  |
| Intercept Estimate | 29.58 |
| Intercept SE | 4.78 |
| Intercept df | 5.03 |
| Intercept t-value | 6.19 |
| Intercept p-value | 0.002** |
| NE Estimate | -2.83 |
| NE SE | 0.16 |
| NE df | 2094.54 |
| NE t-value | -17.25 |
| NE p-value | < 0.0001*** |
| **Random Effects** |  |
| SubjectID (Intercept) Variance | 73.97 |
| SubjectID (Intercept) SD | 8.60 |
| Residual Variance | 24315.32 |
| Residual SD | 155.90 |
| **R-squared Values** |  |
| Marginal R-squared (R²m) | 0.11 |
| Bootstrapped 2.5% CI | 0.08 |
| Bootstrapped 97.5% CI | 0.14 |
| Conditional R-squared (R²c) | 0.11 |
| Bootstrapped 2.5% CI | 0.09 |
| Bootstrapped 97.5% CI | 0.15 |
| **Correlation of Fixed Effects** |  |
| Correlation (Intercept, NE) | 0.08 |

###### 2h: NE (AUC) & RR (AUC): pre-stimulation minus post stimulation correlation (Threshold -15 to -5)

The correlation between RR and NE showed a negative relationship. Due to slight issues with normality of residuals, the marginal and conditional R squared estimates were bootstrapped, which supported the primary findings.

| **Statistic** | **Value** |
| --- | --- |
| **Fixed Effects Estimates** |  |
| Intercept Estimate | 32.00 |
| Intercept SE | 8.30 |
| Intercept df | 7.77 |
| Intercept t-value | 3.86 |
| Intercept p-value | 0.005** |
| NE Estimate | -3.43 |
| NE SE | 0.27 |
| NE df | 654.00 |
| NE t-value | -12.92 |
| NE p-value | < 0.0001*** |
| **Random Effects** |  |
| SubjectID (Intercept) Variance | 162.00 |
| SubjectID (Intercept) SD | 12.73 |
| Residual Variance | 23211.00 |
| Residual SD | 152.35 |
| **R-squared Values** |  |
| Marginal R-squared (R²m) | 0.18 |
| Bootstrapped 2.5% CI | 0.13 |
| Bootstrapped 97.5% CI | 0.25 |
| Conditional R-squared (R²c) | 0.19 |
| Bootstrapped 2.5% CI | 0.14 |
| Bootstrapped 97.5% CI | 0.27 |
| **Correlation of Fixed Effects** |  |
| Correlation (Intercept, NE) | 0.37 |

###### 2i: RR PSD (AUC)

The analysis examined the effects of laser threshold levels on RR PSD AUC, with condition (ChR2 and YFP) as an additional predictor. The results revealed significant effects of laser threshold levels on RR PSD AUC. Specifically, in the ChR2 condition, compared to the reference category (baseline), higher thresholds indicated an increase in AUC. Pairwise comparisons within ChR2 suggested an increase in RR PSD AUC between baseline and threshold 5.

In the YFP condition, pairwise comparisons suggested no significant differences.

Given the presence of issues with normality, heteroscedasticity, and slight autocorrelation of residuals, block bootstrapping based on Subject ID was used to provide robust CIs. These estimates often overlapped with 0, casting doubt on how accurate the initial pairwise estimates were.

**Fixed Effects of Laser Thresholds and Condition on RR PSD AUC**

| **Effect** | **Estimate** | **Std. Error** | **df** | **t-value** | **p-value** |
| --- | --- | --- | --- | --- | --- |
| Intercept (Baseline) | 3.078e-05 | 3.379e-06 | 8.68 | 9.108 | 9.9e-06 *** |
| -15 vs Baseline | -7.386e-07 | 2.515e-06 | 303.3 | -0.294 | 0.7692 |
| -10 vs Baseline | 1.582e-06 | 2.518e-06 | 303.5 | 0.628 | 0.5304 |
| -5 vs Baseline | 2.401e-06 | 2.489e-06 | 303.6 | 0.964 | 0.3356 |
| 0 vs Baseline | 3.549e-06 | 2.547e-06 | 304.3 | 1.393 | 0.1645 |
| 5 vs Baseline | 9.062e-06 | 2.844e-06 | 304.2 | 3.187 | 0.0016 ** |
| YFP vs ChR2 | -1.727e-06 | 5.356e-06 | 8.77 | -0.322 | 0.7547 |
| -15 vs Baseline * YFP | 2.717e-06 | 3.985e-06 | 303.2 | 0.682 | 0.4959 |
| -10 vs Baseline * YFP | -2.316e-06 | 3.749e-06 | 303.3 | -0.618 | 0.5371 |
| -5 vs Baseline * YFP | -2.171e-06 | 3.824e-06 | 303.3 | -0.568 | 0.5706 |
| 0 vs Baseline * YFP | -5.122e-06 | 3.747e-06 | 303.9 | -1.367 | 0.1726 |
| 5 vs Baseline * YFP | -9.746e-06 | 4.488e-06 | 303.8 | -2.172 | 0.0307 * |

**Pairwise Comparisons with Bootstrapped Confidence Intervals**

| **Contrast** | **Estimate** | **SE** | **df** | **t-ratio** | **p-value** | **Lower Bound (2.5%)** | **Upper Bound (97.5%)** |
| --- | --- | --- | --- | --- | --- | --- | --- |
| **Within ChR2** |  |  |  |  |  |  |  |
| Baseline vs -15 | 7.39e-07 | 2.51e-06 | 7.19e+07 | 0.294 | 0.9997 | -6.79e-06 | 1.09e-05 |
| Baseline vs -10 | -1.58e-06 | 2.52e-06 | 1.25e+08 | -0.628 | 0.9890 | -1.07e-05 | 1.11e-05 |
| Baseline vs -5 | -2.40e-06 | 2.49e-06 | 1.43e+08 | -0.964 | 0.9292 | -5.22e-06 | 1.60e-05 |
| Baseline vs 0 | -3.55e-06 | 2.55e-06 | 1.33e+08 | -1.393 | 0.7312 | -9.79e-06 | 1.86e-05 |
| Baseline vs 5 | -9.06e-06 | 2.84e-06 | 7.70e+07 | -3.187 | 0.0180 * | -7.35e-06 | 1.57e-05 |
| -15 vs -10 | -2.32e-06 | 3.27e-06 | 9.25e+07 | -0.709 | 0.9810 | -5.89e-06 | 4.91e-06 |
| -15 vs -5 | -3.14e-06 | 3.26e-06 | 9.87e+07 | -0.964 | 0.9293 | -6.96e-06 | 5.45e-06 |
| -15 vs 0 | -4.29e-06 | 3.29e-06 | 9.68e+07 | -1.303 | 0.7836 | -1.14e-05 | 3.71e-06 |
| -15 vs 5 | -9.80e-06 | 3.52e-06 | 7.31e+07 | -2.785 | 0.0598 | -1.06e-05 | 3.34e-06 |
| -10 vs -5 | -8.19e-07 | 3.27e-06 | 1.40e+08 | -0.251 | 0.9999 | -1.06e-05 | 3.34e-06 |
| -10 vs 0 | -1.97e-06 | 3.27e-06 | 1.38e+08 | -0.602 | 0.9910 | -8.35e-06 | 3.16e-06 |
| -10 vs 5 | -7.48e-06 | 3.51e-06 | 9.56e+07 | -2.134 | 0.2698 | -8.35e-06 | 3.16e-06 |
| -5 vs 0 | -1.15e-06 | 3.30e-06 | 1.45e+08 | -0.348 | 0.9993 | -8.61e-06 | 3.34e-06 |
| -5 vs 5 | -6.66e-06 | 3.52e-06 | 9.85e+07 | -1.890 | 0.4080 | -1.06e-05 | 3.16e-06 |
| 0 vs 5 | -5.51e-06 | 3.49e-06 | 1.02e+08 | -1.578 | 0.6128 | -9.19e-06 | 3.16e-06 |
| **Within YFP** |  |  |  |  |  |  |  |
| Baseline vs -15 | -1.98e-06 | 3.09e-06 | 1.00e+08 | -0.640 | 0.9880 | -9.53e-06 | 4.91e-06 |
| Baseline vs -10 | 7.35e-07 | 2.78e-06 | 9.58e+07 | 0.265 | 0.9998 | -7.15e-06 | 4.91e-06 |
| Baseline vs -5 | -2.30e-07 | 2.90e-06 | 8.80e+07 | -0.079 | 1.0000 | -7.38e-06 | 3.06e-06 |
| Baseline vs 0 | 1.57e-06 | 2.75e-06 | 1.00e+08 | 0.572 | 0.9928 | -8.31e-06 | -6.42e-07 |
| Baseline vs 5 | 6.84e-07 | 3.47e-06 | 1.05e+08 | 0.197 | 1.0000 | -6.56e-06 | 5.83e-06 |
| -15 vs -10 | 2.71e-06 | 3.77e-06 | 9.92e+07 | 0.719 | 0.9797 | -5.64e-06 | 9.95e-06 |
| -15 vs -5 | 1.75e-06 | 3.87e-06 | 9.49e+07 | 0.452 | 0.9976 | -8.61e-06 | 7.81e-06 |
| -15 vs 0 | 3.55e-06 | 3.74e-06 | 1.03e+08 | 0.949 | 0.9337 | -8.94e-06 | 6.01e-06 |
| -15 vs 5 | 2.66e-06 | 4.30e-06 | 1.05e+08 | 0.619 | 0.9897 | -5.66e-06 | 5.02e-06 |
| -10 vs -5 | -9.64e-07 | 3.61e-06 | 9.23e+07 | -0.267 | 0.9998 | -1.13e-05 | 4.43e-06 |
| -10 vs 0 | 8.39e-07 | 3.49e-06 | 9.96e+07 | 0.240 | 0.9999 | -1.16e-05 | 4.43e-06 |
| -10 vs 5 | -5.10e-08 | 4.08e-06 | 1.03e+08 | -0.013 | 1.0000 | -1.20e-05 | 1.34e-05 |
| -5 vs 0 | 1.80e-06 | 3.59e-06 | 9.46e+07 | 0.502 | 0.9961 | -1.06e-05 | 5.02e-06 |
| -5 vs 5 | 9.13e-07 | 4.17e-06 | 9.97e+07 | 0.219 | 0.9999 | -7.11e-06 | 1.27e-05 |
| 0 vs 5 | -8.90e-07 | 4.04e-06 | 1.07e+08 | -0.220 | 0.9999 | -5.66e-06 | 5.02e-06 |
| **ChR2 vs YFP** |  |  |  |  |  |  |  |
| Threshold 0 | 1.73e-06 | 5.40e-06 | 7061 | 0.320 | 0.7489 | -8.61e-06 | 3.34e-06 |
| Threshold -15 | -9.90e-07 | 6.33e-06 | 13355 | -0.156 | 0.8757 | -8.61e-06 | 3.34e-06 |
| Threshold -10 | 4.04e-06 | 6.18e-06 | 12378 | 0.654 | 0.5131 | -1.13e-05 | 4.43e-06 |
| Threshold -5 | 3.90e-06 | 6.23e-06 | 12787 | 12787 | 0.5316 | -1.16e-05 | 4.43e-06 |
| Threshold 0 | 6.85e-06 | 6.17e-06 | 12426 | 1.110 | 0.2672 | -1.13e-05 | 4.43e-06 |
| Threshold 5 | 1.15e-05 | 6.64e-06 | 16411 | 1.727 | 0.0842 | -1.16e-05 | 4.43e-06 |

###### 2i: RR PSD frequency at peak power

The analysis examined the effects of laser threshold levels on RR frequency at peak power of the PSD, with condition (ChR2 and YFP) as an additional predictor. The results revealed significant effects of laser threshold levels on RR frequency at peak power of the PSD at higher thresholds. Specifically, in the ChR2 condition, pairwise comparisons found significant increases in frequency at peak power between lower and higher levels.

In the YFP condition, pairwise comparisons did not suggest any differences.

Given the presence of issues with normality, and slight autocorrelation of residuals, block bootstrapping based on Subject ID was used to provide robust CIs. These estimates tended to not overlap with 0 within the ChR2 comparisons, supporting the primary findings.

**Fixed Effects of Laser Thresholds and Condition on RR Frequency at Peak Power of the PSD**

| **Effect** | **Estimate** | **Std. Error** | **df** | **t-value** | **p-value** |
| --- | --- | --- | --- | --- | --- |
| Intercept (Baseline) | 1.785e-02 | 2.445e-03 | 12.45 | 7.300 | 7.70e-06 *** |
| -15 vs Baseline | 6.957e-04 | 4.243e-03 | 304.3 | 0.164 | 0.8699 |
| -10 vs Baseline | 3.834e-03 | 4.243e-03 | 305.7 | 0.904 | 0.3669 |
| -5 vs Baseline | 1.355e-02 | 4.193e-03 | 306.2 | 3.232 | 0.0014 ** |
| 0 vs Baseline | 2.273e-02 | 4.276e-03 | 309.1 | 5.316 | 2.04e-07 *** |
| 5 vs Baseline | 2.532e-02 | 4.776e-03 | 308.9 | 5.302 | 2.19e-07 *** |
| YFP vs ChR2 | -9.377e-04 | 3.917e-03 | 13.10 | -0.239 | 0.8145 |
| -15 vs Baseline * YFP | -2.909e-03 | 6.725e-03 | 303.9 | -0.432 | 0.6657 |
| -10 vs Baseline * YFP | 1.569e-03 | 6.323e-03 | 304.6 | 0.248 | 0.8041 |
| -5 vs Baseline * YFP | -1.206e-02 | 6.449e-03 | 304.7 | -1.870 | 0.0624 . |
| 0 vs Baseline * YFP | -2.243e-02 | 6.304e-03 | 307.5 | -3.559 | 0.0004 *** |
| 5 vs Baseline * YFP | -3.052e-02 | 7.552e-03 | 307.2 | -4.041 | 6.73e-05 *** |

**Pairwise Comparisons with Bootstrapped Confidence Intervals**

| **Contrast** | **Estimate** | **SE** | **df** | **t-ratio** | **p-value** | **Lower Bound (2.5%)** | **Upper Bound (97.5%)** |
| --- | --- | --- | --- | --- | --- | --- | --- |
| **Within ChR2** |  |  |  |  |  |  |  |
| Baseline vs -15 | -0.000696 | 0.00424 | 7.21e+07 | -0.164 | 1.0000 | -6.06e-03 | 5.41e-03 |
| Baseline vs -10 | -0.003834 | 0.00424 | 1.26e+08 | -0.904 | 0.9457 | -3.03e-03 | 1.08e-02 |
| Baseline vs -5 | -0.013550 | 0.00419 | 1.45e+08 | -3.231 | 0.0156 * | -2.14e-03 | 1.21e-02 |
| Baseline vs 0 | -0.022733 | 0.00428 | 1.36e+08 | -5.316 | <.0001 *** | 1.41e-03 | 2.88e-02 |
| Baseline vs 5 | -0.025322 | 0.00478 | 7.86e+07 | -5.302 | <.0001 *** | 4.60e-02 | 4.60e-02 |
| -15 vs -10 | -0.003139 | 0.00553 | 9.26e+07 | -0.568 | 0.9931 | -6.91e-03 | 6.89e-03 |
| -15 vs -5 | -0.012854 | 0.00549 | 9.89e+07 | -2.340 | 0.1782 | -6.91e-03 | 6.89e-03 |
| -15 vs 0 | -0.022037 | 0.00554 | 9.73e+07 | -3.975 | 0.0010 ** | 1.50e-03 | 4.21e-02 |
| -15 vs 5 | -0.024627 | 0.00593 | 7.36e+07 | -4.152 | 0.0005 *** | 1.50e-03 | 4.21e-02 |
| -10 vs -5 | -0.009716 | 0.00550 | 1.41e+08 | -1.766 | 0.4880 | -2.43e-02 | 2.39e-02 |
| -10 vs 0 | -0.018898 | 0.00552 | 1.38e+08 | -3.426 | 0.0081 ** | -3.32e-02 | 1.66e-02 |
| -10 vs 5 | -0.021488 | 0.00591 | 9.62e+07 | -3.634 | 0.0038 ** | -3.32e-02 | 1.66e-02 |
| -5 vs 0 | -0.009183 | 0.00554 | 1.48e+08 | -1.658 | 0.5595 | -3.32e-02 | 1.66e-02 |
| -5 vs 5 | -0.011772 | 0.00592 | 1.00e+08 | -1.987 | 0.3495 | -3.32e-02 | 1.66e-02 |
| 0 vs 5 | -0.002590 | 0.00590 | 1.02e+08 | -0.439 | 0.9979 | -3.32e-02 | 1.66e-02 |
| **Within YFP** |  |  |  |  |  |  |  |
| Baseline vs -15 | 0.002213 | 0.00522 | 1.00e+08 | 0.424 | 0.9983 | -4.20e-02 | 6.84e-05 |
| Baseline vs -10 | -0.005404 | 0.00469 | 9.59e+07 | -1.153 | 0.8591 | -3.89e-02 | -6.65e-03 |
| Baseline vs -5 | -0.001489 | 0.00490 | 8.82e+07 | -0.304 | 0.9997 | -3.89e-02 | -6.65e-03 |
| Baseline vs 0 | -0.000301 | 0.00463 | 1.01e+08 | -0.065 | 1.0000 | -3.89e-02 | -6.65e-03 |
| Baseline vs 5 | 0.005198 | 0.00585 | 1.06e+08 | 0.888 | 0.9494 | -3.89e-02 | -6.65e-03 |
| -15 vs -10 | -0.007617 | 0.00637 | 9.94e+07 | -1.196 | 0.8390 | -3.89e-02 | -6.65e-03 |
| -15 vs -5 | -0.003702 | 0.00653 | 9.52e+07 | -0.567 | 0.9931 | -3.89e-02 | -6.65e-03 |
| -15 vs 0 | -0.002514 | 0.00632 | 1.03e+08 | -0.398 | 0.9987 | -3.89e-02 | -6.65e-03 |
| -15 vs 5 | 0.002985 | 0.00726 | 1.05e+08 | 0.411 | 0.9985 | -3.89e-02 | -6.65e-03 |
| -10 vs -5 | 0.003914 | 0.00610 | 9.24e+07 | 0.641 | 0.9879 | -3.89e-02 | -6.65e-03 |
| -10 vs 0 | 0.005103 | 0.00589 | 9.99e+07 | 0.867 | 0.9544 | -3.89e-02 | -6.65e-03 |
| -10 vs 5 | 0.010601 | 0.00688 | 1.03e+08 | 1.541 | 0.6376 | -3.89e-02 | -6.65e-03 |
| -5 vs 0 | 0.001188 | 0.00606 | 9.50e+07 | 0.196 | 1.0000 | -3.89e-02 | -6.65e-03 |
| -5 vs 5 | 0.006687 | 0.00703 | 1.00e+08 | 0.951 | 0.9330 | -3.89e-02 | -6.65e-03 |
| 0 vs 5 | 0.005499 | 0.00682 | 1.08e+08 | 0.806 | 0.9665 | -3.89e-02 | -6.65e-03 |
| **ChR2 vs YFP** |  |  |  |  |  |  |  |
| Baseline | 0.000938 | 0.00392 | 433947 | 0.239 | 0.8110 | -3.89e-02 | -6.65e-03 |
| Threshold -15 | 0.003846 | 0.00682 | 3666166 | 0.564 | 0.5730 | -3.89e-02 | -6.65e-03 |
| Threshold -10 | -0.000632 | 0.00643 | 3308704 | -0.098 | 0.9217 | -3.89e-02 | -6.65e-03 |
| Threshold -5 | 0.012998 | 0.00656 | 3594680 | 1.983 | 0.0474 * | -3.89e-02 | -6.65e-03 |
| Threshold 0 | 0.023369 | 0.00639 | 3429477 | 3.657 | 0.0003 *** | -3.89e-02 | -6.65e-03 |
| Threshold 5 | 0.031458 | 0.00762 | 5780300 | 4.130 | <.0001 *** | -3.89e-02 | -6.65e-03 |

###### 2i: RR PSD peak power

The analysis assessed the effects of laser threshold levels on RR PSD peak power, comparing the conditions ChR2 and YFP. The results showed no significant effects of laser threshold levels on RR PSD peak power within the ChR2 condition or YFP condition.

Block bootstrapping based on Subject ID was used due to issues with normality, heteroscedasticity, and slight autocorrelation of residuals, ensuring robust CIs. These intervals confirmed that there was no effect of the manipulation.

**Fixed Effects of Laser Thresholds and Condition on RR PSD Peak Power**

| **Effect** | **Estimate** | **Std. Error** | **df** | **t-value** | **p-value** |
| --- | --- | --- | --- | --- | --- |
| Intercept (Baseline) | 7.721e-04 | 9.594e-05 | 9.571 | 8.048 | 1.46e-05 *** |
| -15 vs Baseline | -6.556e-05 | 1.068e-04 | 303.5 | -0.614 | 0.540 |
| -10 vs Baseline | 2.254e-05 | 1.069e-04 | 304.1 | 0.211 | 0.833 |
| -5 vs Baseline | 7.573e-05 | 1.057e-04 | 304.3 | 0.717 | 0.474 |
| 0 vs Baseline | 4.164e-05 | 1.080e-04 | 305.7 | 0.385 | 0.700 |
| 5 vs Baseline | -8.995e-05 | 1.206e-04 | 305.6 | -0.746 | 0.456 |
| YFP vs ChR2 | -6.069e-05 | 1.525e-04 | 9.785 | -0.398 | 0.699 |
| -15 vs Baseline * YFP | 2.373e-04 | 1.692e-04 | 303.4 | 1.402 | 0.162 |
| -10 vs Baseline * YFP | -4.665e-05 | 1.592e-04 | 303.6 | -0.293 | 0.770 |
| -5 vs Baseline * YFP | -4.848e-05 | 1.624e-04 | 303.7 | -0.299 | 0.765 |
| 0 vs Baseline * YFP | -2.232e-05 | 1.590e-04 | 304.9 | -0.140 | 0.888 |
| 5 vs Baseline * YFP | 8.291e-05 | 1.905e-04 | 304.7 | 0.435 | 0.664 |

**Pairwise Comparisons with Bootstrapped Confidence Intervals**

| **Contrast** | **Estimate** | **SE** | **df** | **t-ratio** | **p-value** | **Lower Bound (2.5%)** | **Upper Bound (97.5%)** |
| --- | --- | --- | --- | --- | --- | --- | --- |
| **Within ChR2** |  |  |  |  |  |  |  |
| Baseline vs -15 | 3.14e-05 | 7.18e-05 | 73600 | 0.438 | 0.9980 | -1.73e-04 | 3.44e-04 |
| Baseline vs -10 | -4.57e-06 | 7.24e-05 | 80700 | -0.063 | 1.0000 | -4.56e-04 | 1.48e-04 |
| Baseline vs -5 | -8.83e-05 | 7.45e-05 | 122000 | -1.186 | 0.8440 | -1.56e-04 | 3.36e-04 |
| Baseline vs 0 | 6.56e-05 | 0.000107 | 7.19e+07 | 0.614 | 0.9901 | -3.74e-04 | 5.06e-04 |
| Baseline vs 5 | -2.25e-05 | 0.000107 | 1.25e+08 | -0.211 | 0.9999 | -3.97e-04 | 4.17e-04 |
| -15 vs -10 | -7.57e-05 | 0.000106 | 1.44e+08 | -0.717 | 0.9800 | -2.39e-04 | 1.30e-04 |
| -15 vs -5 | -4.16e-05 | 0.000108 | 1.34e+08 | -0.385 | 0.9989 | -4.55e-04 | 3.07e-04 |
| -15 vs 0 | 9.00e-05 | 0.000121 | 7.74e+07 | 0.746 | 0.9762 | -4.44e-04 | 7.02e-05 |
| -15 vs 5 | -8.81e-05 | 0.000139 | 9.25e+07 | -0.634 | 0.9885 | -1.69e-04 | 2.44e-04 |
| -10 vs -5 | -1.41e-04 | 0.000138 | 9.88e+07 | -1.022 | 0.9109 | -4.20e-04 | 3.22e-04 |
| -10 vs 0 | -1.07e-04 | 0.000140 | 9.69e+07 | -0.767 | 0.9729 | -3.95e-04 | 3.07e-04 |
| -10 vs 5 | 2.44e-05 | 0.000149 | 7.33e+07 | 0.163 | 1.0000 | -1.06e-04 | 3.67e-04 |
| -5 vs 0 | -5.32e-05 | 0.000139 | 1.40e+08 | -0.384 | 0.9989 | -3.95e-04 | 3.08e-04 |
| -5 vs 5 | -1.91e-05 | 0.000139 | 1.38e+08 | -0.138 | 1.0000 | -1.80e-04 | 3.68e-04 |
| 0 vs 5 | 1.12e-04 | 0.000149 | 9.57e+07 | 0.756 | 0.9747 | -1.23e-04 | 2.66e-04 |
| **Within YFP** |  |  |  |  |  |  |  |
| Baseline vs -15 | -1.60e-04 | 8.98e-05 | 94100 | -1.780 | 0.4787 | -4.74e-04 | 1.89e-04 |
| Baseline vs -10 | -1.72e-04 | 0.000131 | 1.00e+08 | -1.308 | 0.7809 | -1.37e-04 | 3.90e-04 |
| Baseline vs -5 | 2.41e-05 | 0.000118 | 9.58e+07 | 0.204 | 1.0000 | -4.75e-04 | 1.89e-04 |
| Baseline vs 0 | -2.73e-05 | 0.000123 | 8.81e+07 | -0.221 | 0.9999 | -4.20e-04 | 3.68e-04 |
| Baseline vs 5 | -1.93e-05 | 0.000117 | 1.00e+08 | -0.166 | 1.0000 | -3.97e-04 | 5.70e-04 |
| -15 vs -10 | 7.04e-06 | 0.000147 | 1.05e+08 | 0.048 | 1.0000 | -1.00e-04 | 4.81e-04 |
| -15 vs -5 | 1.96e-04 | 0.000160 | 9.93e+07 | 1.222 | 0.8263 | -1.31e-04 | 4.00e-04 |
| -15 vs 0 | 1.44e-04 | 0.000164 | 9.50e+07 | 0.879 | 0.9516 | -1.13e-04 | 5.24e-04 |
| -15 vs 5 | 1.52e-04 | 0.000159 | 1.03e+08 | 0.958 | 0.9309 | -1.37e-04 | 4.00e-04 |
| -10 vs -5 | 1.79e-04 | 0.000183 | 1.05e+08 | 0.978 | 0.9249 | -4.75e-04 | 1.89e-04 |
| -10 vs 0 | -5.14e-05 | 0.000154 | 9.24e+07 | -0.335 | 0.9994 | -4.75e-04 | 1.89e-04 |
| -10 vs 5 | -4.34e-05 | 0.000148 | 9.97e+07 | -0.293 | 0.9997 | -1.20e-04 | 1.34e-05 |
| -5 vs 0 | -1.71e-05 | 0.000173 | 1.03e+08 | -0.099 | 1.0000 | -4.75e-04 | 1.34e-05 |
| -5 vs 5 | 7.93e-06 | 0.000153 | 9.47e+07 | 0.052 | 1.0000 | -3.45e-04 | 4.00e-04 |
| 0 vs 5 | 3.43e-05 | 0.000177 | 9.98e+07 | 0.194 | 1.0000 | -4.75e-04 | 4.00e-04 |
| **ChR2 vs YFP** |  |  |  |  |  |  |  |
| Threshold 0 | 6.07e-05 | 0.000153 | 41888 | 0.397 | 0.6917 | -4.75e-04 | 3.40e-04 |
| Threshold -15 | -1.77e-04 | 0.000208 | 140058 | -0.850 | 0.3953 | -4.75e-04 | 1.89e-04 |
| Threshold -10 | 1.07e-04 | 0.000200 | 124803 | 0.538 | 0.5906 | -4.75e-04 | 1.89e-04 |
| Threshold -5 | 1.09e-04 | 0.000202 | 131908 | 0.540 | 0.5894 | -4.75e-04 | 1.34e-05 |
| Threshold 0 | 8.30e-05 | 0.000199 | 126195 | 0.417 | 0.6766 | -4.75e-04 | 4.00e-04 |
| Threshold 5 | -2.22e-05 | 0.000225 | 195784 | -0.099 | 0.9212 | -4.75e-04 | 4.00e-04 |

##### Figure 3: Heart rate decelerations

3b: RR (SD)

The analysis assessed the effects of laser threshold levels on RR SD for CHR2 animals. The results showed an initial decrease in RR SD as thresholds progressed but an eventual increase as threshold reached highest levels.

Block bootstrapping based on Subject ID was used due to issues with heteroscedasticity and slight autocorrelation of residuals, ensuring robust CIs.

**Fixed Effects of Laser Thresholds on RR (SD)**

| **Effect** | **Estimate** | **Std. Error** | **df** | **t-value** | **p-value** |
| --- | --- | --- | --- | --- | --- |
| Intercept (-15) | 1.106 e-02 | 6.631 e-04 | 5.969 | 16.683 | 3.11 e-06 *** |
| -10 vs -15 | -1.239 e-03 | 2.358 e-04 | 2478 | -5.255 | 1.61 e-07 |
| -5 vs -15 | -1.533 e-03 | 2.143 e-04 | 2479 | -7.151 | 1.13 e-12 |
| 0 vs -15 | -1.040 e-03 | 2.056 e-04 | 2479 | -5.059 | 4.52 e-07 |
| 5 vs -15 | -9.873 e-04 | 2.090 e-04 | 2479 | -4.725 | 2.43 e-06 |

**Pairwise Comparisons with Bootstrapped Confidence Intervals (Turkey adjusted)**

| **Contrast** | **Estimate** | **SE** | **df** | **t-ratio** | **p-value** | **Lower Bound (2.5 %)** | **Upper Bound (97.5 %)** |
| --- | --- | --- | --- | --- | --- | --- | --- |
| -15 vs -10 | 1.24 e-03 | 2.36 e-04 | 2478 | 5.255 | < .0001 | 5.91 e-04 | 1.76 e-03 |
| -15 vs -5 | 1.53 e-03 | 2.14 e-04 | 2479 | 7.151 | < .0001 | 2.92 e-04 | 2.19 e-03 |
| -15 vs 0 | 1.04 e-03 | 2.06 e-04 | 2479 | 5.059 | < .0001 | -2.09 e-04 | 1.52 e-03 |
| -15 vs 5 | 9.87 e-04 | 2.09 e-04 | 2479 | 4.725 | < .0001 | 3.13 e-04 | 1.24 e-03 |
| -10 vs -5 | 2.94 e-04 | 1.60 e-04 | 2479 | 1.835 | 0.3535 | -1.00 e-03 | 1.35 e-03 |
| -10 vs 0 | -1.99 e-04 | 1.47 e-04 | 2478 | -1.351 | 0.6593 | -1.31 e-03 | 6.11 e-04 |
| -10 vs 5 | -2.52 e-04 | 1.51 e-04 | 2478 | -1.666 | 0.4552 | -8.60 e-04 | 2.26 e-04 |
| -5 vs 0 | -4.93 e-04 | 1.11 e-04 | 2480 | -4.426 | 0.0001 | -1.29 e-03 | 6.17 e-04 |
| -5 vs 5 | -5.46 e-04 | 1.15 e-04 | 2480 | -4.743 | < .0001 | -1.19 e-03 | 2.32 e-04 |
| 0 vs 5 | -5.27 e-05 | 9.29 e-05 | 2478 | -0.567 | 0.9797 | -7.73 e-04 | 7.93 e-04 |

3b: RR Sample Entropy (10 s after LC stimulation)

The analysis assessed the effects of laser threshold levels on sample entropy for CHR2 animals. The results showed no significant differences between thresholds.

Block bootstrapping based on Subject ID was used due to issues with normality, heteroscedasticity, and slight autocorrelation of residuals, ensuring robust CIs.

**Fixed Effects of Laser Thresholds on RR sample entrophy**

| **Effect** | **Estimate** | **Std. Error** | **df** | **t-value** | **p-value** |
| --- | --- | --- | --- | --- | --- |
| Intercept (-15) | 2.365e-01 | 8.088e-02 | 5.086e+02 | 2.924 | 0.00361 ** |
| -10 vs -15 | -9.063e-02 | 9.718e-02 | 2.212e+03 | -0.933 | 0.35110 |
| -5 vs -15 | -9.317e-02 | 8.793e-02 | 1.975e+03 | -1.060 | 0.28948 |
| 0 vs -15 | -1.140e-02 | 8.422e-02 | 1.821e+03 | -0.135 | 0.89234 |
| 5 vs -15 | 6.041e-03 | 8.524e-02 | 1.472e+03 | 0.071 | 0.94350 |

**Pairwise Comparisons with Bootstrapped Confidence Intervals (Turkey adjusted)**

| **Contrast** | **Estimate** | **SE** | **df** | **t-ratio** | **p-value** | **Lower Bound (2.5 %)** | **Upper Bound (97.5 %)** |
| --- | --- | --- | --- | --- | --- | --- | --- |
| -15 vs -10 | 9.063e-02 | 9.800e-02 | 2.191e+03 | 0.925 | 0.8872 | -6.809e-02 | 1.159e+00 |
| -15 vs -5 | 9.317e-02 | 8.880e-02 | 1.938e+03 | 1.049 | 0.8325 | -6.402e-02 | 1.106e+00 |
| -15 vs 0 | 1.140e-02 | 8.540e-02 | 1.776e+03 | 0.134 | 0.9999 | -1.764e-01 | 1.122e+00 |
| -15 vs 5 | -6.041e-03 | 8.680e-02 | 1.416e+03 | -0.070 | 1.0000 | -1.899e-01 | 1.077e+00 |
| -10 vs -5 | 2.530e-03 | 6.650e-02 | 2.156e+03 | 0.038 | 1.0000 | -3.968e-02 | 2.918e-02 |
| -10 vs 0 | -7.923e-02 | 6.120e-02 | 2.339e+03 | -1.294 | 0.6951 | -1.293e-01 | -2.701e-02 |
| -10 vs 5 | -9.667e-02 | 6.290e-02 | 1.981e+03 | -1.538 | 0.5381 | -1.450e-01 | -3.762e-02 |
| -5 vs 0 | -8.177e-02 | 4.570e-02 | 966 | -1.789 | 0.3808 | -1.340e-01 | -1.028e-02 |
| -5 vs 5 | -9.921e-02 | 4.750e-02 | 926 | -2.088 | 0.2261 | -1.472e-01 | -2.525e-02 |
| 0 vs 5 | -1.744e-02 | 3.860e-02 | 2.289e+03 | -0.451 | 0.9914 | -3.519e-02 | 4.124e-03 |

3c: RR mean

The analysis assessed the effects of laser threshold levels on mean RR intervals for CHR2 animals. The results showed significantly lower values at higher thresholds.

Block bootstrapping based on Subject ID was used due to issues with heteroscedasticity and slight autocorrelation of residuals, ensuring robust CIs.

**Fixed Effects of Laser Thresholds on mean RR**

| **Effect** | **Estimate** | **Std. Error** | **df** | **t-value** | **p-value** |
| --- | --- | --- | --- | --- | --- |
| Intercept (-15) | 1.262 e-01 | 3.270 e-03 | 517.4 | 38.600 | 1.44e-07 *** |
| -10 vs -15 | 3.914 e-04 | 5.211 e-04 | 2478 | 0.751 | 0.453 |
| -5 vs -15 | –5.366 e-04 | 4.738 e-04 | 2478 | –1.133 | 0.258 |
| 0 vs -15 | –6.073 e-04 | 4.544 e-04 | 2478 | –1.336 | 0.182 |
| 5 vs -15 | –1.879 e-03 | 4.619 e-04 | 2478 | –4.068 | 4.90 e-05 ** |

**Pairwise Comparisons with Bootstrapped Confidence Intervals (Turkey adjusted)**

| **Contrast** | **Estimate** | **SE** | **df** | **t-ratio** | **p-value** | **Lower Bound (2.5 %)** | **Upper Bound (97.5 %)** |
| --- | --- | --- | --- | --- | --- | --- | --- |
| -15 vs -10 | –3.91 e-04 | 9.80 e-02 | 2191 | –0.751 | 0.9443 | –2.278 e-03 | 4.871 e-03 |
| -15 vs -5 | 5.37 e-04 | 8.88 e-02 | 1938 | 1.049 | 0.7893 | –2.038 e-03 | 7.136 e-03 |
| -15 vs 0 | 6.07 e-04 | 8.54 e-02 | 1776 | 1.336 | 0.6684 | –1.218 e-03 | 5.709 e-03 |
| -15 vs 5 | 1.88 e-03 | 8.68 e-02 | 1416 | 4.067 | 0.0005 | –1.233 e-03 | 7.086 e-03 |
| -10 vs -5 | 9.28 e-04 | 6.65 e-02 | 2478 | 2.620 | 0.0671 | –2.984 e-03 | 4.786 e-03 |
| -10 vs 0 | 9.99 e-04 | 6.12 e-02 | 2478 | 3.070 | 0.0184 | –2.754 e-03 | 4.002 e-03 |
| -10 vs 5 | 2.27 e-03 | 6.29 e-02 | 2478 | 6.804 | < .0001 | –1.940 e-03 | 5.920 e-03 |
| -5 vs 0 | 7.07 e-05 | 4.57 e-02 | 966 | 0.287 | 0.9985 | –2.197 e-03 | 2.095 e-03 |
| -5 vs 5 | 1.34 e-03 | 4.75 e-02 | 926 | 5.278 | < .0001 | –1.501 e-03 | 3.808 e-03 |
| 0 vs 5 | 1.27 e-03 | 3.86 e-02 | 2289 | 6.189 | < .0001 | –5.180 e-04 | 3.424 e-03 |

3c: mean BPM

The analysis assessed the effects of laser threshold levels on mean BPM (heart rate) for CHR2 animals. The results showed significantly higher values at higher thresholds.

Block bootstrapping based on Subject ID was used due to issues with heteroscedasticity and autocorrelation of residuals, ensuring robust CIs.

**Fixed Effects of Laser Thresholds on mean BPM**

| **Effect** | **Estimate** | **Std. Error** | **df** | **t-value** | **p-value** |
| --- | --- | --- | --- | --- | --- |
| Intercept (–15) | 480.5947 | 12.9597 | 5.1925 | 37.084 | 1.69 e-07 *** |
| –10 vs –15 | –0.9836 | 2.1712 | 2478.0696 | –0.453 | 0.651 |
| –5 vs –15 | 2.8136 | 1.9741 | 2478.1408 | 1.425 | 0.154 |
| 0 vs –15 | 2.2554 | 1.8933 | 2478.1167 | 1.191 | 0.234 |
| 5 vs –15 | 8.3729 | 1.9244 | 2478.1490 | 4.351 | 1.41 e-05 *** |

**Pairwise Comparisons with Bootstrapped Confidence Intervals (Turkey adjusted)**

| **Contrast** | **Estimate** | **SE** | **df** | **t-ratio** | **p-value** | **Lower Bound (2.5 %)** | **Upper Bound (97.5 %)** |
| --- | --- | --- | --- | --- | --- | --- | --- |
| –15 vs –10 | 0.9836 | 2.1712 | 2478 | 0.453 | 0.9913 | –19.93 | 8.53 |
| –15 vs –5 | –2.8140 | 1.9741 | 2478 | –1.425 | 0.6113 | –29.44 | 7.74 |
| –15 vs 0 | –2.2554 | 1.8933 | 2478 | –1.336 | 0.6684 | –22.46 | 4.48 |
| –15 vs 5 | –8.3729 | 1.9244 | 2478 | –4.351 | 0.0001 | –29.36 | 4.22 |
| –10 vs –5 | –3.7970 | 1.4760 | 2478 | –2.573 | 0.0757 | –20.44 | 12.20 |
| –10 vs 0 | –3.2390 | 1.3550 | 2478 | –2.390 | 0.1183 | –14.88 | 11.41 |
| –10 vs 5 | –9.3570 | 1.3900 | 2478 | –6.731 | < .0001 | –24.21 | 7.71 |
| –5 vs 0 | 0.5580 | 1.0250 | 2478 | 0.544 | 0.9826 | –7.69 | 10.29 |
| –5 vs 5 | –5.5590 | 1.0590 | 2478 | –5.248 | < .0001 | –15.92 | 6.67 |
| 0 vs 5 | –6.1170 | 0.8560 | 2478 | –7.147 | < .0001 | –15.63 | 1.00 |

3d: Correlation between heart rate (BPM) and RR (SD)

The analysis assessed the relationship between mean BPM (heart rate) and the standard deviation of RR intervals for CHR2 animals. The results showed a positive relationship.

Block bootstrapping based on Subject ID was used due to issues with normality, heteroscedasticity, and autocorrelation of residuals, ensuring robust CIs.

| **Statistic** | **Value** |
| --- | --- |
| **Fixed Effects Estimates** |  |
| Intercept Estimate | 461.558 |
| Intercept SE | 11.866 |
| Intercept df | 5.231 |
| Intercept t-value | 38.90 |
| Intercept p-value | 1.21 × 10⁻⁷ *** |
| RR_SD Estimate | 2 307.431 |
| RR_SD SE | 179.666 |
| RR_SD df | 2 482.956 |
| RR_SD t-value | 12.84 |
| RR_SD p-value | < 2 × 10⁻¹⁶ *** |
| **Random Effects** |  |
| SubjectID (Intercept) Variance | 824.8 |
| SubjectID (Intercept) SD | 28.72 |
| Residual Variance | 293.4 |
| Residual SD | 17.13 |
| **R-squared Values** |  |
| Marginal R-squared (R²ₘ) | 0.0265 |
| Bootstrapped 2.5 % CI | 0.01992 |
| Bootstrapped 97.5 % CI | 0.04030 |
| Conditional R-squared (R²_c) | 0.7446 |
| Bootstrapped 2.5 % CI | 0.72505 |
| Bootstrapped 97.5 % CI | 0.76315 |
| **Correlation of Fixed Effects** |  |
| Correlation (Intercept, RR_SD) | –0.151 |
| **Fixed Effects Estimates** |  |
| Intercept Estimate | 461.558 |

3g: RR value at HR decelerations (RR peak)

The analysis assessed the effects of laser threshold levels on the peak RR value (representing the heart rate decelerations) for CHR2 animals. The results showed significantly higher values at higher thresholds.

Block bootstrapping based on Subject ID was used due to issues with heteroscedasticity and autocorrelation of residuals, ensuring robust CIs.

**Fixed Effects of Laser Thresholds on RR peak**

| **Effect** | **Estimate** | **Std. Error** | **df** | **t-value** | **p-value** |
| --- | --- | --- | --- | --- | --- |
| Intercept (–15) | 1.297 × 10⁻¹ | 3.479 × 10⁻³ | 5.567 | 37.291 | 6.79 × 10⁻⁸ *** |
| –10 vs –15 | 4.257 × 10⁻⁶ | 9.755 × 10⁻⁴ | 2 478 | 0.004 | 0.9965 |
| –5 vs –15 | 5.418 × 10⁻⁴ | 8.870 × 10⁻⁴ | 2 478 | 0.611 | 0.5413 |
| 0 vs –15 | 3.953 × 10⁻³ | 8.507 × 10⁻⁴ | 2 478 | 4.647 | 3.54 × 10⁻⁶ *** |
| 5 vs –15 | 2.882 × 10⁻³ | 8.647 × 10⁻⁴ | 2 478 | 3.333 | 8.72 × 10⁻⁴ ** |

**Pairwise Comparisons with Bootstrapped Confidence Intervals (Turkey adjusted)**

| **Contrast** | **Estimate** | **SE** | **df** | **t-ratio** | **p-value** | **Lower Bound (2.5 %)** | **Upper Bound (97.5 %)** |
| --- | --- | --- | --- | --- | --- | --- | --- |
| –15 vs –10 | –4.26 e-06 | 9.76 e-04 | 2 478 | –0.004 | 1.0000 | –0.003605 | 0.003144 |
| –15 vs –5 | –5.42 e-04 | 8.87 e-04 | 2 478 | –0.611 | 0.9734 | –0.006521 | 0.003345 |
| –15 vs 0 | –3.95 e-03 | 8.51 e-04 | 2 478 | –4.647 | < .0001 | –0.008918 | –0.001932 |
| –15 vs 5 | –2.88 e-03 | 8.65 e-04 | 2 478 | –3.333 | 0.0078 | –0.007573 | –0.000276 |
| –10 vs –5 | –5.38 e-04 | 6.63 e-04 | 2 478 | –0.811 | 0.9274 | –0.005907 | 0.004343 |
| –10 vs 0 | –3.95 e-03 | 6.09 e-04 | 2 478 | –6.485 | < .0001 | –0.009089 | –0.001618 |
| –10 vs 5 | –2.88 e-03 | 6.25 e-04 | 2 478 | –4.607 | < .0001 | –0.007530 | 0.000248 |
| –5 vs 0 | –3.41 e-03 | 4.61 e-04 | 2 479 | –7.404 | < .0001 | –0.007849 | 0.000265 |
| –5 vs 5 | –2.34 e-03 | 4.76 e-04 | 2 479 | –4.916 | < .0001 | –0.005621 | 0.000925 |
| 0 vs 5 | 1.07 e-03 | 3.85 e-04 | 2 478 | 2.786 | 0.0428 | –0.000220 | 0.003070 |

3h: Heart rate (BPM) at HR decelerations

The analysis assessed the effects of laser threshold levels on the heart rate (BPM) at the heart rate decelerations for CHR2 animals. The results showed significantly lower values at higher thresholds.

Block bootstrapping based on Subject ID was used due to issues with heteroscedasticity and slight autocorrelation of residuals, ensuring robust CIs.

**Fixed Effects of Laser Thresholds on RR peak**

| **Effect** | **Estimate** | **Std. Error** | **df** | **t-value** | **p-value** |
| --- | --- | --- | --- | --- | --- |
| Intercept (–15) | 467.599 | 12.617 | 5.6437 | 37.060 | 5.87 × 10⁻⁸ *** |
| –10 vs –15 | –0.996 | 3.753 | 2478.23 | –0.265 | 0.7907 |
| –5 vs –15 | –2.213 | 3.412 | 2478.45 | –0.649 | 0.5167 |
| 0 vs –15 | –15.840 | 3.272 | 2478.38 | –4.841 | 1.37 × 10⁻⁶ *** |
| 5 vs –15 | –11.900 | 3.326 | 2478.48 | –3.578 | 0.000353 *** |

**Pairwise Comparisons with Bootstrapped Confidence Intervals (Turkey adjusted)**

| **Contrast** | **Estimate** | **SE** | **df** | **t-ratio** | **p-value** | **Lower Bound (2.5 %)** | **Upper Bound (97.5 %)** |
| --- | --- | --- | --- | --- | --- | --- | --- |
| –15 vs –10 | 0.996 | 3.752 | 2478 | 0.265 | 0.9989 | –10.876 | 15.649 |
| –15 vs –5 | 2.213 | 3.412 | 2478 | 0.649 | 0.9669 | –12.917 | 23.952 |
| –15 vs 0 | 15.840 | 3.272 | 2478 | 4.841 | < .0001 | 7.359 | 39.426 |
| –15 vs 5 | 11.900 | 3.326 | 2478 | 3.578 | 0.0032 | 3.055 | 30.208 |
| –10 vs –5 | 1.217 | 2.552 | 2479 | 0.477 | 0.9894 | –18.008 | 22.373 |
| –10 vs 0 | 14.844 | 2.343 | 2478 | 6.337 | < .0001 | 5.524 | 33.534 |
| –10 vs 5 | 10.904 | 2.400 | 2478 | 4.538 | 0.0001 | –1.153 | 29.082 |
| –5 vs 0 | 13.627 | 1.770 | 2480 | 7.689 | < .0001 | –0.957 | 31.123 |
| –5 vs 5 | 9.687 | 1.830 | 2479 | 5.291 | < .0001 | –3.585 | 23.221 |
| 0 vs 5 | –3.940 | 1.480 | 2478 | –2.663 | 0.0598 | –10.709 | 0.478 |

3i: RR amplitude

The analysis assessed the effects of laser threshold levels on RR amplitude leading up to heart rate decelerations for CHR2 animals. The results showed significantly higher values at higher thresholds. Additionally, the estimates were also compared to 0 to determine when the HR decelerations were statistically present between thresholds. As all model assumptions were met, no bootstrapping was performed.

**Fixed Effects of Laser Thresholds on RR amplitude**

| **Effect** | **Estimate** | **Std. Error** | **df** | **t-value** | **p-value** |
| --- | --- | --- | --- | --- | --- |
| Intercept (–15) | 1.089 × 10⁻³ | 1.665 × 10⁻³ | 14.02 | 0.654 | 0.524 |
| –10 vs –15 | –1.538 × 10⁻³ | 1.306 × 10⁻³ | 2480 | –1.178 | 0.239 |
| –5 vs –15 | 4.196 × 10⁻⁴ | 1.187 × 10⁻³ | 2482 | 0.353 | 0.724 |
| 0 vs –15 | 5.053 × 10⁻³ | 1.139 × 10⁻³ | 2481 | 4.438 | 9.50 × 10⁻⁶ *** |
| 5 vs –15 | 6.395 × 10⁻³ | 1.157 × 10⁻³ | 2482 | 5.525 | 3.64 × 10⁻⁸ *** |

**Pairwise Comparisons with Bootstrapped Confidence Intervals (Turkey adjusted)**

| **Contrast** | **Estimate** | **SE** | **df** | **t-ratio** | **p-value** |
| --- | --- | --- | --- | --- | --- |
| –15 vs –10 | 1.54 × 10⁻³ | 1.306 × 10⁻³ | 2480 | 1.178 | 0.7643 |
| –15 vs –5 | –4.20 × 10⁻⁴ | 1.188 × 10⁻³ | 2482 | –0.353 | 0.9967 |
| –15 vs 0 | –5.05 × 10⁻³ | 1.139 × 10⁻³ | 2481 | –4.436 | 0.0001 |
| –15 vs 5 | –6.39 × 10⁻³ | 1.158 × 10⁻³ | 2482 | –5.522 | < .0001 |
| –10 vs –5 | –1.96 × 10⁻³ | 8.88 × 10⁻⁴ | 2482 | –2.205 | 0.1780 |
| –10 vs 0 | –6.59 × 10⁻³ | 8.16 × 10⁻⁴ | 2480 | –8.083 | < .0001 |
| –10 vs 5 | –7.93 × 10⁻³ | 8.36 × 10⁻⁴ | 2481 | –9.484 | < .0001 |
| –5 vs 0 | –4.63 × 10⁻³ | 6.17 × 10⁻⁴ | 2479 | –7.512 | < .0001 |
| –5 vs 5 | –5.97 × 10⁻³ | 6.37 × 10⁻⁴ | 2482 | –9.376 | < .0001 |
| 0 vs 5 | –1.34 × 10⁻³ | 5.15 × 10⁻⁴ | 2480 | –2.604 | 0.0699 |

**Marginal means compared to zero**

| **Threshold** | **Estimate** | **SE** | **df** | **t-ratio** | **p-value** |
| --- | --- | --- | --- | --- | --- |
| –15 | 1.09 × 10⁻³ | 1.67 × 10⁻³ | 14.38 | 0.654 | 0.5237 |
| –10 | –4.50 × 10⁻⁴ | 1.46 × 10⁻³ | 8.53 | –0.308 | 0.7655 |
| –5 | 1.51 × 10⁻³ | 1.35 × 10⁻³ | 6.20 | 1.119 | 0.3048 |
| 0 | 6.14 × 10⁻³ | 1.31 × 10⁻³ | 5.50 | 4.692 | 0.0042 |
| 5 | 7.48 × 10⁻³ | 1.32 × 10⁻³ | 5.65 | 5.680 | 0.0016 |

3j: Correlation between heart rate (BPM) and RR amplitude

The analysis assessed the relationship between mean BPM (heart rate) and amplitude of RR for the heart rate decelerations for CHR2 animals. The results showed a positive relationship.

Block bootstrapping based on Subject ID was used due to issues with heteroscedasticity and autocorrelation of residuals, ensuring robust CIs.

| **Statistic** | **Value** |
| --- | --- |
| **Fixed Effects Estimates** |  |
| Intercept Estimate | 484.039 |
| Intercept SE | 12.620 |
| Intercept df | 5.002 |
| Intercept t-value | 38.356 |
| Intercept p-value | 2.26 × 10⁻⁷ *** |
| CorrRR_peak Estimate | 186.782 |
| CorrRR_peak SE | 39.188 |
| CorrRR_peak df | 1698.396 |
| CorrRR_peak t-value | 4.766 |
| CorrRR_peak p-value | 2.04 × 10⁻⁶ *** |
| **Random Effects** |  |
| SubjectID (Intercept) Variance | 953.7 |
| SubjectID (Intercept) SD | 30.88 |
| Residual Variance | 293.2 |
| Residual SD | 17.12 |
| **R-squared Values** |  |
| Marginal R-squared (R²ₘ) | 0.00333 |
| Bootstrapped 2.5 % CI | 0.00096 |
| Bootstrapped 97.5 % CI | 0.00695 |
| Conditional R-squared (R²꜀) | 0.76566 |
| Bootstrapped 2.5 % CI | 0.74586 |
| Bootstrapped 97.5 % CI | 0.78801 |
| **Correlation of Fixed Effects** |  |
| Correlation (Intercept, CorrRR_peak) | –0.021 |

3m: NE amplitude

The analysis assessed the effects of laser threshold levels on NE amplitudes leading up to the heart rate decelerations for CHR2 animals. The results showed significantly higher values at higher thresholds.

Block bootstrapping based on Subject ID was used due to issues with heteroscedasticity and slight autocorrelation of residuals, ensuring robust CIs.

**Fixed Effects of Laser Thresholds on NE amplitude**

| **Effect** | **Estimate** | **Std. Error** | **df** | **t-value** | **p-value** |
| --- | --- | --- | --- | --- | --- |
| Intercept (–15) | 3.04188 | 0.20344 | 7.4153 | 14.952 | 8.33 × 10⁻⁷ *** |
| –10 vs –15 | –0.28923 | 0.10547 | 2478.7701 | –2.742 | 0.00614 ** |
| –5 vs –15 | –1.22316 | 0.09589 | 2479.4918 | –12.756 | < 2 × 10⁻¹⁶ ** |
| 0 vs –15 | –1.95045 | 0.09196 | 2479.2580 | –21.209 | < 2 × 10⁻¹⁶ ** |
| 5 vs –15 | –2.06728 | 0.09347 | 2479.5792 | –22.116 | < 2 × 10⁻¹⁶ ** |

**Pairwise Comparisons with Bootstrapped Confidence Intervals (Turkey adjusted)**

| **Contrast** | **Estimate** | **SE** | **df** | **t-ratio** | **p-value** | **Lower (2.5 %)** | **Upper (97.5 %)** |
| --- | --- | --- | --- | --- | --- | --- | --- |
| –15 vs –10 | 0.289 | 0.1055 | 2479 | 2.742 | 0.0483 | –0.135 | 2.079 |
| –15 vs –5 | 1.223 | 0.0959 | 2480 | 12.755 | < 0.0001 | 0.970 | 4.202 |
| –15 vs 0 | 1.950 | 0.0920 | 2479 | 21.207 | < 0.0001 | 1.165 | 4.146 |
| –15 vs 5 | 2.067 | 0.0935 | 2480 | 22.113 | < 0.0001 | 0.484 | 1.188 |
| –10 vs –5 | 0.934 | 0.0717 | 2480 | 13.028 | < 0.0001 | 0.869 | 2.279 |
| –10 vs 0 | 1.661 | 0.0658 | 2479 | 25.232 | < 0.0001 | 1.053 | 2.342 |
| –10 vs 5 | 1.778 | 0.0675 | 2479 | 26.330 | < 0.0001 | 0.357 | 1.095 |
| –5 vs 0 | 0.727 | 0.0498 | 2482 | 14.601 | < 0.0001 | 0.562 | 1.204 |
| –5 vs 5 | 0.844 | 0.0515 | 2482 | 16.403 | < 0.0001 | –0.177 | 0.343 |
| 0 vs 5 | 0.117 | 0.0416 | 2479 | 2.810 | 0.0400 | –0.177 | 0.343 |

3n: mean NE values

The analysis assessed the effects of laser threshold levels on mean NE values leading up to the heart rate decelerations for CHR2 animals. The results showed significantly higher values at higher thresholds.

Block bootstrapping based on Subject ID was used due to issues with normality, heteroscedasticity, and autocorrelation of residuals, ensuring robust CIs.

**Fixed Effects of Laser Thresholds on mean NE**

| **Effect** | **Estimate** | **Std. Error** | **df** | **t-value** | **p-value** |
| --- | --- | --- | --- | --- | --- |
| Intercept (–15) | 1.5189 | 1.5462 | 5.1687 | 0.982 | 0.3696 |
| –10 vs –15 | 0.4710 | 0.2424 | 2478.0615 | 1.943 | 0.0521 . |
| –5 vs –15 | 1.3623 | 0.2204 | 2478.1239 | 6.181 | < .0001 *** |
| 0 vs –15 | 3.0204 | 0.2114 | 2478.1027 | 14.288 | < .0001 *** |
| 5 vs –15 | 1.9586 | 0.2149 | 2478.1310 | 9.115 | < .0001 *** |

**Pairwise Comparisons with Bootstrapped Confidence Intervals (Turkey adjusted)**

| **Contrast** | **Estimate** | **SE** | **df** | **t-ratio** | **p-value** | **Lower (2.5 %)** | **Upper (97.5 %)** |
| --- | --- | --- | --- | --- | --- | --- | --- |
| –15 vs –10 | –0.4710 | 0.2424 | 2478 | –1.943 | 0.2949 | –5.99 | 0.28 |
| –15 vs –5 | –1.3623 | 0.2204 | 2478 | –6.181 | < .0001 | –6.98 | –0.55 |
| –15 vs 0 | –3.0204 | 0.2114 | 2478 | –14.288 | < .0001 | –10.32 | –1.36 |
| –15 vs 5 | –1.9586 | 0.2149 | 2478 | –9.115 | < .0001 | –8.26 | –0.82 |
| –10 vs –5 | –0.8910 | 0.1648 | 2478 | –5.410 | < .0001 | –1.68 | 0.25 |
| –10 vs 0 | –2.5490 | 0.1513 | 2478 | –16.848 | < .0001 | –4.73 | –1.09 |
| –10 vs 5 | –1.4880 | 0.1552 | 2478 | –9.584 | < .0001 | –2.84 | –0.37 |
| –5 vs 0 | –1.6580 | 0.1145 | 2478 | –14.482 | < .0001 | –3.44 | –0.25 |
| –5 vs 5 | –0.5960 | 0.1183 | 2478 | –5.041 | < .0001 | –2.57 | 0.89 |
| 0 vs 5 | 1.0620 | 0.0956 | 2478 | 11.111 | < .0001 | –0.55 | 3.02 |

3o: Correlation between mean NE values and RR peak

The analysis assessed the relationship between mean NE values leading up to heart rate decelerations and the RR value at the heart rate decelerations for CHR2 animals. The results showed a positive relationship.

Block bootstrapping based on Subject ID was used due to issues with heteroscedasticity and slight autocorrelation of residuals, ensuring robust CIs.

| **Statistic** | **Value** |
| --- | --- |
| **Fixed Effects Estimates** |  |
| Intercept Estimate | 0.1302 |
| Intercept SE | 0.003112 |
| Intercept df | 5.061 |
| Intercept t-value | 41.83 |
| Intercept p-value | 2.26 × 10⁻⁷ ***(***) |
| MeanNE Estimate | 0.0005497 |
| MeanNE SE | 0.00007379 |
| MeanNE df | 2455 |
| MeanNE t-value | 7.45 |
| MeanNE p-value | 1.28 × 10⁻¹³ ***(***) |
| **Random Effects** |  |
| SubjectID (Intercept) Variance | 0.00005754 |
| SubjectID (Intercept) SD | 0.007586 |
| Residual Variance | 0.00006202 |
| Residual SD | 0.007875 |
| **R-squared Values** |  |
| Marginal R-squared (R²ₘ) | 0.04019 |
| Bootstrapped 2.5 % CI | 0.02003 |
| Bootstrapped 97.5 % CI | 0.07100 |
| Conditional R-squared (R²꜀) | 0.50212 |
| Bootstrapped 2.5 % CI | 0.47400 |
| Bootstrapped 97.5 % CI | 0.53600 |
| **Correlation of Fixed Effects** |  |
| Correlation (Intercept, MeanNE) | –0.084 |

3p: Sigma amplitude

The analysis assessed the effects of laser threshold levels on sigma amplitude leading up to the heart rate decelerations for CHR2 animals. The results showed significantly higher values at higher thresholds.

Block bootstrapping based on Subject ID was used due to issues with slight heteroscedasticity and slight autocorrelation of residuals, ensuring robust CIs.

**Fixed Effects of Laser Thresholds on sigma amplitude**

| **Effect** | **Estimate** | **Std. Error** | **df** | **t-value** | **p-value** |
| --- | --- | --- | --- | --- | --- |
| Intercept (–15) | –0.12429 | 0.08177 | 14.57763 | –1.520 | 0.1499 |
| –10 vs –15 | –0.01789 | 0.06486 | 2480.29899 | –0.276 | 0.7827 |
| –5 vs –15 | 0.09392 | 0.05896 | 2481.92269 | 1.593 | 0.1113 |
| 0 vs –15 | 0.11393 | 0.05655 | 2481.48177 | 2.015 | 0.0440 * |
| 5 vs –15 | 0.15781 | 0.05747 | 2482.11542 | 2.746 | 0.0061 ** |

**Pairwise Comparisons with Bootstrapped Confidence Intervals (Turkey adjusted)**

| **Contrast** | **Estimate** | **SE** | **df** | **t-ratio** | **p-value** | **Lower (2.5 %)** | **Upper (97.5 %)** |
| --- | --- | --- | --- | --- | --- | --- | --- |
| –15 vs –10 | 0.0179 | 0.0649 | 2478 | 0.276 | 0.9987 | –0.3819 | 0.1388 |
| –15 vs –5 | –0.0939 | 0.0590 | 2482 | –1.592 | 0.5024 | –0.5322 | –0.0161 |
| –15 vs 0 | –0.1139 | 0.0566 | 2482 | –2.014 | 0.2594 | –0.5788 | 0.0095 |
| –15 vs 5 | –0.1578 | 0.0575 | 2482 | –2.745 | 0.0480 | –0.6681 | –0.0331 |
| –10 vs –5 | –0.1118 | 0.0441 | 2482 | –2.536 | 0.0831 | –0.2735 | –0.0341 |
| –10 vs 0 | –0.1318 | 0.0405 | 2480 | –3.255 | 0.0101 | –0.2876 | –0.0045 |
| –10 vs 5 | –0.1757 | 0.0415 | 2481 | –4.230 | 0.0002 | –0.3674 | –0.0536 |
| –5 vs 0 | –0.0200 | 0.0306 | 2479 | –0.653 | 0.9661 | –0.0746 | 0.0285 |
| –5 vs 5 | –0.0639 | 0.0316 | 2481 | –2.019 | 0.2571 | –0.1590 | 0.0136 |
| 0 vs 5 | –0.0439 | 0.0256 | 2481 | –1.716 | 0.4243 | –0.0893 | –0.0035 |

3q: Correlation between sigma amplitude and RR peak

The analysis assessed the relationship between sigma amplitude leading up to heart rate decelerations and the RR value at the heart rate decelerations for CHR2 animals in the two highest LC stimulation frequency conditions. The results showed a positive relationship.

Block bootstrapping based on Subject ID was used due to slight issues with normality and autocorrelation of residuals, as well as heteroscedasticity, ensuring robust CIs.

| **Statistic** | **Value** |
| --- | --- |
| **Fixed Effects Estimates** |  |
| Intercept Estimate | 0.1322 |
| Intercept SE | 0.003403 |
| Intercept df | 4.996 |
| Intercept t-value | 38.839 |
| Intercept p-value | 2.15 × 10⁻⁷ *** |
| Sigma_ampl Estimate | 0.001607 |
| Sigma_ampl SE | 0.0003041 |
| Sigma_ampl df | 2482 |
| Sigma_ampl t-value | 5.285 |
| Sigma_ampl p-value | 1.36 × 10⁻⁷ *** |
| **Random Effects** |  |
| SubjectID (Intercept) Variance | 0.00006930 |
| SubjectID (Intercept) SD | 0.008325 |
| Residual Variance | 0.00006267 |
| Residual SD | 0.007917 |
| **R-squared Values** |  |
| Marginal R-squared (R²ₘ) | 0.00564 |
| Bootstrapped 2.5 % CI | 0.00200 |
| Bootstrapped 97.5 % CI | 0.01115 |
| Conditional R-squared (R²꜀) | 0.52780 |
| Bootstrapped 2.5 % CI | 0.49837 |
| Bootstrapped 97.5 % CI | 0.56131 |
| **Correlation of Fixed Effects** |  |
| Correlation (Intercept, Sigma_ampl) | 0.002 |

##### Figure 4: Locus coeruleus suppression

###### 4e: NE AUC

The analysis investigated the effect of condition (Arch vs. YFP (baseline)) on NE AUC using a linear mixed-effects model. The model included a random intercept for SubjectID to account for individual differences. The fixed effects estimates indicated that the condition (Arch) had a statistically significant negative impact on NE AUC.

Given slight issues with normality, bootstrapping was performed to provide robust CIs for the difference in estimates between conditions. These estimates did not overlap with zero, further supporting the primary finding of a significant effect.

| **Statistic** | **Value** |
| --- | --- |
| **Fixed Effects Estimates** |  |
| Intercept Estimate | -5.54 |
| Intercept SE | 7.26 |
| Intercept df | 6.26 |
| Intercept t-value | -0.76 |
| Intercept p-value | 0.47 |
| Condition (Arch) Estimate | -29.39 |
| Condition (Arch) SE | 10.09 |
| Condition (Arch) df | 5.84 |
| Condition (Arch) t-value | -2.91 |
| Condition (Arch) p-value | 0.03* |
| **Bootstrapped Difference in Estimates** |  |
| Bootstrapped 2.5% CI | -39.85 |
| Bootstrapped 97.5% CI | -18.18 |
| **Random Effects** |  |
| SubjectID (Intercept) Variance | 140.90 |
| SubjectID (Intercept) SD | 11.87 |
| Residual Variance | 655.10 |
| Residual SD | 25.60 |

###### 4e: RR AUC

The analysis investigated the effect of condition (Arch vs. YFP (baseline)) on RR AUC using a linear mixed-effects model. The model included a random intercept for SubjectID to account for individual differences. The fixed effects estimates indicated that the condition (Arch) did not have a statistically significant impact on RR AUC.

Given slight issues with normality and heteroscedasticity, bootstrapping was performed to provide robust CIs for the difference in estimates between conditions. These estimates did not overlap with zero, suggesting that there may be an effect of condition on RR AUC.

| **Statistic** | **Value** |
| --- | --- |
| **Fixed Effects Estimates** |  |
| Intercept Estimate | 27.35 |
| Intercept SE | 52.14 |
| Intercept df | 7.19 |
| Intercept t-value | 0.53 |
| Intercept p-value | 0.62 |
| Condition (Arch) Estimate | 95.67 |
| Condition (Arch) SE | 72.50 |
| Condition (Arch) df | 6.73 |
| Condition (Arch) t-value | 1.32 |
| Condition (Arch) p-value | 0.23 |
| **Bootstrapped Difference in Estimates** |  |
| Bootstrapped 2.5% CI | 25.49 |
| Bootstrapped 97.5% CI | 167.59 |
| **Random Effects** |  |
| SubjectID (Intercept) Variance | 7324.00 |
| SubjectID (Intercept) SD | 85.58 |
| Residual Variance | 33217.00 |
| Residual SD | 182.25 |

###### 4e: NE (AUC) & RR (AUC) correlation

The analysis investigated the effect of NE AUC on RR AUC using a linear mixed-effects model. The model included a random intercept for SubjectID to account for individual differences. The fixed effects estimates indicated that NE AUC had a statistically significant negative impact on RR AUC.

Given issues with normality and heteroscedasticity, bootstrapping was performed to provide robust CIs. These bootstrapped estimates of the marginal and conditional R squared overlapped with the initial estimate, further supporting the primary finding of a significant negative effect of NE AUC on RR AUC.

YFP

| **Statistic** | **Value** |
| --- | --- |
| **Fixed Effects Estimates** |  |
| Intercept Estimate | 17.828 |
| Intercept SE | 27.803 |
| Intercept df | 36.0 |
| Intercept t-value | 0.641 |
| Intercept p-value | 0.525 |
| NE Estimate | −1.958 |
| NE SE | 0.980 |
| NE df | 36.0 |
| NE t-value | −1.997 |
| NE p-value | 0.053 |
| **Random Effects** |  |
| SubjectID (Intercept) Variance | 0 |
| SubjectID (Intercept) SD | 0 |
| Residual Variance | 28 429 |
| Residual SD | 168.61 |
| **R-squared Values** |  |
| Marginal R-squared (R²ₘ) | 0.09732 |
| Bootstrapped 2.5 % CI | 0.00068 |
| Bootstrapped 97.5 % CI | 0.44597 |
| Conditional R-squared (R²꜀) | 0.09732 |
| Bootstrapped 2.5 % CI | 0.00199 |
| Bootstrapped 97.5 % CI | 0.49608 |
| **Correlation of Fixed Effects** |  |
| Correlation (Intercept, NE) | 0.179 |

| Arch  **Statistic** | **Value** |
| --- | --- |
| **Fixed Effects Estimates** |  |
| Intercept Estimate | 23.392 |
| Intercept SE | 64.710 |
| Intercept df | 5.807 |
| Intercept t-value | 0.361 |
| Intercept p-value | 0.7305 |
| NE Estimate | –2.851 |
| NE SE | 1.008 |
| NE df | 45.736 |
| NE t-value | –2.827 |
| NE p-value | 0.00695 ** |
| **Random Effects** |  |
| SubjectID (Intercept) Variance | 9 224 |
| SubjectID (Intercept) SD | 96.04 |
| Residual Variance | 30 020 |
| Residual SD | 173.26 |
| **R-squared Values** |  |
| Marginal R-squared (R²ₘ) | 0.13092 |
| Bootstrapped 2.5 % CI | 0.00336 |
| Bootstrapped 97.5 % CI | 0.41485 |
| Conditional R-squared (R²꜀) | 0.33519 |
| Bootstrapped 2.5 % CI | 0.19629 |
| Bootstrapped 97.5 % CI | 0.63459 |
| **Correlation of Fixed Effects** |  |
| Correlation (Intercept, NE) | 0.545 |

###### **Correlation comparison between Arch and YFP**

###### To test whether the coupling between the NE responce and RR differed between Arch- and YFP-expressing animals, we fit linear mixed-effects models with RR as the response and the NE response, group (Arch vs. YFP, YFP as reference), and their interaction as fixed effects. Each NREM bout was treated as an observation, and a by-animal random intercept accounted for the non-independence of bouts recorded from the same mouse. The Group× NE interaction term tested whether the NE–RR slope differed between groups. As a descriptive check, we also compared Pearson correlations using Fisher r-to-z transformation, noting that this approach does not account for repeated measures structure.

###### **Linear mixed-effects models of the NE–RR relationship**

| **NE→RR slope, YFP** | **NE→RR slope, Arch** | **Slope difference** | **R²ₘ** | **R²c** |
| --- | --- | --- | --- | --- |
| **−2.07 [−4.29, -0.151]** | **−3.07 [−5.10, −1.033]** | **0.994, p =0.51** | **0.18** | **0.28** |

###### **Fisher r-to-z transformation**

| **r_Arch** | **n_arch** | **r_YFP** | **n_YFP** | **Z** | **P_value** |
| --- | --- | --- | --- | --- | --- |
| **-0.468** | **48** | **-0.316** | **38** | **-0.802** | **0.4225** |

###### 4g: NE PSD (AUC)

The analysis examined the effects of condition (YFP and Arch) on NE PSD AUC. The results revealed a tendency for lower values for Arch compared to YFP, though this tendency was not statistically significant

Due to issues with normality and heteroscedasticity, bootstrapping based on Subject ID was used to provide robust CIs of the estimate-differences. The bootstrapped estimates suggested a robust difference between the two conditions.

| **Statistic** | **Value** |
| --- | --- |
| **Fixed Effects Estimates** |  |
| Intercept Estimate | 0.40462 |
| Intercept SE | 0.11467 |
| Intercept df | 6.03425 |
| Intercept t-value | 3.529 |
| Intercept p-value | 0.0123* |
| Condition (Arch) Estimate | -0.05404 |
| Condition (Arch) SE | 0.16429 |
| Condition (Arch) df | 6.34555 |
| Condition (Arch) t-value | -0.329 |
| Condition (Arch) p-value | 0.7528 |
| **Bootstrapped Difference in Estimates** |  |
| Bootstrapped 2.5% CI | -0.205904529 |
| Bootstrapped 97.5% CI | -0.006895428 |
| **Random Effects** |  |
| SubjectID (Intercept) Variance | 0.04577 |
| SubjectID (Intercept) SD | 0.2139 |
| Residual Variance | 25.94916 |
| Residual SD | 25.94916 |

###### 4g: RR PSD (AUC)

The analysis examined the effects of condition (YFP and Arch) on RR PSD AUC. The results revealed a tendency for lower values for Arch compared to YFP, though this tendency was not statistically significant

Due to issues with normality, bootstrapping based on Subject ID was used to provide robust CIs of the estimate-differences. The bootstrapped estimates suggested a robust difference between the two conditions.

| **Statistic** | **Value** |
| --- | --- |
| **Fixed Effects Estimates** |  |
| Intercept Estimate | 0.028871 |
| Intercept SE | 0.004390 |
| Intercept df | 5.759773 |
| Intercept t-value | 6.577 |
| Intercept p-value | 0.000699 *** |
| Condition (Arch) Estimate | -0.008149 |
| Condition (Arch) SE | 0.006366 |
| Condition (Arch) df | 6.333728 |
| Condition (Arch) t-value | -1.280 |
| Condition (Arch) p-value | 0.245392 |
| **Bootstrapped Difference in Estimates** |  |
| Bootstrapped 2.5% CI | -1.218052e-05 |
| Bootstrapped 97.5% CI | -5.005943e-06 |
| **Random Effects** |  |
| SubjectID (Intercept) Variance | 5.712e-05 |
| SubjectID (Intercept) SD | 0.007558 |
| Residual Variance | 7.613e-02 |
| Residual SD | 0.275925 |

###### 4g: NE frequency at peak power

The analysis examined the effects of condition (YFP and Arch) on frequency at peak PSD for NE. The results revealed no significant difference in frequency for Arch compared to YFP.

Due to issues with normality, bootstrapping based on Subject ID was used to provide robust CIs of the estimate-differences. The bootstrapped estimates did not overlap with 0, suggesting there might be an effect of the manipulation.

| **Statistic** | **Value** |
| --- | --- |
| **Fixed Effects Estimates** |  |
| Intercept Estimate | 0.018393 |
| Intercept SE | 0.001994 |
| Intercept df | 2.302453 |
| Intercept t-value | 9.226 |
| Intercept p-value | 0.00724 ** |
| Condition (Arch) Estimate | 0.002735 |
| Condition (Arch) SE | 0.002988 |
| Condition (Arch) df | 2.899209 |
| Condition (Arch) t-value | 0.915 |
| Condition (Arch) p-value | 0.42964 |
| **Bootstrapped Difference in Estimates** |  |
| Bootstrapped 2.5% CI | 0.0006023232 |
| Bootstrapped 97.5% CI | 0.0109565471 |
| **Random Effects** |  |
| SubjectID (Intercept) Variance | 5.539e-06 |
| SubjectID (Intercept) SD | 0.002353 |
| Residual Variance | 3.989e-02 |
| Residual SD | 0.199722 |

###### 4g: RR frequency at peak power

The analysis examined the effects of condition (YFP and Arch) on RR PSD AUC. The results revealed a tendency for lower values for Arch compared to YFP, though this tendency was not statistically significant.

Due to issues with normality, bootstrapping based on Subject ID was used to provide robust CIs of the estimate-differences. The bootstrapped estimates support the primary finding of no difference between the estimates.

| **Statistic** | **Value** |
| --- | --- |
| **Fixed Effects Estimates** |  |
| Intercept Estimate | 0.032385 |
| Intercept SE | 0.002781 |
| Intercept df | 106.999997 |
| Intercept t-value | 11.645 |
| Intercept p-value | <2e-16 *** |
| Condition (Arch) Estimate | -0.002238 |
| Condition (Arch) SE | 0.003890 |
| Condition (Arch) df | 106.999997 |
| Condition (Arch) t-value | -0.575 |
| Condition (Arch) p-value | 0.566 |
| **Bootstrapped Difference in Estimates** |  |
| Bootstrapped 2.5% CI | -0.009087315 |
| Bootstrapped 97.5% CI | 0.005413220 |
| **Random Effects** |  |
| SubjectID (Intercept) Variance | 2.341e-17 |
| SubjectID (Intercept) SD | 4.839e-09 |
| Residual Variance | 1.411e-01 |
| Residual SD | 3.757e-01 |

###### 4g: NE PSD peak power

The analysis examined the effects of condition (YFP and Arch) on NE peak power. The results revealed a tendency for lower values for Arch compared to YFP, though this tendency was not statistically significant.

Due to issues with normality, bootstrapping based on Subject ID was used to provide robust CIs of the estimate-differences. The bootstrapped estimates overlap with 0, suggesting support for the lack of difference.

| **Statistic** | **Value** |
| --- | --- |
| **Fixed Effects Estimates** |  |
| Intercept Estimate | 15.3334 |
| Intercept SE | 5.7206 |
| Intercept df | 5.3937 |
| Intercept t-value | 2.680 |
| Intercept p-value | 0.0406 * |
| Condition (Arch) Estimate | -0.7902 |
| Condition (Arch) SE | 8.2309 |
| Condition (Arch) df | 5.7669 |
| Condition (Arch) t-value | -0.096 |
| Condition (Arch) p-value | 0.9268 |
| **Bootstrapped Difference in Estimates** |  |
| Bootstrapped 2.5% CI | -9.854971 |
| Bootstrapped 97.5% CI | 2.573926 |
| **Random Effects** |  |
| SubjectID (Intercept) Variance | 108.1 |
| SubjectID (Intercept) SD | 10.4 |
| Residual Variance | 86661.3 |
| Residual SD | 294.4 |

###### 4g: RR PSD peak power

The analysis examined the effects of condition (YFP and Arch) on RR peak power. The results revealed a tendency for lower values for Arch compared to YFP, though this tendency was not statistically significant.

Due to issues with normality and heteroscedasticity, bootstrapping based on Subject ID was used to provide robust CIs of the estimate-differences. The bootstrapped estimates suggested a robust difference between the two conditions.

| **Statistic** | **Value** |
| --- | --- |
| **Fixed Effects Estimates** |  |
| Intercept Estimate | 6.261e-04 |
| Intercept SE | 1.091e-04 |
| Intercept df | 5.201 |
| Intercept t-value | 5.737 |
| Intercept p-value | 0.00198 ** |
| Condition (Arch) Estimate | -4.805e-05 |
| Condition (Arch) SE | -4.805e-05 |
| Condition (Arch) df | 5.492 |
| Condition (Arch) t-value | -0.307 |
| Condition (Arch) p-value | 0.77009 |
| **Bootstrapped Difference in Estimates** |  |
| Bootstrapped 2.5% CI | -2.661873e-04 |
| Bootstrapped 97.5% CI | -4.473375e-05 |
| **Random Effects** |  |
| SubjectID (Intercept) Variance | 4.105e-08 |
| SubjectID (Intercept) SD | 0.0002026 |
| Residual Variance | 2.504e-05 |
| Residual SD | 0.0050041 |

###### 4i: Theta (AUC)

The analysis investigated the effect of condition (Arch vs. YFP (baseline)) on theta AUC using a linear mixed-effects model. The model included a random intercept for SubjectID to account for individual differences. The fixed effects estimates indicated that the condition (Arch) had a statistically significant impact on Theta AUC, leading to an increase in AUC for Arch compared to YFP.

Given slight issues with normality and heteroscedasticity, bootstrapping was performed to provide robust CIs for the difference in estimates between conditions. These estimates did not overlap with zero, further supporting the primary finding of a significant effect of condition (Arch) on theta AUC.

| **Statistic** | **Value** |
| --- | --- |
| **Fixed Effects Estimates** |  |
| Intercept Estimate | -2.13 |
| Intercept SE | 1.21 |
| Intercept df | 5.96 |
| Intercept t-value | -1.75 |
| Intercept p-value | 0.13 |
| Condition (Arch) Estimate | 6.83 |
| Condition (Arch) SE | 1.68 |
| Condition (Arch) df | 5.50 |
| Condition (Arch) t-value | 4.06 |
| Condition (Arch) p-value | 0.01** |
| **Bootstrapped Difference in Estimates** |  |
| Bootstrapped 2.5% CI | 4.93 |
| Bootstrapped 97.5% CI | 8.92 |
| **Random Effects** |  |
| SubjectID (Intercept) Variance | 3.61 |
| SubjectID (Intercept) SD | 1.90 |
| Residual Variance | 21.42 |
| Residual SD | 4.63 |

###### 4i: Theta (AUC) & RR (AUC) correlation

The analysis investigated the effect of theta AUC on RR AUC using a linear mixed-effects model. The model included a random intercept for SubjectID to account for individual differences. The fixed effects estimates indicated that theta AUC had a statistically significant positive impact on RR AUC.

Given slight issues with normality, bootstrapping was performed to provide robust CIs. These bootstrapped estimates of the conditional and marginal R squared estimates overlapped with the initial estimate, further supporting the primary finding of a significant positive effect of theta AUC on RR AUC.

| **Statistic** | **Value** |
| --- | --- |
| **Fixed Effects Estimates** |  |
| Intercept Estimate | -1.29 |
| Intercept SE | 78.84 |
| Intercept df | 4.73 |
| Intercept t-value | -0.02 |
| Intercept p-value | 0.99 |
| Theta Estimate | 26.35 |
| Theta SE | 7.45 |
| Theta df | 44.87 |
| Theta t-value | 3.54 |
| Theta p-value | 0.001*** |
| **Random Effects** |  |
| SubjectID (Intercept) Variance | 17656.00 |
| SubjectID (Intercept) SD | 132.90 |
| Residual Variance | 26536.00 |
| Residual SD | 162.90 |
| **R-squared Values** |  |
| Marginal R-squared (R²m) | 0.16 |
| Bootstrapped 2.5% CI | 0.02 |
| Bootstrapped 97.5% CI | 0.33 |
| Conditional R-squared (R²c) | 0.49 |
| Bootstrapped 2.5% CI | 0.32 |
| Bootstrapped 97.5% CI | 0.71 |
| **Correlation of Fixed Effects** |  |
| Correlation (Intercept, Theta) | -0.45 |

###### 4i: Sigma (AUC)

The analysis investigated the effect of condition (Arch vs. YFP (baseline)) on sigma AUC using a linear mixed-effects model. The model included a random intercept for SubjectID to account for individual differences. The fixed effects estimates indicated that the condition (Arch) had a statistically significant impact on sigma AUC, leading to an increase in AUC for Arch compared to YFP.

Given slight issues with normality, bootstrapping was performed to provide robust CIs for the difference in estimates between conditions. These estimates did not overlap with zero, further supporting the primary finding of a significant effect of condition (Arch) on sigma AUC.

| **Statistic** | **Value** |
| --- | --- |
| **Fixed Effects Estimates** |  |
| Intercept Estimate | -1.39 |
| Intercept SE | 0.97 |
| Intercept df | 5.78 |
| Intercept t-value | -1.43 |
| Intercept p-value | 0.20 |
| Condition (Arch) Estimate | 6.95 |
| Condition (Arch) SE | 1.33 |
| Condition (Arch) df | 4.95 |
| Condition (Arch) t-value | 5.25 |
| Condition (Arch) p-value | 0.003** |
| **Bootstrapped Difference in Estimates** |  |
| Bootstrapped 2.5% CI | 4.74 |
| Bootstrapped 97.5% CI | 9.05 |
| **Random Effects** |  |
| SubjectID (Intercept) Variance | 1.08 |
| SubjectID (Intercept) SD | 1.04 |
| Residual Variance | 25.55 |
| Residual SD | 5.06 |

###### 4i: Sigma (AUC) & RR (AUC) correlation

The analysis investigated the effect of sigma AUC on RR AUC using a linear mixed-effects model. The model included a random intercept for SubjectID to account for individual differences. The fixed effects estimates indicated that sigma AUC had a statistically significant positive impact on RR AUC.

Given slight issues with normality, bootstrapping was performed to provide robust CIs. These bootstrapped estimates of the conditional and marginal R squared estimates overlapped with the initial estimate, further supporting the primary finding of a significant positive effect of sigma AUC on RR AUC.

| **Statistic** | **Value** |
| --- | --- |
| **Fixed Effects Estimates** |  |
| Intercept Estimate | 18.79 |
| Intercept SE | 67.41 |
| Intercept df | 4.52 |
| Intercept t-value | 0.28 |
| Intercept p-value | 0.79 |
| Sigma Estimate | 18.91 |
| Sigma SE | 5.10 |
| Sigma df | 43.87 |
| Sigma t-value | 3.71 |
| Sigma p-value | 0.001*** |
| **Random Effects** |  |
| SubjectID (Intercept) Variance | 12762.00 |
| SubjectID (Intercept) SD | 113.00 |
| Residual Variance | 26388.00 |
| Residual SD | 162.40 |
| **R-squared Values** |  |
| Marginal R-squared (R²m) | 0.17 |
| Bootstrapped 2.5% CI | 0.04 |
| Bootstrapped 97.5% CI | 0.35 |
| Conditional R-squared (R²c) | 0.44 |
| Bootstrapped 2.5% CI | 0.28 |
| Bootstrapped 97.5% CI | 0.68 |
| **Correlation of Fixed Effects** |  |
| Correlation (Intercept, Sigma) | -0.42 |

###### 4i: Beta (AUC)

The analysis investigated the effect of condition (Arch vs. YFP (baseline)) on beta AUC using a linear mixed-effects model. The model included a random intercept for SubjectID to account for individual differences. The fixed effects estimates indicated that the condition (Arch) had a statistically significant impact on beta AUC, leading to an increase in AUC for Arch compared to YFP.

Given slight issues with normality and heteroscedasticity, bootstrapping was performed to provide robust CIs for the difference in estimates between conditions. These estimates did not overlap with zero, further supporting the primary finding of a significant effect of condition (Arch) on beta AUC.

| **Statistic** | **Value** |
| --- | --- |
| **Fixed Effects Estimates** |  |
| Intercept Estimate | -1.08 |
| Intercept SE | 0.80 |
| Intercept df | 5.50 |
| Intercept t-value | -1.34 |
| Intercept p-value | 0.23 |
| Condition (Arch) Estimate | 4.85 |
| Condition (Arch) SE | 1.09 |
| Condition (Arch) df | 4.74 |
| Condition (Arch) t-value | 4.43 |
| Condition (Arch) p-value | 0.008** |
| **Bootstrapped Difference in Estimates** | |
| Bootstrapped 2.5% CI | 3.09 |
| Bootstrapped 97.5% CI | 6.65 |
| **Random Effects** |  |
| SubjectID (Intercept) Variance | 0.81 |
| SubjectID (Intercept) SD | 0.90 |
| Residual Variance | 16.64 |
| Residual SD | 4.08 |

###### 4i: Beta (AUC) & RR (AUC) correlation

The analysis investigated the effect of beta AUC on RR AUC using a linear mixed-effects model. The model included a random intercept for SubjectID to account for individual differences. The fixed effects estimates indicated that beta AUC had a positive, though not statistically significant, impact on RR AUC.

Given slight issues with normality and heteroscedasticity, bootstrapping was performed to provide robust CIs. These bootstrapped estimates of the conditional and marginal R squared estimates overlapped with the initial estimate, further supporting the primary finding of a significant positive effect of beta AUC on RR AUC.

| **Statistic** | **Value** |
| --- | --- |
| **Fixed Effects Estimates** |  |
| Intercept Estimate | 69.87 |
| Intercept SE | 71.42 |
| Intercept df | 4.41 |
| Intercept t-value | 0.98 |
| Intercept p-value | 0.38 |
| Beta Estimate | 14.08 |
| Beta SE | 7.69 |
| Beta df | 43.31 |
| Beta t-value | 1.83 |
| Beta p-value | 0.07 |
| **Random Effects** |  |
| SubjectID (Intercept) Variance | 14295.00 |
| SubjectID (Intercept) SD | 119.60 |
| Residual Variance | 32083.00 |
| Residual SD | 179.10 |
| **R-squared Values** |  |
| Marginal R-squared (R²m) | 0.05 |
| Bootstrapped 2.5% CI | 0.00 |
| Bootstrapped 97.5% CI | 0.22 |
| Conditional R-squared (R²c) | 0.34 |
| Bootstrapped 2.5% CI | 0.17 |
| Bootstrapped 97.5% CI | 0.60 |
| **Correlation of Fixed Effects** |  |
| Correlation (Intercept, Beta) | -0.41 |

###### 4j: Novel-to-Familiar Ratio & RR (AUC) correlation

The analysis investigated the effect of the NFR on RR AUC using a linear model. The fixed effects estimates indicated that the NFR had a statistically significant positive impact on RR AUC.

Given issues with normality due to the model being based on only 8 data points, bootstrapping was performed to provide robust CIs. These bootstrapped estimates of the marginal and conditional R squared overlapped with the initial estimate, further supporting the primary finding of a significant positive effect of the NFR on RR AUC.

| **Statistic** | **Value** |
| --- | --- |
| **Fixed Effects Estimates** |  |
| Intercept Estimate | -60.33 |
| Intercept SE | 46.19 |
| Intercept t-value | -1.31 |
| Intercept p-value | 0.24 |
| Novel Familiar Ratio Estimate | 94.64 |
| Novel Familiar Ratio SE | 27.98 |
| Novel Familiar Ratio t-value | 3.38 |
| Novel Familiar Ratio p-value | 0.01* |
| **Model Summary** |  |
| Residual Standard Error | 65.27 |
| Degrees of Freedom (Residuals) | 6 |
| Multiple R-squared | 0.66 |
| Adjusted R-squared | 0.60 |
| F-statistic | 11.44 |
| F-statistic Degrees of Freedom (df1, df2) | 1, 6 |
| F-statistic p-value | 0.01 |
| **R-squared Values** |  |
| Multiple R-squared | 0.66 |
| Bootstrapped 2.5% CI | 0.00 |
| Bootstrapped 97.5% CI | 0.97 |
| Adjusted R-squared | 0.60 |
| Bootstrapped 2.5% CI | -0.16 |
| Bootstrapped 97.5% CI | 0.96 |

###### 4j: RR (AUC) & NE (AUC) correlation

The analysis investigated the effect of the NE AUC on RR AUC using a linear model. The fixed effects estimates indicated that the NE AUC had a significant negative impact on RR AUC.

Given issues with normality due to the model being based on only 8 data points, bootstrapping was performed to provide robust CIs. These bootstrapped estimates of the conditional and marginal R squared estimates overlapped with the initial estimate, further supporting the primary finding.

| **Statistic** | **Value** |
| --- | --- |
| **Fixed Effects Estimates** |  |
| Intercept Estimate | -0.9322 |
| Intercept SE | 38.0134 |
| Intercept t-value | -0.025 |
| Intercept p-value | 0.9812 |
| NE Estimate | -3.7321 |
| NE SE | 1.3557 |
| NE t-value | -2.753 |
| NE p-value | 0.0332 * |
| **Model Summary** |  |
| Residual Standard Error | 73.97 |
| Degrees of Freedom (Residuals) | 6 |
| Multiple R-squared | 0.5581 |
| Adjusted R-squared | 0.4845 |
| F-statistic | 7.579 |
| F-statistic Degrees of Freedom (df1, df2) | 1, 6 |
| F-statistic p-value | 0.03316 |
| **R-squared Values** |  |
| Multiple R-squared | 0.56 |
| Bootstrapped 2.5% CI | 0.03 |
| Bootstrapped 97.5% CI | 0.92 |
| Adjusted R-squared | 0.48 |
| Bootstrapped 2.5% CI | -0.14 |
| Bootstrapped 97.5% CI | 0.90 |

##### Figure 5: Heart rate bursts

###### 5d: NE (ampl.) & RR (ampl.)

The correlation between the RR and NE amplitude in connection with a HRB showed a negative relationship. Due to slight issues with heteroscedasticity, the marginal and conditional R squared estimates were bootstrapped, which supported the primary findings.

| **Statistic** | **Value** |
| --- | --- |
| **Fixed Effects Estimates** |  |
| Intercept Estimate | -0.013 |
| Intercept SE | 0.00 |
| Intercept df | 5.80 |
| Intercept t-value | -8.21 |
| Intercept p-value | 0.0002 |
| NE Estimate | -0.004 |
| NE SE | 0.000 |
| NE df | 1652 |
| NE t-value | -21.07 |
| NE p-value | <0.0001*** |
| **Random Effects** |  |
| SubjectID (Intercept) Variance | 1.73e-05 |
| SubjectID (Intercept) SD | 0.004 |
| Residual Variance | 7.17e-05 |
| Residual SD | 0.008 |
| **R-squared Values** |  |
| Marginal R-squared (R²m) | 0.21 |
| Bootstrapped 2.5% CI | 0.18 |
| Bootstrapped 97.5% CI | 0.25 |
| Conditional R-squared (R²c) | 0.37 |
| Bootstrapped 2.5% CI | 0.32 |
| Bootstrapped 97.5% CI | 0.42 |
| **Correlation of Fixed Effects** |  |
| Correlation (Intercept, NE) | -0.15 |

###### 5f: Sigma (ampl.) & RR (ampl.)

The correlation between the RR and Sigma amplitude in connection with a HRB showed a positive relationship. Due to slight issues with heteroscedasticity, the marginal and conditional R squared estimates were bootstrapped, which supported the primary findings.

| **Statistic** | **Value** |
| --- | --- |
| **Fixed Effects Estimates** |  |
| Intercept Estimate | -0.012 |
| Intercept SE | 0.0015 |
| Intercept df | 12.03 |
| Intercept t-value | -8.376 |
| Intercept p-value | 2.3e-06*** |
| Sigma Estimate | 0.005 |
| Sigma SE | 0.0006 |
| Sigma df | 1543 |
| Sigma t-value | 8.649 |
| Sigma p-value | <0.0001*** |
| **Random Effects** |  |
| SubjectID (Intercept) Variance | 9.556e-06 |
| SubjectID (Intercept) SD | 0.003 |
| Residual Variance | 8.716e-05 |
| Residual SD | 0.009 |
| **R-squared Values** |  |
| Marginal R-squared (R²m) | 0.06 |
| Bootstrapped 2.5% CI | 0.03 |
| Bootstrapped 97.5% CI | 0.08 |
| Conditional R-squared (R²c) | 0.15 |
| Bootstrapped 2.5% CI | 0.12 |
| Bootstrapped 97.5% CI | 0.20 |
| **Correlation of Fixed Effects** |  |
| Correlation (Intercept, Sigma) | 0.530 |

###### 5g: Sigma (ampl.) & NE (ampl.)

The correlation between the Sigma and NE amplitude in connection with a HRB showed a negative relationship. Due to issues with heteroscedasticity, the marginal and conditional R squared estimates were bootstrapped, which supported the primary findings.

| **Fixed Effects Estimates** |  |
| --- | --- |
| Intercept Estimate | -0.016 |
| Intercept SE | 0.25 |
| Intercept df | 6.85 |
| Intercept t-value | -0.065 |
| Intercept p-value | 0.95 |
| Sigma Estimate | -0.93 |
| Sigma SE | 0.059 |
| Sigma df | 1648.29 |
| Sigma t-value | -15.9 |
| Sigma p-value | <2e-16 *** |
| **Random Effects** |  |
| SubjectID (Intercept) Variance | 0.38 |
| SubjectID (Intercept) SD | 0.62 |
| Residual Variance | 0.78 |
| Residual SD | 0.88 |
| **R-squared Values** |  |
| Marginal R-squared (R²m) | 0.13 |
| Bootstrapped 2.5% CI | 0.10 |
| Bootstrapped 97.5% CI | 0.16 |
| Conditional R-squared (R²c) | 0.42 |
| Bootstrapped 2.5% CI | 0.36 |
| Bootstrapped 97.5% CI | 0.50 |
| **Correlation of Fixed Effects** |  |
| Correlation (Intercept, Sigma) | 0.289 |

###### 5k: Sigma & RR (ampl.)

The correlation between the post-HRB RR amplitude and Sigma power in connection with a HRB showed a positive relationship. Due to slight issues with heteroscedastic, the marginal and conditional R squared estimates were bootstrapped, which supported the primary findings.

| **Statistic** | **Value** |
| --- | --- |
| **Fixed Effects Estimates** |  |
| Intercept Estimate | 0.2472 |
| Intercept SE | 0.02576 |
| Intercept df | 29.88 |
| Intercept t-value | 9.594 |
| Intercept p-value | 1.24 × 10⁻¹⁰ *** |
| sigma Estimate | 0.02558 |
| sigma SE | 0.002343 |
| sigma df | 4739.00 |
| sigma t-value | 10.918 |
| sigma p-value | < 2 × 10⁻¹⁶ *** |
| **Random Effects** |  |
| SubjectID (Intercept) Variance | 0.01755 |
| SubjectID (Intercept) SD | 0.1325 |
| Residual Variance | 0.01681 |
| Residual SD | 0.1297 |
| **R-squared Values** |  |
| Marginal R-squared (R²ₘ) | 0.01563 |
| Bootstrapped 2.5 % CI | 0.01041 |
| Bootstrapped 97.5 % CI | 0.02187 |
| Conditional R-squared (R²꜀) | 0.51843 |
| Bootstrapped 2.5 % CI | 0.50030 |
| Bootstrapped 97.5 % CI | 0.54075 |
| **Correlation of Fixed Effects** |  |
| Correlation (Intercept, sigma) | –0.220 |

###### 5m: Sigma (bl. Cor.) & memory improvement

The correlation between memory improvement and baseline corrected sigma power in connection with a HRB showed a slight positive relationship when factoring in baseline memory scores (BMS) and the non-ACE baseline. Due to slight issues with heteroscedastic, the marginal and conditional R squared estimates were bootstrapped, which supported the primary findings.

| **Statistic** | **Value** |
| --- | --- |
| **Fixed Effects Estimates** |  |
| Intercept Estimate | -0.071 |
| Intercept SE | 0.219 |
| Intercept t-value | -0.33 |
| Intercept p-value | 0.747 |
| Subbase Estimate | 1.098 |
| Subbase SE | 0.539 |
| Subbase t-value | 2.04 |
| Subbase p-value | 0.053 . |
| nonace_bl Estimate | -0.131 |
| nonace_bl SE | 0.077 |
| nonace_bl t-value | -1.71 |
| nonace_bl p-value | 0.101 |
| BMS Estimate | -0.183 |
| BMS SE | 0.199 |
| BMS t-value | -0.92 |
| BMS p-value | 0.367 |
| **Residuals** |  |
| Residual SD | 0.173 |
| Residual df | 24 |
| **R-squared Values** |  |
| Multiple R-squared | 0.186 |
| 2.5% CI (Mul. R²) | 0.073 |
| 97.5% CI (Mul. R²) | 0.515 |
| Adjusted R-squared | 0.084 |
| 2.5% CI (Adj. R²) | -0.055 |
| 97.5% CI (Adj. R²) | 0.451 |
| F-statistic | 1.83 |
| F-statistic df | 3, 24 |
| F-statistic p-value | 0.169 |

##### Supplementary figure 1:

S1g(1): NE amplitude & NE AUC correlation

The correlation between NE amplitude and AUC showed a strong positive relationship. Due to slight issues with normality of residuals, the marginal and conditional R squared estimates were bootstrapped, which supported the primary findings.

| **Statistic** | **Value** |
| --- | --- |
| **Fixed Effects Estimates** |  |
| Intercept Estimate | -25.4796 |
| Intercept SE | 3.297 |
| Intercept df | 7.21 |
| Intercept t-value | -7.73 |
| Intercept p-value | <0.0001*** |
| NE amp Estimate | 17.37 |
| NE amp SE | 0.4234 |
| NE amp df | 554.8077 |
| NE amp t-value | 41.032 |
| NE amp p-value | <0.0001*** |
| **Random Effects** |  |
| SubjectID (Intercept) Variance | 68.64 |
| SubjectID (Intercept) SD | 8.285 |
| Residual Variance | 74.54 |
| Residual SD | 8.634 |
| **R-squared Values** |  |
| Marginal R-squared (R²m) | 0.67 |
| Bootstrapped 2.5% CI | 0.61 |
| Bootstrapped 97.5% CI | 0.72 |
| Conditional R-squared (R²c) | 0.83 |
| Bootstrapped 2.5% CI | 0.80 |
| Bootstrapped 97.5% CI | 0.86 |
| **Correlation of Fixed Effects** |  |
| Correlation (Intercept, NE amp) | -0.29 |

S1g(2): RR amplitude & RR AUC correlation

The correlation between RR amplitude and AUC showed a positive relationship. Due to slight issues with normality of residuals, the marginal and conditional R squared estimates were bootstrapped, which supported the primary findings.

| **Statistic** | **Value** |
| --- | --- |
| **Fixed Effects Estimates** |  |
| Intercept Estimate | 64.5570 |
| Intercept SE | 20.7764 |
| Intercept df | 8.4991 |
| Intercept t-value | 3.107 |
| Intercept p-value | 0.0135* |
| RR amp Estimate | 8.2631 |
| RR amp SE | 0.4359 |
| RR amp df | 557.6070 |
| RR amp t-value | 18.956 |
| RR amp p-value | <0.0001*** |
| **Random Effects** |  |
| SubjectID (Intercept) Variance | 2313 |
| SubjectID (Intercept) SD | 48.09 |
| Residual Variance | 15795 |
| Residual SD | 125.68 |
| **R-squared Values** |  |
| Marginal R-squared (R²m) | 0.38 |
| Bootstrapped 2.5% CI | 0.32 |
| Bootstrapped 97.5% CI | 0.44 |
| Conditional R-squared (R²c) | 0.46 |
| Bootstrapped 2.5% CI | 0.40 |
| Bootstrapped 97.5% CI | 0.53 |
| **Correlation of Fixed Effects** |  |
| Correlation (Intercept, RR amp) | 0.40 |

S1g(3): RR amplitude & NE amplitude correlation

The correlation between NE amplitude and RR amplitude showed a negative relationship. Due to slight issues with normality of residuals, the marginal and conditional R squared estimates were bootstrapped, which supported the primary findings.

| **Statistic** | **Value** |
| --- | --- |
| **Fixed Effects Estimates** |  |
| Intercept Estimate | -8.569 |
| Intercept SE | 2.453 |
| Intercept df | 11.001 |
| Intercept t-value | -3.494 |
| Intercept p-value | 0.00502** |
| NE Estimate | -4.698 |
| NE SE | 0.561 |
| NE df | 557.946 |
| NE t-value | -8.375 |
| NE p-value | <0.0001*** |
| **Random Effects** |  |
| SubjectID (Intercept) Variance | 29.02 |
| SubjectID (Intercept) SD | 5.387 |
| Residual Variance | 132.00 |
| Residual SD | 11.489 |
| **R-squared Values** |  |
| Marginal R-squared (R²m) | 0.12 |
| Bootstrapped 2.5% CI | 0.07 |
| Bootstrapped 97.5% CI | 0.17 |
| Conditional R-squared (R²c) | 0.28 |
| Bootstrapped 2.5% CI | 0.21 |
| Bootstrapped 97.5% CI | 0.37 |
| **Correlation of Fixed Effects** |  |
| Correlation (Intercept, NE) | -0.517 |

##### Supplementary figure 2: EEG for Figure 2

###### S2c: Delta Power (AUC)

The analysis investigated the effects of laser threshold levels on delta Power (AUC), with the condition (ChR2 and YFP) included as an additional predictor. The findings revealed that laser threshold levels did not significantly affect delta across most comparisons. However, significant differences were found between the ChR2 and YFP conditions at thresholds -15, -10, and -5, with ChR2 consistently showing higher delta power (AUC) than YFP at these levels.

Given slight issues with normality, bootstrapping was performed to provide robust CIs for the estimates in the pairwise comparisons. These estimates supported the primary findings of ChR2 being larger than YFP at thresholds -15, -10, and -5.

**Fixed Effects Estimates**

| **Predictor** | **Estimate** | **Std. Error** | **df** | **t-value** | **p-value** |
| --- | --- | --- | --- | --- | --- |
| **(Intercept)** | 0.8180 | 0.5129 | 70.64 | 1.595 | 0.11518 |
| **Threshold - 15 (Reference)** | - | - | - | - | - |
| Threshold - 10 | 0.4326 | 0.5130 | 3726 | 0.843 | 0.39906 |
| Threshold - 5 | 0.0056 | 0.4660 | 3726 | 0.012 | 0.99037 |
| Threshold 0 | -0.3057 | 0.4470 | 3727 | -0.684 | 0.49414 |
| Threshold 5 | -0.4871 | 0.4542 | 3724 | -1.072 | 0.28362 |
| **Condition (YFP)** | -3.7820 | 1.3900 | 545.3 | -2.721 | 0.00672 ** |
| Threshold - 10 * Condition (YFP) | 1.1330 | 1.4560 | 3723 | 0.778 | 0.43669 |
| Threshold - 5 * Condition (YFP) | 1.6620 | 1.3480 | 3722 | 1.233 | 0.21767 |
| Threshold 0 * Condition (YFP) | 3.2000 | 1.3370 | 3725 | 2.394 | 0.01673 * |
| Threshold 5 * Condition (YFP) | 3.5630 | 1.3450 | 3726 | 2.649 | 0.00810 ** |

**Pairwise Comparisons Within ChR2 and YFP Conditions**

**Within ChR2 Condition**

| **Contrast** | **Estimate** | **SE** | **df** | **t-ratio** | **p-value** | **Lower Bound (2.5%)** | **Upper Bound (97.5%)** |
| --- | --- | --- | --- | --- | --- | --- | --- |
| -15 vs -10 | -0.4326 | 0.513 | 3726 | -0.843 | 0.9171 | -1.3848 | 0.5583 |
| -15 vs -5 | -0.0056 | 0.466 | 3726 | -0.012 | 1.0000 | -0.7410 | 0.8357 |
| -15 vs 0 | 0.3057 | 0.447 | 3727 | 0.683 | 0.9601 | -0.3865 | 1.0734 |
| -15 vs 5 | 0.4871 | 0.455 | 3724 | 1.071 | 0.8213 | -0.3330 | 1.3304 |
| -10 vs -5 | 0.4269 | 0.349 | 3724 | 1.225 | 0.7369 | -0.2718 | 1.0947 |
| -10 vs 0 | 0.7382 | 0.320 | 3726 | 2.305 | 0.1434 | 0.0475 | 1.3979 |
| -10 vs 5 | 0.9196 | 0.328 | 3727 | 2.800 | 0.0411 * | 0.2272 | 1.6111 |
| -5 vs 0 | 0.3113 | 0.242 | 3613 | 1.288 | 0.6989 | -0.1109 | 0.7376 |
| -5 vs 5 | 0.4927 | 0.250 | 3652 | 1.972 | 0.2801 | 0.0079 | 0.9751 |
| 0 vs 5 | 0.1814 | 0.202 | 3727 | 0.897 | 0.8981 | -0.2126 | 0.6037 |

**Within YFP Condition**

| **Contrast** | **Estimate** | **SE** | **df** | **t-ratio** | **p-value** | **Lower Bound (2.5%)** | **Upper Bound (97.5%)** |
| --- | --- | --- | --- | --- | --- | --- | --- |
| -15 vs -10 | -1.5655 | 1.363 | 3722 | -1.148 | 0.7807 | -4.2057 | 1.0688 |
| -15 vs -5 | -1.6679 | 1.265 | 3719 | -1.318 | 0.6798 | -3.8285 | 0.4928 |
| -15 vs 0 | -2.8944 | 1.260 | 3724 | -2.297 | 0.1460 | -5.1592 | -1.1637 |
| -15 vs 5 | -3.0755 | 1.266 | 3724 | -2.429 | 0.1077 | -5.9632 | -0.6701 |
| -10 vs -5 | -0.1024 | 0.609 | 3727 | -0.168 | 0.9998 | -2.0375 | 0.8148 |
| -10 vs 0 | -1.3289 | 0.594 | 3722 | -2.237 | 0.1662 | -3.1622 | -0.6701 |
| -10 vs 5 | -1.5100 | 0.607 | 3716 | -2.489 | 0.0932 | -3.8285 | -0.8148 |
| -5 vs 0 | -1.2265 | 0.313 | 3681 | -3.922 | 0.0008 *** | -3.0016 | 0.4124 |
| -5 vs 5 | -1.4076 | 0.334 | 3658 | -4.208 | 0.0003 *** | -4.4575 | 0.8216 |
| 0 vs 5 | -0.1811 | 0.280 | 3722 | -0.646 | 0.9674 | -1.7954 | 0.3923 |

**Between Conditions (ChR2 vs YFP)**

| **Threshold** | **Estimate** | **SE** | **df** | **t-value** | **p-value** | **Lower Bound (2.5%)** | **Upper Bound (97.5%)** |
| --- | --- | --- | --- | --- | --- | --- | --- |
| Threshold -15 | 3.7820 | 1.390 | 536.5 | 2.720 | 0.0067 ** | 1.6619 | 5.8176 |
| Threshold -10 | 2.6490 | 0.774 | 58.0 | 3.421 | 0.0011 ** | 0.8216 | 4.4575 |
| Threshold -5 | 2.1200 | 0.545 | 14.4 | 3.891 | 0.0016 ** | 1.5914 | 2.7189 |
| Threshold 0 | 0.5820 | 0.504 | 10.6 | 1.154 | 0.2739 | 0.1259 | 1.0253 |
| Threshold 5 | 0.2190 | 0.521 | 12.0 | 0.422 | 0.6808 | -0.3407 | 0.7691 |

###### S2c: Theta Power (AUC)

The analysis investigated the effects of laser threshold levels on theta power (AUC), with the condition (ChR2 and YFP) included as an additional predictor. The findings revealed that laser threshold levels significantly affected theta power (AUC) in the ChR2 condition, with higher thresholds (0 and 5) consistently showing higher theta power (AUC) than lower thresholds (-15 and -10). In the YFP condition, the effects of laser threshold levels were generally not significant.

Significant differences were found between the ChR2 and YFP conditions at thresholds -15, -10, and -5, with ChR2 consistently showing higher theta power (AUC) than YFP at these levels. Given slight issues with heteroscedasticity, bootstrapping was performed to provide robust CIs for the estimates in the pairwise comparisons. These estimates supported the primary findings of ChR2 having larger theta power (AUC) than YFP at thresholds -15, -10, and -5.

**Fixed Effects of Laser Thresholds and Condition on Theta Power (AUC)**

| **Predictor** | **Estimate** | **Std. Error** | **df** | **t-value** | **p-value** |
| --- | --- | --- | --- | --- | --- |
| **(Intercept)** | 3.9322 | 0.4973 | 46.65 | 7.907 | 3.69e-10 *** |
| **Threshold - 15 (Reference)** | - | - | - | - | - |
| Threshold - 10 | 0.5946 | 0.4675 | 3725 | 1.272 | 0.2035 |
| Threshold - 5 | 2.6650 | 0.4248 | 3727 | -6.273 | 3.95e-10 *** |
| Threshold 0 | 3.1611 | 0.4075 | 3727 | -7.757 | 1.11e-14 *** |
| Threshold 5 | 3.3155 | 0.4141 | 3727 | -8.006 | 1.56e-15 *** |
| **Condition (YFP)** | -6.1599 | 1.2949 | 323.4 | -4.757 | 2.96e-06 *** |
| Threshold - 10 * Condition (YFP) | 2.7366 | 1.3274 | 3722 | 2.062 | 0.0393 * |
| Threshold - 5 * Condition (YFP) | 4.7442 | 1.2287 | 3721 | 3.861 | 0.0001 *** |
| Threshold 0 * Condition (YFP) | 5.1323 | 1.2185 | 3723 | 4.212 | 2.59e-05 *** |
| Threshold 5 * Condition (YFP) | 5.5208 | 1.2257 | 3724 | 4.504 | 6.86e-06 *** |

**Pairwise Comparisons with Bootstrapped Confidence Intervals**

| **Contrast** | **Estimate** | **SE** | **df** | **t-ratio** | **p-value** | **Lower Bound (2.5%)** | **Upper Bound (97.5%)** |
| --- | --- | --- | --- | --- | --- | --- | --- |
| **Within ChR2 Condition** | | | |  |  |  |  |
| -15 vs -10 | -0.5946 | 0.468 | 3725 | -1.272 | 0.7088 | -0.3492 | 1.5405 |
| -15 vs -5 | -2.6650 | 0.425 | 3727 | -6.270 | <.0001 | -4.0965 | -0.5586 |
| -15 vs 0 | -3.1611 | 0.408 | 3727 | -7.753 | <.0001 | -4.0965 | -1.2443 |
| -15 vs 5 | -3.3155 | 0.414 | 3727 | -8.002 | <.0001 | -4.0778 | -0.3767 |
| -10 vs -5 | -2.0704 | 0.318 | 3727 | -6.516 | <.0001 | -3.8936 | -0.2822 |
| -10 vs 0 | -2.5665 | 0.292 | 3725 | -8.792 | <.0001 | -4.0965 | -0.5586 |
| -10 vs 5 | -2.7209 | 0.299 | 3727 | -9.089 | <.0001 | -4.0778 | -0.3767 |
| -5 vs 0 | -0.4961 | 0.220 | 3677 | -2.250 | 0.1617 | -1.2443 | -0.5586 |
| -5 vs 5 | -0.6506 | 0.228 | 3697 | -2.855 | 0.0351 | -4.0965 | -0.5586 |
| 0 vs 5 | -0.1545 | 0.184 | 3726 | -0.838 | 0.9188 | -4.0778 | -0.3767 |
| **Within YFP Condition** | |  |  |  |  |  |  |
| -15 vs -10 | -2.1420 | 1.242 | 3722 | -1.724 | 0.4192 | -4.3741 | 1.1079 |
| -15 vs -5 | -2.0793 | 1.153 | 3719 | -1.803 | 0.3716 | -4.3741 | 1.1079 |
| -15 vs 0 | -1.9712 | 1.149 | 3723 | -1.716 | 0.4239 | -4.3741 | 1.1079 |
| -15 vs 5 | -2.2053 | 1.154 | 3723 | -1.911 | 0.3114 | -4.3741 | 1.1079 |
| -10 vs -5 | 0.0627 | 0.555 | 3726 | 0.113 | 1.0000 | -4.3741 | 1.1079 |
| -10 vs 0 | 0.1708 | 0.542 | 3726 | 0.315 | 0.9979 | -4.3741 | 1.1079 |
| -10 vs 5 | -0.0633 | 0.553 | 3724 | -0.114 | 1.0000 | -4.3741 | 1.1079 |
| -5 vs 0 | 0.1080 | 0.285 | 3711 | 0.379 | 0.9956 | -4.3741 | 1.1079 |
| -5 vs 5 | -0.1260 | 0.305 | 3700 | -0.413 | 0.9939 | -4.3741 | 1.1079 |
| 0 vs 5 | -0.2340 | 0.255 | 3721 | -0.916 | 0.8907 | -4.3741 | 1.1079 |
| **Between ChR2 and YFP** | | | |  |  |  |  |
| Threshold -15 | 6.1600 | 1.295 | 321.9 | 4.757 | <.0001 | 4.3921 | 8.2065 |
| Threshold -10 | 3.4230 | 0.755 | 39.5 | 4.535 | 0.0001 | 2.1086 | 4.7342 |
| Threshold -5 | 1.4160 | 0.564 | 12.4 | 2.509 | 0.0269 | 0.8629 | 1.9665 |
| Threshold 0 | 1.0280 | 0.532 | 9.81 | 1.932 | 0.0827 | 0.6285 | 1.4308 |
| Threshold 5 | 0.6390 | 0.545 | 10.8 | 1.173 | 0.2660 | 0.1349 | 1.1694 |

###### S2c: Sigma Power (AUC)

The analysis investigated the effects of laser threshold levels on sigma power (AUC), with the condition (ChR2 and YFP) included as an additional predictor. The findings revealed significant effects of laser threshold levels on sigma power (AUC) in the ChR2 condition, with higher thresholds (0 and 5) consistently showing higher sigma power (AUC) than lower thresholds (-15 and -10). In contrast, the YFP condition showed minimal differences across laser threshold levels, with most comparisons being non-significant.

Significant differences were found between the ChR2 and YFP conditions at all laser thresholds, with ChR2 consistently showing higher sigma power (AUC) than YFP. Due to issues with heteroscedasticity, bootstrapping was performed to provide robust CIs for the estimates in the pairwise comparisons, reinforcing the primary findings that ChR2 had consistently larger sigma power (AUC) compared to YFP.

**Fixed Effects of Laser Thresholds and Condition on Sigma Power (AUC)**

| **Predictor** | **Estimate** | **Std. Error** | **df** | **t-value** | **p-value** |
| --- | --- | --- | --- | --- | --- |
| **(Intercept)** | 4.8694 | 0.5023 | 125.25 | 9.695 | < 2e-16 *** |
| **Threshold - 15 (Reference)** | - | - | - | - | - |
| Threshold - 10 | 0.9859 | 0.5348 | 3727 | -1.843 | 0.0653 . |
| Threshold - 5 | 3.3084 | 0.4857 | 3716 | -6.811 | 1.12e-11 *** |
| Threshold 0 | 3.9798 | 0.4660 | 3720 | -8.541 | < 2e-16 *** |
| Threshold 5 | 4.1390 | 0.4734 | 3709 | -8.743 | < 2e-16 *** |
| **Condition (YFP)** | -5.3655 | 1.4205 | 1010.8 | -3.777 | 0.0002 *** |
| Threshold - 10 * Condition (YFP) | 3.0573 | 1.5191 | 3725 | 2.013 | 0.0442 * |
| Threshold - 5 * Condition (YFP) | 3.9984 | 1.4063 | 3723 | 2.843 | 0.0045 ** |
| Threshold 0 * Condition (YFP) | 3.7923 | 1.3943 | 3726 | 2.720 | 0.0066 ** |
| Threshold 5 * Condition (YFP) | 4.1802 | 1.4024 | 3727 | 2.981 | 0.0029 ** |

**Pairwise Comparisons with Bootstrapped Confidence Intervals**

| **Contrast** | **Estimate** | **SE** | **df** | **t-ratio** | **p-value** | **Lower Bound (2.5%)** | **Upper Bound (97.5%)** |
| --- | --- | --- | --- | --- | --- | --- | --- |
| **Within ChR2 Condition** | | |  |  |  |  |  |
| -15 vs -10 | -0.9859 | 0.535 | 3727 | -1.842 | 0.3493 | 0.0031 | 2.0313 |
| -15 vs -5 | -3.3084 | 0.486 | 3716 | -6.803 | <.0001 | -3.4439 | -0.1934 |
| -15 vs 0 | -3.9798 | 0.466 | 3720 | -8.531 | <.0001 | -4.3831 | 2.9441 |
| -15 vs 5 | -4.1390 | 0.474 | 3709 | -8.731 | <.0001 | -5.6600 | 1.9381 |
| -10 vs -5 | -2.3225 | 0.363 | 3712 | -6.390 | <.0001 | -3.4439 | -0.1934 |
| -10 vs 0 | -2.9939 | 0.334 | 3727 | -8.962 | <.0001 | -4.3831 | 2.9441 |
| -10 vs 5 | -3.1531 | 0.343 | 3723 | -9.205 | <.0001 | -5.6600 | 1.9381 |
| -5 vs 0 | -0.6714 | 0.252 | 3435 | -2.666 | 0.0592 | -3.4439 | -0.1934 |
| -5 vs 5 | -0.8306 | 0.260 | 3519 | -3.190 | 0.0125 | -4.3831 | 2.9441 |
| 0 vs 5 | -0.1592 | 0.211 | 3727 | -0.755 | 0.9434 | -5.6600 | 1.9381 |
| **Within YFP Condition** | | | |  |  |  |  |
| -15 vs -10 | -2.0714 | 1.422 | 3724 | -1.457 | 0.5909 | -3.7307 | 3.6538 |
| -15 vs -5 | -0.6900 | 1.320 | 3720 | -0.523 | 0.9851 | -3.4439 | 2.9441 |
| -15 vs 0 | 0.1875 | 1.315 | 3725 | 0.143 | 0.9999 | -3.4439 | 2.9441 |
| -15 vs 5 | -0.0411 | 1.321 | 3726 | -0.031 | 1.0000 | -3.4439 | 2.9441 |
| -10 vs -5 | 1.3814 | 0.635 | 3726 | 2.174 | 0.1899 | -3.4439 | 2.9441 |
| -10 vs 0 | 2.2589 | 0.619 | 3706 | 3.647 | 0.0025 | -3.4439 | 2.9441 |
| -10 vs 5 | 2.0303 | 0.632 | 3685 | 3.210 | 0.0117 | -5.6600 | 1.9381 |
| -5 vs 0 | 0.8776 | 0.326 | 3587 | 2.693 | 0.0552 | -3.4439 | 2.9441 |
| -5 vs 5 | 0.6489 | 0.349 | 3526 | 1.862 | 0.3384 | -5.6600 | 1.9381 |
| 0 vs 5 | -0.2287 | 0.292 | 3723 | -0.782 | 0.9358 | -5.6600 | 1.9381 |
| **Between ChR2 and YFP** | | | |  |  |  |  |
| Threshold -15 | 5.3700 | 1.421 | 1000.5 | 3.776 | 0.0002 | 1.6925 | 9.1430 |
| Threshold -10 | 2.3100 | 0.754 | 101.9 | 3.062 | 0.0028 | 0.6605 | 3.9295 |
| Threshold -5 | 1.3700 | 0.489 | 18.4 | 2.797 | 0.0117 | 1.0611 | 2.0758 |
| Threshold 0 | 1.5700 | 0.439 | 12.0 | 3.585 | 0.0037 | 0.5844 | 1.7349 |
| Threshold 5 | 1.1900 | 0.459 | 14.4 | 2.582 | 0.0214 | 1.0611 | 2.0758 |

###### S2c: Beta Power (AUC)

The analysis examined the effects of laser threshold levels on beta power (AUC), with the condition (ChR2 and YFP) included as an additional predictor. The findings revealed significant effects of laser threshold levels on beta power (AUC) in the ChR2 condition. Higher thresholds consistently showed increased beta power (AUC) compared to lower thresholds. However, within the YFP condition, differences between thresholds were generally non-significant, with some exceptions.

Significant differences were also found between the ChR2 and YFP conditions across all laser thresholds, with ChR2 consistently showing higher beta power (AUC) than YFP. Given issues with heteroscedasticity, bootstrapping was performed to provide robust CIs for the estimates in the pairwise comparisons, reinforcing the reliability of the observed effects.

**Fixed Effects of Laser Thresholds and Condition on Beta Power (AUC)**

| **Predictor** | **Estimate** | **Std. Error** | **df** | **t-value** | **p-value** |
| --- | --- | --- | --- | --- | --- |
| **(Intercept)** | 2.8518 | 0.2787 | 423.16 | 10.233 | < 2e-16 *** |
| **Threshold - 15 (Reference)** | - | - | - | - | - |
| Threshold - 10 | 0.6970 | 0.3218 | 3687.84 | 2.166 | 0.03039 * |
| Threshold - 5 | 2.2435 | 0.2918 | 3572.54 | 7.687 | 1.93e-14 *** |
| Threshold 0 | 2.4832 | 0.2801 | 3584.55 | 8.867 | < 2e-16 *** |
| Threshold 5 | 2.5411 | 0.2842 | 3452.77 | 8.941 | < 2e-16 *** |
| **Condition (YFP)** | -2.1935 | 0.8359 | 2443.79 | -2.624 | 0.00874 ** |
| Threshold - 10 * Condition (YFP) | 1.0572 | 0.9153 | 3726.95 | 1.155 | 0.24812 |
| Threshold - 5 * Condition (YFP) | 1.5692 | 0.8475 | 3726.44 | 1.852 | 0.06418 . |
| Threshold 0 * Condition (YFP) | 1.3997 | 0.8397 | 3721.95 | 1.667 | 0.09564 . |
| Threshold 5 * Condition (YFP) | 1.3836 | 0.8443 | 3714.08 | 1.639 | 0.10137 |

**Pairwise Comparisons with Bootstrapped Confidence Intervals**

| **Contrast** | **Estimate** | **SE** | **df** | **t-ratio** | **p-value** | **Lower Bound (2.5%)** | **Upper Bound (97.5%)** |
| --- | --- | --- | --- | --- | --- | --- | --- |
| **Within ChR2 Condition** | | | | | |  |  |
| -15 vs -10 | -0.6970 | 0.322 | 3687 | 2.162 | 0.1946 | 0.1233 | 1.2453 |
| -15 vs -5 | -2.2435 | 0.293 | 3568 | 7.664 | <.0001 | -2.2024 | 1.7329 |
| -15 vs 0 | -2.4832 | 0.281 | 3581 | 8.839 | <.0001 | -2.3669 | 2.8827 |
| -15 vs 5 | -2.5411 | 0.285 | 3446 | 8.903 | <.0001 | -2.2024 | 2.8827 |
| -10 vs -5 | -1.5464 | 0.219 | 3585 | 7.072 | <.0001 | -2.2024 | 1.7329 |
| -10 vs 0 | -1.7862 | 0.201 | 3705 | 8.875 | <.0001 | -2.3669 | 2.8827 |
| -10 vs 5 | -1.8441 | 0.206 | 3627 | 8.938 | <.0001 | -2.2024 | 2.8827 |
| -5 vs 0 | -0.2398 | 0.151 | 2478 | 1.590 | 0.5041 | -0.4259 | 2.9185 |
| -5 vs 5 | -0.2977 | 0.156 | 2634 | 1.906 | 0.3142 | -0.4259 | 2.9185 |
| 0 vs 5 | -0.0579 | 0.127 | 3695 | 0.456 | 0.9911 | -0.4259 | 2.9185 |
| **Within YFP Condition** | | | |  |  |  |  |
| -15 vs -10 | -0.3602 | 0.857 | 3727 | -0.420 | 0.9935 | -0.4260 | 2.9185 |
| -15 vs -5 | 0.6742 | 0.796 | 3720 | 0.847 | 0.9157 | -0.4723 | 2.8827 |
| -15 vs 0 | 1.0836 | 0.792 | 3726 | 1.368 | 0.6485 | -0.4723 | 2.8827 |
| -15 vs 5 | 1.1576 | 0.796 | 3723 | 1.455 | 0.5922 | -0.4723 | 2.8827 |
| -10 vs -5 | 1.0344 | 0.383 | 3699 | 2.704 | 0.0536 | -0.4723 | 2.8827 |
| -10 vs 0 | 1.4438 | 0.373 | 3546 | 3.875 | 0.0010 | -0.4723 | 2.8827 |
| -10 vs 5 | 1.5177 | 0.380 | 3428 | 3.992 | 0.0006 | -0.4723 | 2.8827 |
| -5 vs 0 | 0.4093 | 0.196 | 2828 | 2.091 | 0.2239 | -0.4723 | 2.8827 |
| -5 vs 5 | 0.4833 | 0.209 | 2585 | 2.311 | 0.1416 | -0.4723 | 2.8827 |
| 0 vs 5 | 0.0740 | 0.176 | 3726 | 0.420 | 0.9935 | -0.4723 | 2.8827 |
| **Between ChR2 and YFP** | | |  |  |  |  |  |
| Threshold -15 | 2.193 | 0.837 | 2420.5 | 2.622 | 0.0088 | 0.4453 | 3.8712 |
| Threshold -10 | 1.136 | 0.415 | 367.7 | 2.736 | 0.0065 | 0.0854 | 2.1957 |
| Threshold -5 | 0.624 | 0.230 | 39.3 | 2.709 | 0.0099 | 0.2607 | 1.1213 |
| Threshold 0 | 0.794 | 0.191 | 18.7 | 4.167 | 0.0005 | 0.4214 | 1.1761 |
| Threshold 5 | 0.810 | 0.207 | 25.6 | 3.913 | 0.0006 | 0.5129 | 1.1761 |

###### S2c: Gamma Power (AUC)

The analysis explored the effects of laser threshold levels on gamma power (AUC), considering the condition (ChR2 and YFP) as an additional predictor. The results indicated that the laser threshold levels had a significant impact on gamma power (AUC) in the ChR2 condition, with lower thresholds generally leading to lower gamma power (AUC). Within the YFP condition, differences between thresholds were mostly non-significant, although some comparisons did show effects.

The comparison between ChR2 and YFP conditions revealed significant differences at thresholds -15 and -10, with ChR2 showing consistently lower gamma power (AUC) compared to YFP at these levels. Due to slight issues with normality, bootstrapping was employed to provide robust CIs for the estimates, ensuring the reliability of the observed effects.

**Fixed Effects of Laser Thresholds and Condition on Gamma Power (AUC)**

| **Predictor** | **Estimate** | **Std. Error** | **df** | **t-value** | **p-value** |
| --- | --- | --- | --- | --- | --- |
| **(Intercept)** | -0.7268 | 0.2008 | 101.44 | -3.619 | 0.00046 *** |
| **Threshold - 15 (Reference)** | - | - | - | - | - |
| Threshold - 10 | 0.7988 | 0.2101 | 3727.00 | 3.801 | 0.00015 *** |
| Threshold - 5 | 0.7077 | 0.1909 | 3720.33 | 3.708 | 0.00021 *** |
| Threshold 0 | 0.7428 | 0.1831 | 3723.35 | 4.057 | 5.08e-05 *** |
| Threshold 5 | 0.7198 | 0.1860 | 3715.82 | 3.869 | 0.00011 *** |
| **Condition (YFP)** | 1.3230 | 0.5612 | 823.06 | 2.358 | 0.01862 * |
| Threshold - 10 * Condition (YFP) | -0.4726 | 0.5968 | 3724.18 | -0.792 | 0.42848 |
| Threshold - 5 * Condition (YFP) | -0.9157 | 0.5525 | 3722.47 | -1.657 | 0.09751 . |
| Threshold 0 * Condition (YFP) | -1.2928 | 0.5478 | 3725.79 | -2.360 | 0.01833 * |
| Threshold 5 * Condition (YFP) | -1.7129 | 0.5510 | 3726.58 | -3.109 | 0.00189 ** |

**Pairwise Comparisons with Bootstrapped Confidence Intervals**

| **Contrast** | **Estimate** | **SE** | **df** | **t-ratio** | **p-value** | **Lower Bound (2.5%)** | **Upper Bound (97.5%)** |
| --- | --- | --- | --- | --- | --- | --- | --- |
| **Within ChR2 Condition** | | | |  |  |  |  |
| -15 vs -10 | -0.7988 | 0.2102 | 3727 | -3.799 | 0.0014 | -1.2243 | -0.3941 |
| -15 vs -5 | -0.7077 | 0.1911 | 3720 | -3.704 | 0.0020 | -1.0868 | -0.3522 |
| -15 vs 0 | -0.7428 | 0.1833 | 3723 | -4.053 | 0.0005 | -1.1176 | -0.3984 |
| -15 vs 5 | -0.7198 | 0.1863 | 3716 | -3.864 | 0.0011 | -1.0908 | -0.3554 |
| -10 vs -5 | 0.0910 | 0.1428 | 3718 | 0.637 | 0.9689 | -0.2134 | 0.3427 |
| -10 vs 0 | 0.0560 | 0.1312 | 3727 | 0.426 | 0.9931 | -0.1972 | 0.3628 |
| -10 vs 5 | 0.0790 | 0.1346 | 3725 | 0.587 | 0.9770 | -0.2082 | 0.1416 |
| -5 vs 0 | -0.0351 | 0.0990 | 3506 | -0.354 | 0.9966 | -0.1803 | 0.1636 |
| -5 vs 5 | -0.0120 | 0.1023 | 3573 | -0.118 | 1.0000 | -0.1172 | 0.1821 |
| 0 vs 5 | 0.0230 | 0.0829 | 3727 | 0.278 | 0.9987 | -0.1172 | 0.1821 |
| **Within YFP Condition** | | | |  |  |  |  |
| -15 vs -10 | -0.3262 | 0.5587 | 3723 | -0.584 | 0.9775 | -2.1862 | -0.3985 |
| -15 vs -5 | 0.2080 | 0.5185 | 3719 | 0.401 | 0.9946 | -1.4832 | -0.2408 |
| -15 vs 0 | 0.5500 | 0.5164 | 3725 | 1.065 | 0.8245 | -0.6717 | -0.1711 |
| -15 vs 5 | 0.9931 | 0.5188 | 3726 | 1.914 | 0.3098 | -0.7232 | -0.3423 |
| -10 vs -5 | 0.5342 | 0.2497 | 3727 | 2.139 | 0.2036 | -0.6717 | -0.1711 |
| -10 vs 0 | 0.8761 | 0.2434 | 3713 | 3.600 | 0.0030 | -0.6717 | -0.1711 |
| -10 vs 5 | 1.3193 | 0.2485 | 3698 | 5.309 | <.0001 | -0.6717 | -0.1711 |
| -5 vs 0 | 0.3420 | 0.1281 | 3626 | 2.670 | 0.0587 | -0.6717 | -0.1711 |
| -5 vs 5 | 0.7851 | 0.1370 | 3580 | 5.732 | <.0001 | -0.6717 | -0.1711 |
| 0 vs 5 | 0.4431 | 0.1148 | 3722 | 3.858 | 0.0011 | -0.6717 | -0.1711 |
| **Between ChR2 and YFP** | |  |  |  |  |  |  |
| Threshold -15 | -1.3230 | 0.561 | 831.2 | -2.357 | 0.0187 | -2.1862 | -0.3985 |
| Threshold -10 | -0.8505 | 0.302 | 84.9 | -2.817 | 0.0060 | -1.4832 | -0.2408 |
| Threshold -5 | -0.4073 | 0.201 | 16.9 | -2.028 | 0.0585 | -1.4832 | -0.2408 |
| Threshold 0 | -0.0303 | 0.182 | 11.5 | -0.166 | 0.8709 | -1.4832 | -0.2408 |
| Threshold 5 | 0.3898 | 0.190 | 13.5 | 2.055 | 0.0598 | -1.4832 | -0.2408 |

##### Supplementary figure 3: YFP for Figure 2

###### S3f: NE (AUC) & RR (AUC): pre-stimulation minus post stimulation correlation (all thresholds)

The correlation between RR and NE showed a slight negative relationship. Due to slight issues with normality of residuals, the marginal and conditional R squared estimates were bootstrapped, which supported the primary findings.

| **Statistic** | **Value** |
| --- | --- |
| **Fixed Effects Estimates** |  |
| Intercept Estimate | 7.02 |
| Intercept SE | 5.44 |
| Intercept df | 2.70 |
| Intercept t-value | 1.29 |
| Intercept p-value | 0.30 |
| NE Estimate | -0.91 |
| NE SE | 0.14 |
| NE df | 1244.68 |
| NE t-value | -6.75 |
| NE p-value | <0.0001*** |
| **Random Effects** |  |
| SubjectID (Intercept) Variance | 37.76 |
| SubjectID (Intercept) SD | 6.15 |
| Residual Variance | 24372.26 |
| Residual SD | 156.12 |
| **R-squared Values** |  |
| Marginal R-squared (R²m) | 0.04 |
| Bootstrapped 2.5% CI | 0.02 |
| Bootstrapped 97.5% CI | 0.06 |
| Conditional R-squared (R²c) | 0.04 |
| Bootstrapped 2.5% CI | 0.02 |
| Bootstrapped 97.5% CI | 0.06 |
| **Correlation of Fixed Effects** |  |
| Correlation (Intercept, NE) | 0.08 |

###### S3f: NE (AUC) & RR (AUC): pre-stimulation minus post stimulation correlation (Threshold -15 to -5)

The correlation between RR and NE showed a negative relationship. Due to slight issues with normality of residuals, the marginal and conditional R squared estimates were bootstrapped, which supported the primary findings.

| **Statistic** | **Value** |
| --- | --- |
| **Fixed Effects Estimates** |  |
| Intercept Estimate | 0.92 |
| Intercept SE | 21.73 |
| Intercept df | 2.45 |
| Intercept t-value | 0.04 |
| Intercept p-value | 0.97 |
| NE Estimate | -1.08 |
| NE SE | 0.27 |
| NE df | 356.94 |
| NE t-value | -4.03 |
| NE p-value | <0.0001*** |
| **Random Effects** |  |
| SubjectID (Intercept) Variance | 1539.00 |
| SubjectID (Intercept) SD | 39.23 |
| Residual Variance | 21918.00 |
| Residual SD | 148.05 |
| **R-squared Values** |  |
| Marginal R-squared (R²m) | 0.04 |
| Bootstrapped 2.5% CI | 0.01 |
| Bootstrapped 97.5% CI | 0.09 |
| Conditional R-squared (R²c) | 0.10 |
| Bootstrapped 2.5% CI | 0.04 |
| Bootstrapped 97.5% CI | 0.25 |
| **Correlation of Fixed Effects** |  |
| Correlation (Intercept, NE) | 0.14 |

**S3h: Correlation comparison between ChR2 and YFP**

To test whether the coupling between the NE responce and RR differed between ChR2- and YFP-expressing animals, we fit linear mixed-effects models with RR as the response and the NE response, group (ChR2 vs. YFP, YFP as reference), and their interaction as fixed effects. Each NREM bout was treated as an observation, and a by-animal random intercept accounted for the non-independence of bouts recorded from the same mouse. The Group× NE interaction term tested whether the NE–RR slope differed between groups. As a descriptive check, we also compared Pearson correlations using Fisher r-to-z transformation, noting that this approach does not account for repeated measures structure.

**Linear mixed-effects models of the NE–RR relationship**

|  | **NE→RR slope, YFP** | **NE→RR slope, ChR2** | **Slope difference** | **R²ₘ** | **R²c** |
| --- | --- | --- | --- | --- | --- |
| **Including all threshold** | **−0.911 [−1.18, -0.65]** | **−2.826 [−3.15, −2.50]** | **1.915, p < 0.0001** | **0.091** | **0.093** |
| **Including threshold -15, -10 and -5** | **−1.095 [−1.63, −0.56]** | **−3.462 [−3.99, −2.94]** | **2.367, p < 0.0001** | **0.164** | **0.178** |

**Fisher r-to-z transformation**

|  | **r_ChR2** | **n_ChR2** | **r_YFP** | **n_YFP** | **Z** | **P_value** |
| --- | --- | --- | --- | --- | --- | --- |
| **Including all threshold** | **-0.325** | **2488** | **-0.187** | **1249** | **-4.253** | **2.11e-05** |
| **Including threshold -15, -10 and -5** | **-0.419** | **783** | **-0.419** | **362** | **-3.651** | **2.62e-04** |
